# A multi-stakeholder Delphi consensus study proposes actionable solutions to address antibody validation failures

**DOI:** 10.64898/2026.03.04.709541

**Authors:** Katherine Blades, Michael Biddle, Robert Froud, Eva M. Krockow, Harvinder Virk

**Author notes:** joint corresponding authors: Harvinder Virk –, Eva Krockow –.

## Abstract

The experimental use of antibodies that have not been validated for context-specific use frequently misdirects biomedical research. Experimental results that derive from the use of inadequately validated antibodies are estimated to waste over $1 billion annually in the United States alone and to consume millions of animal and human biological samples in experiments whose conclusions may be unreliable. Community validation frameworks, reporting standards, and independent characterisation initiatives have made important progress, and multi-stakeholder coordination efforts are emerging. However, the research community lacks a formally developed, consensus-based action plan that specifies what each stakeholder group should do, by when, and with what priority. We conducted a modified Delphi study with international experts representing academic researchers, scientific publishers, research funders, antibody manufacturers, and institutional research leaders to develop actionable recommendations for improving antibody validation, selection, and reporting practices. Thirty-two participants rated 33 proposed actions on effectiveness and feasibility using 9-point scales, with consensus assessed using the RAND/UCLA Appropriateness Method. Over two rounds, the panel achieved consensus on 15 items as both effective and feasible for implementation by 2030. These spanned institutional actions (training in antibody validation, integration into research integrity frameworks, support for local expertise networks), funder actions (dedicated validation budgets, grant application requirements, endorsement of community reporting standards), publisher actions (complete antibody reporting packages, clear validation standards), manufacturer actions (assignment of unique identifiers at source), and cross-stakeholder coordination (a shared roadmap for improvement). An additional 15 items were rated as effective but with uncertain feasibility, reflecting a consistent pattern in which the panel agreed on the value of proposed interventions but expressed reservations about realistic implementation timelines. One item was rejected by the panel with concerns around effectiveness and feasibility. Participants described four interconnected barriers to progress: diffuse ownership of the problem across stakeholders; market dynamics that inadequately reward antibody quality; difficulty justifying investment when returns are distributed across the research system; and coordination challenges among actors with different incentive structures. These barriers are addressable through coordinated action, and the findings complement existing technical and data-sharing initiatives by providing the structured, expert-informed policy framework to help translate awareness of the problem into concrete practice and policy changes.

## Introduction

Antibodies are among the most widely used tools in biomedical research, enabling researchers to detect, quantify, and isolate specific proteins within complex biological samples. However, research antibodies do not always bind exclusively to their intended targets. Poor antibody specificity has repeatedly misdirected biomedical research across diverse fields, leading to incorrect conclusions about protein function, unreliable biomarker discovery, and misidentification of drug targets [1–10]. These failures represent not isolated occurrences but a persistent and systemic issue affecting the reliability of the published literature.

The scale of this problem is substantial. An estimated $28.2 billion per year is spent in the United States on preclinical research that is not reproducible, with issues relating to biological reagents likely being the biggest contributor [11]. Antibody-related failures alone account for an estimated $1 billion annually in the US in direct economic waste [12, 13], a figure that excludes the substantial opportunity cost of misdirected research effort. Beyond the economic dimension, there is a large and largely unquantified ethical cost: the waste of animals both in antibody production and in research studies that employ unsuitable antibodies, and the waste of donated human biological materials in experiments whose conclusions may be unreliable [14]. Recent estimates suggest that at least 4 million animal samples and 6 million human tissue samples have been consumed globally in experiments using antibodies that failed independent testing, without context-specific validation since December, 2000 [14].

How well an antibody performs in binding to its target varies between applications, protocols, and the cell or tissue types used for each experiment. Consequently, it is important that performance is assessed for each context of use, a process referred to as validation, and it is the responsibility of the researcher to carry out these studies for their specific protocol [15]. The complexity and scale of antibody use have contributed to a marketplace containing a high proportion of poorly performing antibodies [1]. Recent independent benchmarking by the YCharOS (Antibody Characterisation by Open Science) consortium quantified the likely scale of the problem: greater than 50% of 614 commercial antibodies against 65 neuroscience-related proteins failed characterisation experiments in three commonly used applications, and 88.4% of papers using poorly performing antibodies in immunofluorescence presented no relevant validation data [1]. Moreover, each protein target was linked to an average of approximately 12 published papers that presented data using poorly performing antibodies, perpetuating the use of these reagents and the associated issues of reproducibility.

Efforts to address this problem have operated on multiple fronts. Technical advances include the IWGAV five-pillar validation framework, which outlines complementary strategies (genetic, orthogonal, independent antibody, tagged protein, and immunocapture mass spectrometry), for establishing antibody selectivity [15], and the large-scale independent characterisation work of the YCharOS consortium, which has provided open, head-to-head comparisons of commercial antibodies using knockout cell lines as isogenic controls [1]. Reporting standards such as the Materials Design Analysis Reporting (MDAR) framework have sought to improve transparency at the point of publication [16], and tools including the Research Resource Identifier (RRID) initiative have enabled more precise tracking of antibody use across the literature [17]. Individual manufacturers have also invested in more rigorous characterisation, in some cases removing or relabelling products found to perform poorly [1, 14].

In parallel, multi-stakeholder coordination has begun to emerge. Kahn et al. [12] assembled a history of efforts to address the antibody characterisation crisis and proposed sector-specific actions for researchers, universities, journals, vendors, scientific societies, disease foundations, and funding agencies. The Only Good Antibodies (OGA) community, of which members of the present study team are founding contributors [18], has brought together stakeholders across the research ecosystem to work toward making best practice in antibody use more feasible, easy, and rewarding, drawing on the Centre for Open Science’s framework for culture change in research. These initiatives represent important progress. However, the field still lacks a formally developed, consensus-based action plan in which proposed interventions have been systematically rated for both effectiveness and feasibility by a diverse expert panel. Without such structured prioritisation, it is difficult for stakeholders to know which actions to implement first, where the strongest agreement exists, and where feasibility barriers require targeted attention.

The underlying reasons for the persistence of this problem are complex and multifaceted, encompassing the inherent challenges of producing and characterising high-quality antibodies, but also researcher awareness, education, and incentive structures [14]. Researchers identify time, cost, and lack of supervisor support as primary barriers to better validation practices, while strongly supporting structural solutions including open data sharing, dedicated validation funding, and publisher requirements [14]. Behavioural analysis drawing on the COM-B (Capabilities + Opportunities + Motivation Behaviour) model suggests that individual capability and motivation interact with the opportunity provided by the research environment, such that improvements to practice require changes not only in individual behaviour but in the systems and structures that shape it [14, 18]. This suggests that the problem requires coordinated action across the research ecosystem.

Having established the scale and drivers of the problem [14], and situated it within the broader landscape of ongoing technical and community efforts [1, 12, 18], this study sought to move from identifying what needs to change, to building structured agreement on how to change it. We convened a Delphi study to develop recommendations for improving antibody validation, selection, and reporting practices, and to analyse the obstacles to their implementation.

## Methods

### Study Design and Ethical Approval

We conducted a modified Delphi study to develop expert-informed recommendations for improving antibody validation, selection, and reporting practices in biomedical research. The Delphi method is a structured consensus-building approach that systematically collects and synthesises expert opinion through iterative questionnaires with controlled feedback [19,20]. This study was approved by the University of Leicester College of Life Sciences Research Ethics Committee (reference number [2025-1122-5743]) and conducted in accordance with the General Data Protection Regulation (GDPR, 2018). All participants provided written informed consent before participation. Data collection and management were conducted using the Clinvivo platform (https://www.clinvivo.com; Clinvivo Ltd, London, UK), a secure GDPR-compliant system designed specifically for Delphi studies.

### Participant Recruitment

We identified experts from five key sectors involved in antibody use and research governance: academic biomedical research, scientific publishing, research funding, antibody manufacturing, and institutional research leadership. Experts were defined as individuals who are representative of their profession, have the power to implement findings, or who would not be challenged as experts in their field [21]. Thirty-eight experts were identified through networks based on their expertise in antibody validation, editorial roles at major biomedical journals, leadership positions at research funding organisations, executive roles in commercial antibody production, or responsibility for research integrity policies at academic institutions. Tailored recruitment emails were sent to each stakeholder group. Participants were informed that the study aimed to develop a consensus-based action plan to improve antibody validation and reporting across the research ecosystem, and that their responses would be analysed together as a single, non-identifiable expert panel to inform policy recommendations for journals, funders, institutions, and manufacturers.

To assemble a panel spanning these sectors, we drew on the leadership and networks of established expert consortia, each asked to identify appropriate contributors from their community: the Antibody Society (representing the antibody industry and its user community), YCharOS (an independent antibody-validation consortium), the NC3Rs (a research funder with an established antibody-reproducibility programme), and the UK Reproducibility Network (providing access to institutional research leadership). A senior contact within each body was tasked with nominating relevant experts, who were then invited by the study team; these same convening bodies had contributed to an earlier multi-stakeholder meeting on antibody reproducibility [22]. Recruitment through recognised convening bodies has limitations, in that it reflects the membership and reach of those networks and may under-represent experts outside them; however, in the absence of any defined sampling frame of antibody experts, it offers a transparent and practical means of assembling a credible cross-stakeholder panel.

### The Delphi Process

The Delphi process consisted of iterative rounds of anonymous questionnaire completion with structured feedback. Initial draft statements for Round 1 were developed from two sources: recommendations emerging from a stakeholder meeting convened by the National Centre for the Replacement, Refinement and Reduction of Animals in Research (NC3Rs) in February 2024 [22], and themes identified through focus group discussions (and validated with a survey) with antibody-using researchers conducted in the summer of 2023 [14]. These sources yielded 23 proposed actions spanning four domains: journal and publisher policies (n=9 items covering reporting requirements, validation transparency, and editorial capacity building), research funder actions (n=9 items covering grant requirements, validation funding, and manufacturer engagement), institutional and educational interventions (n= 3 items covering training, ethics frameworks, and local champions), and cross-stakeholder coordination mechanisms (n= 2 items covering roadmaps, shared infrastructure, and benchmarking initiatives). Each item described a specific action that one or more stakeholder groups could adopt to improve antibody practices (S6 Text). In Round 1, participants were given the opportunity to nominate additional items not already covered in Round 1 for inclusion in Round 2.

Round 1 was conducted from August 12 to September 12, 2025, with two deadline extensions to maximise response rates. Participants received weekly email reminders from Clinvivo, and the research team sent a final reminder on the originally scheduled closing date. Round 2 was conducted from November 6 to November 23, 2025, following the same reminder protocol. The decision to proceed to additional rounds was based on pre-specified criteria: lack of consensus, persistence of significant disagreement requiring resolution, or evidence that additional rounds would meaningfully shift ratings (e.g., based on qualitative feedback).

### Rating System and Assessment of Consensus

Participants rated each proposed action on two 9-point rating scales. The effectiveness scale permitted panellists to rate how meaningfully they thought the item would improve the selection, validation, or reporting of antibodies in research (1=extremely ineffective, 9=extremely effective). The feasibility scale permitted panellists to rate how realistically they thought items could be adopted across relevant settings by 2030 (1=extremely infeasible, 9=extremely feasible). Participants could select “don’t know” if an item fell outside their area of expertise. “Don’t know” responses were treated as non-responses and excluded from the calculations; medians, the IPR and the IPRAS for each item were therefore based only on the experts who provided a numeric rating for that item, so the number of ratings contributing to these statistics varied across items. For each item, participants could provide free-text comments to share their rationale, identify potential risks or unintended consequences, or suggest alternative wordings.

To assess disagreement, we employed the RAND/UCLA Appropriateness Method [19]. For each item, we calculated the median rating, the inter-percentile range (IPR; difference between the 30th and 70th percentiles), and the inter-percentile range adjusted for symmetry (IPRAS). The IPRAS accounts for asymmetric rating distributions using the equation: IPRAS = IPRr + (AI × CFA), where IPRr is the inter-percentile range required for disagreement when perfect symmetry exists (2.35), AI is the asymmetry index (calculated as the distance between the central point of the IPR and the scale midpoint of 5), and CFA is the correction factor for asymmetry (1.5). These values of 2.35 and 1.5 were empirically derived by Fitch et al. [19] to best reproduce classic RAND definitions of disagreement for 9-point scales. Percentiles were calculated using Stata version 18 (StataCorp LLC, College Station, TX; RRID:SCR_012763) with the R-2 formula (Stata’s default method).

An item was classified as having disagreement when IPR > IPRAS for that item. Items were classified according to median ratings: effective/feasible (median 7-9), ineffective/infeasible (median 1-3), or uncertain (median 4-6). For medians falling on category boundaries (3.5 or 6.5) due to even participant numbers, we applied a “round down” rule, classifying 3.5 as ineffective/infeasible and 6.5 as uncertain. This meant a conservative approach was taken to describing ratings as effective or feasible. Because this rule resolves boundary medians into the lower band, it can only have reduced the number of items meeting the effective-and-feasible criterion, never increased it; the main pattern of results is therefore not dependent on a less conservative rounding choice. Qualitative free-text comments were reviewed by the research team to inform item refinement, identify new proposed actions, and develop narrative summaries for feedback to participants in subsequent rounds.

### Progression Decisions Between Rounds

Items were progressed, removed, or modified between rounds based on their ratings, the presence or absence of disagreement, or qualitative feedback. Items rated as both effective (median ≥7) and feasible (median ≥7) without disagreement were accepted as recommendations and did not require re-rating in subsequent rounds unless there were suggestions for rewording in the qualitative comments. Items rated as both ineffective (median ≤3) and infeasible (median ≤3) without disagreement were rejected and removed from further consideration. Items classified as uncertain on either dimension, or those with disagreement, were considered for re-rating in the next round, particularly if qualitative comments suggested that rewording, clarification, or presentation of divergent perspectives might shift opinions. Between Round 1 and Round 2, panel members’ free-text comments were reviewed to identify: (1) suggestions for new action items to add to the consensus process, (2) proposed modifications to existing item wording, and (3) substantive concerns or alternative perspectives that should be shared with the panel. Items were reworded when comments indicated ambiguity or suggested semantically meaningful improvements. New items suggested by multiple participants or representing distinct actionable proposals were added to Round 2. For items with disagreement or uncertainty, we prepared feedback summaries that synthesised the range of panel perspectives to support informed re-rating. Items with substantive wording changes based on panel feedback were re-rated regardless of initial ratings, to ensure consensus reflected the revised formulation. The number of rounds was therefore not fixed in advance but determined by these rules: rounds were planned to continue while items remained in disagreement or uncertainty and movement under directed feedback remained plausible, and the process concluded once the rating-relevant items had settled or stabilised. Consensus on every item was not the objective; a range of outcomes, with items classified as effective and feasible, effective but of uncertain feasibility, or rejected, was the intended result.

## Results

### Participant Characteristics and Response Rates

Thirty-eight individuals were invited to participate in Round 1. Thirty-two completed the round (response rate: 84.2%). All Round 1 participants were invited to Round 2, and 30 completed this round (response rate: 93.8%; overall retention: 78.9% of original invitees).

The 32 Round 1 participants included 13 females (40.6%), 17 males (53.1%), and 2 who preferred not to disclose sex (6.3%). Among the 30 participants who reported their age, the mean was 51.5 years (SD=11.3). Participants were primarily based in the United Kingdom (n=16, 50.0%) and United States (n=13, 40.6%), with additional representation from Canada (n=2, 6.3%) and Germany (n=1, 3.1%). The panel was highly educated, with 27 participants (84.4%) holding PhD-level qualifications, 2 (6.3%) holding master’s degrees, and 3 (9.4%) holding bachelor’s degrees. Participants represented diverse fields within the biomedical research ecosystem: academic biomedical research (n=7, 21.9%), scientific publishing (n=6, 18.8%), antibody manufacturing (n=6, 18.8%), research funding (n=4, 12.5%), industrial biomedical research (n=2, 6.3%), and other fields (n=7, 21.9%) including antibody informatics, infrastructure provision, and the charity sector. Within these sectors, participants held senior decision-making roles: 37.5% (n=12) reported full autonomy to make decisions on behalf of their organisations, 34.4% (n=11) made decisions with input from senior leaders, and 28.1% (n=9) made recommendations with final decisions made by senior leaders. Detailed breakdowns of career stages within each employment sector are provided in S1 Table.

### Overview of Items and Consensus Development

A total of 33 proposed action items were rated across two rounds (Figure 1). “Don’t know” responses were uncommon. Per item they ranged from 0 to 6 of 32 (effectiveness) and 0 to 7 of 32 (feasibility) in Round 1, and from 0 to 4 of 30 and 0 to 8 of 30 in Round 2, with most items rated by the whole panel or nearly so; the few higher counts fell on items directed at a specific sector, where panellists outside that sector could appropriately abstain. Per-item response rates for every item are reported in S2 and S3 Tables.

**Figure 1.**
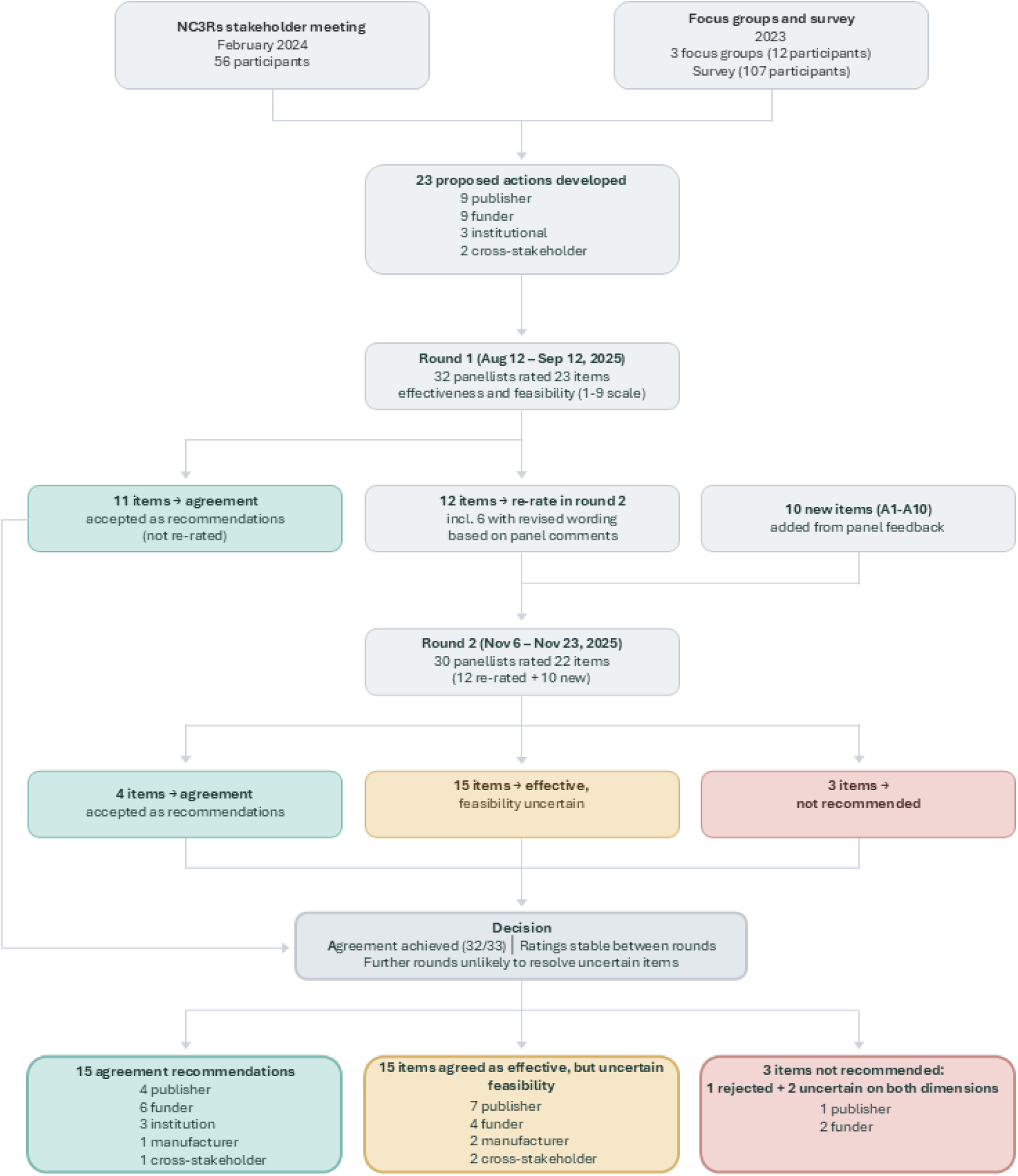
Overview of the Delphi process including formulation of recommendations and their assessment across rounds.

Round 1 presented 23 items spanning four domains: journal and publisher policies (9 items, R1– R9), research funder actions (9 items, R10–R18), institutional and educational interventions (3 items, R19–R21), and cross-stakeholder coordination (2 items, R22–R23), see S6 Text for full item wording. All 32 participants rated effectiveness and feasibility for each of the 23 proposed actions. Panel ratings demonstrated strong consensus on most items, with disagreement identified on only 2 of 46 ratings (effectiveness and feasibility combined).

Following Round 1 analysis, 11 items (R1, R2, R10–R15, R19, R20, R22) achieved consensus as both effective (median ≥7) and feasible (median ≥7) without disagreement and were immediately accepted as recommendations without requiring re-rating. The remaining 12 items from Round 1 were re-presented in Round 2, including six with revised wording based on panel comments (R3, R8, R9, R17, R18, R21). Based on participant free-text suggestions, 10 additional action items (A1– A10) were added to Round 2. Round 2 thus presented 22 items for rating: 12 re-rated items from Round 1 and 10 newly proposed items.

In Round 2 disagreement only occurred on 1 of 44 ratings. Among the 22 items presented in Round 2, four achieved agreement as both effective and feasible (R3, R21, A1, A7), joining the 11 items accepted in Round 1 for a total of 15 consensus recommendations at the close of Round 2. One item (R5) shifted from uncertain in Round 1 to consensually infeasible in Round 2 (median feasibility declined from 4.0 to 3.0) and was the only item to be rated either infeasible or ineffective. Two items (A4, A5) were rated as uncertain on both effectiveness and feasibility.

### Final Consensus Outcomes

Across the two rounds, the panel evaluated 33 proposed actions, achieving consensus on 15 items as both effective (median ≥7) and feasible (median ≥7), without disagreement (Table 1). These 15 recommendations span all four stakeholder domains and represent actions that expert panellists agreed would meaningfully improve antibody practices and could realistically be implemented by 2030.

**TABLE 1:**
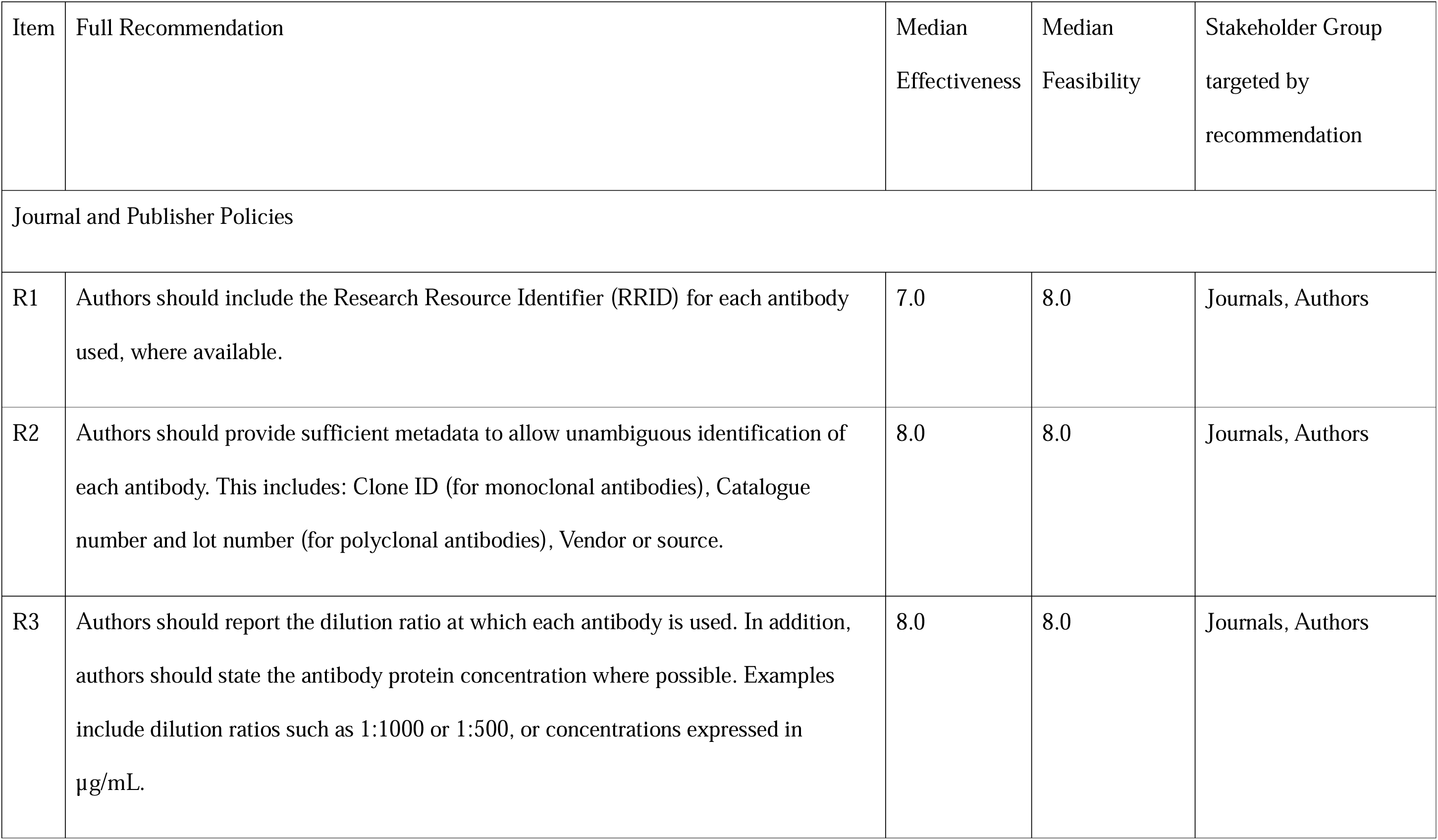
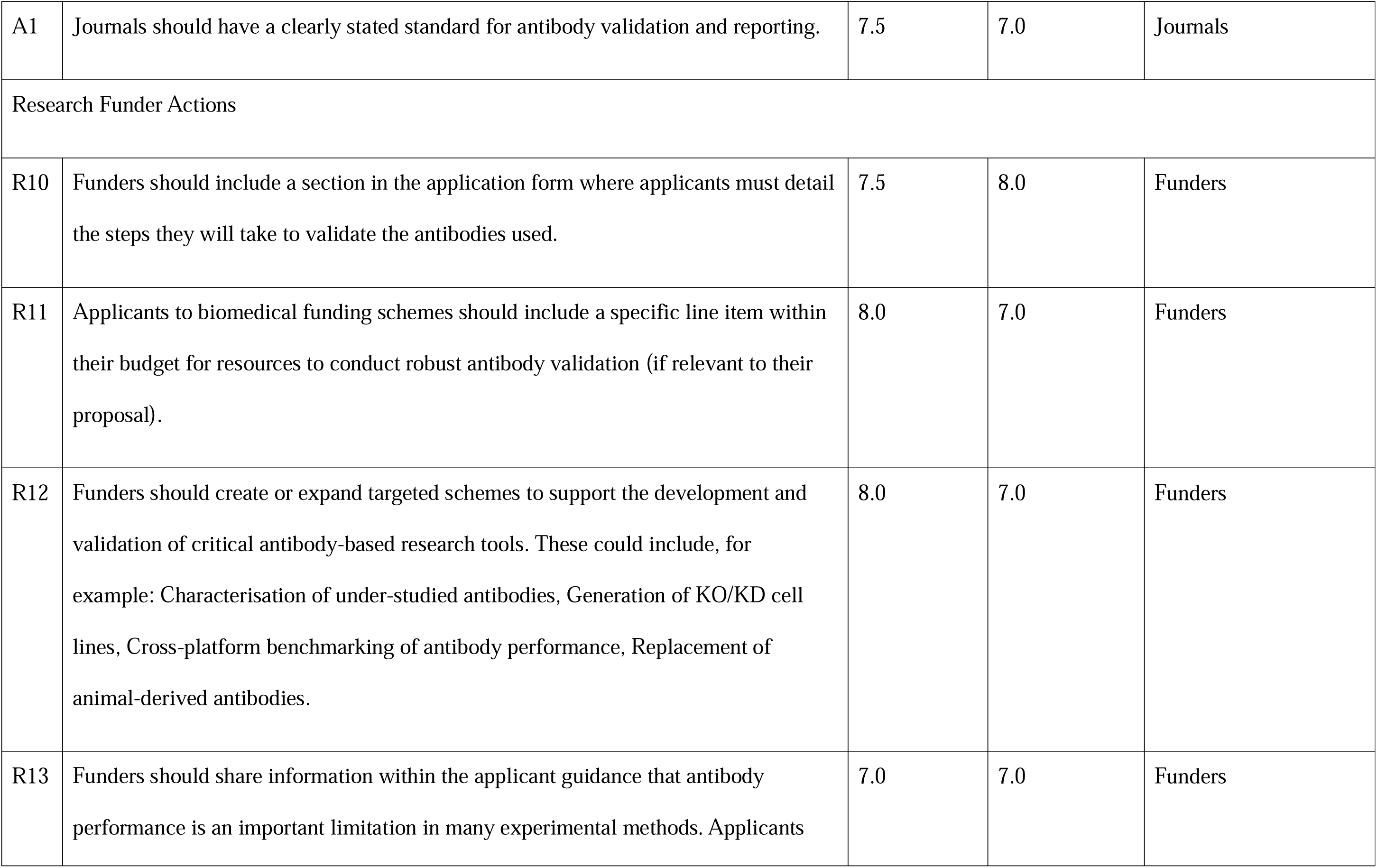
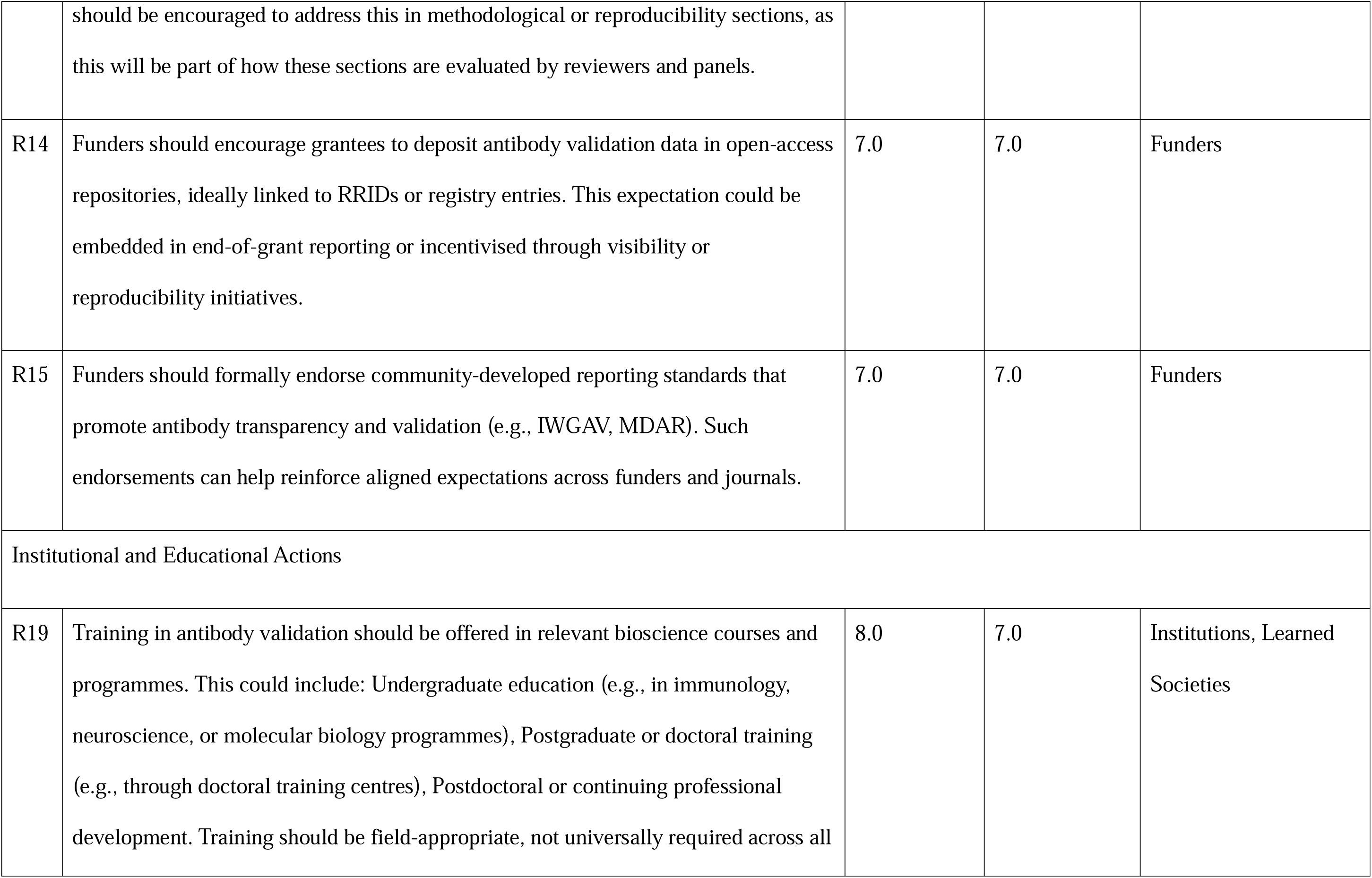
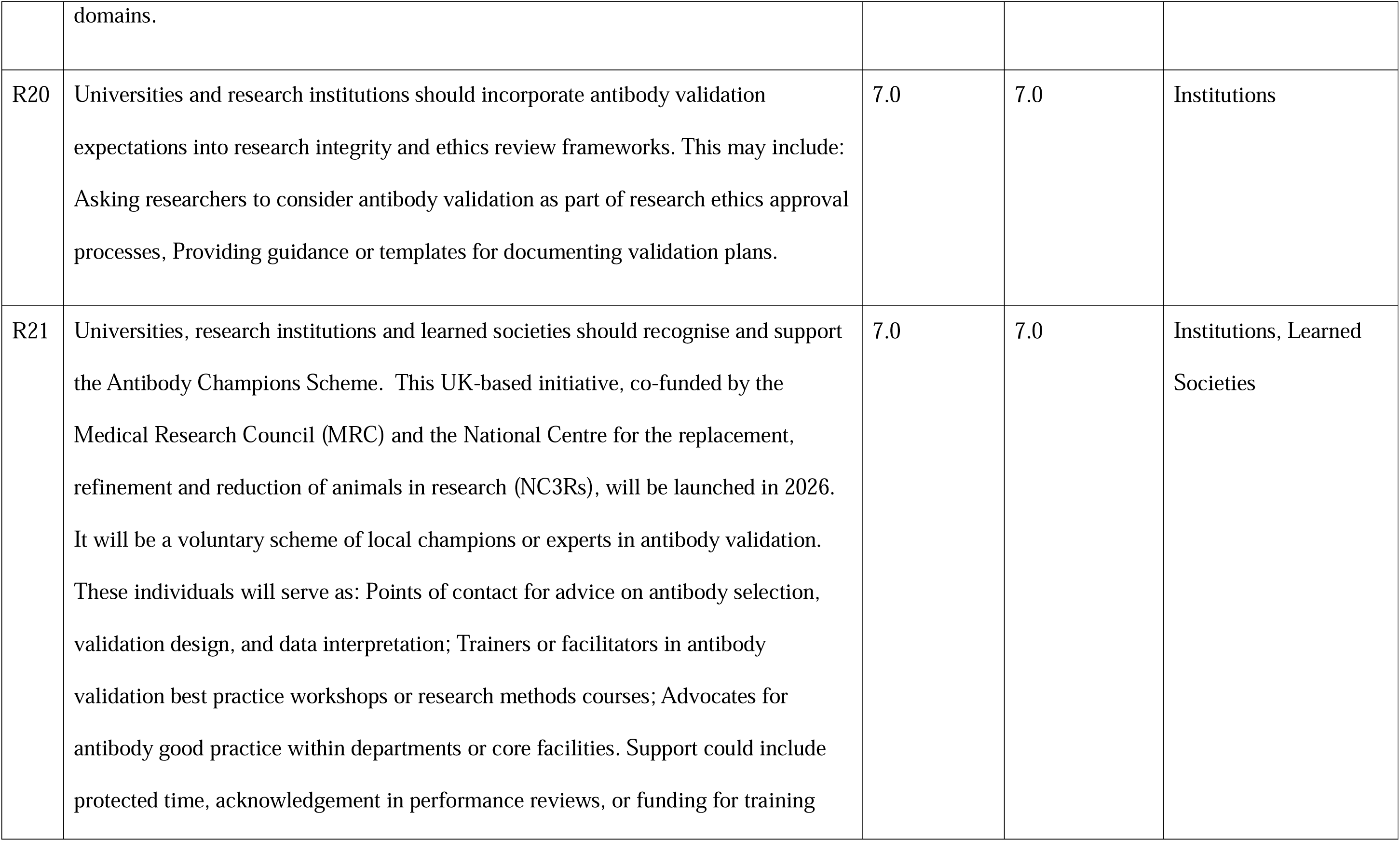
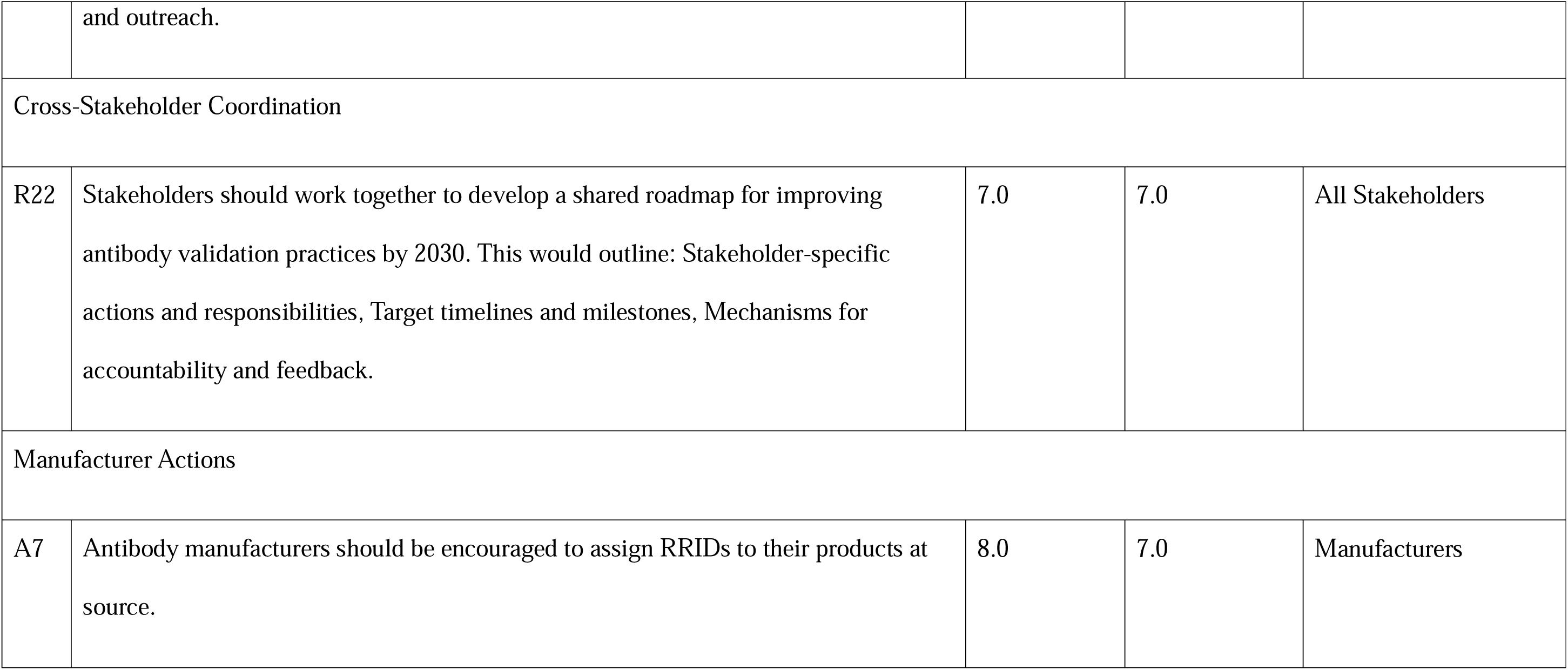
Consensus Recommendations - Items Rated Both Effective and Feasible with agreement.

The panel rated 17 items as uncertain on at least one dimension (Table 2). In 15 of these 17 cases, items were rated as effective (median ≥7) but with uncertain feasibility (median 4–6), indicating that while panellists believed these actions would improve antibody practices, they had reservations about realistic implementation by 2030. This pattern, high confidence in effectiveness, uncertainty about feasibility, was the dominant finding for items not achieving consensus, and is a recurring theme in the stakeholder-specific recommendations presented below. One item was rejected by the panel, with concerns around both effectiveness and feasibility (Table 3). A summary of the ratings across all items is presented in Figure 2. A consolidated summary grouping all items by stakeholder group and rating status is provided in S4 Table.

**Figure 2.**
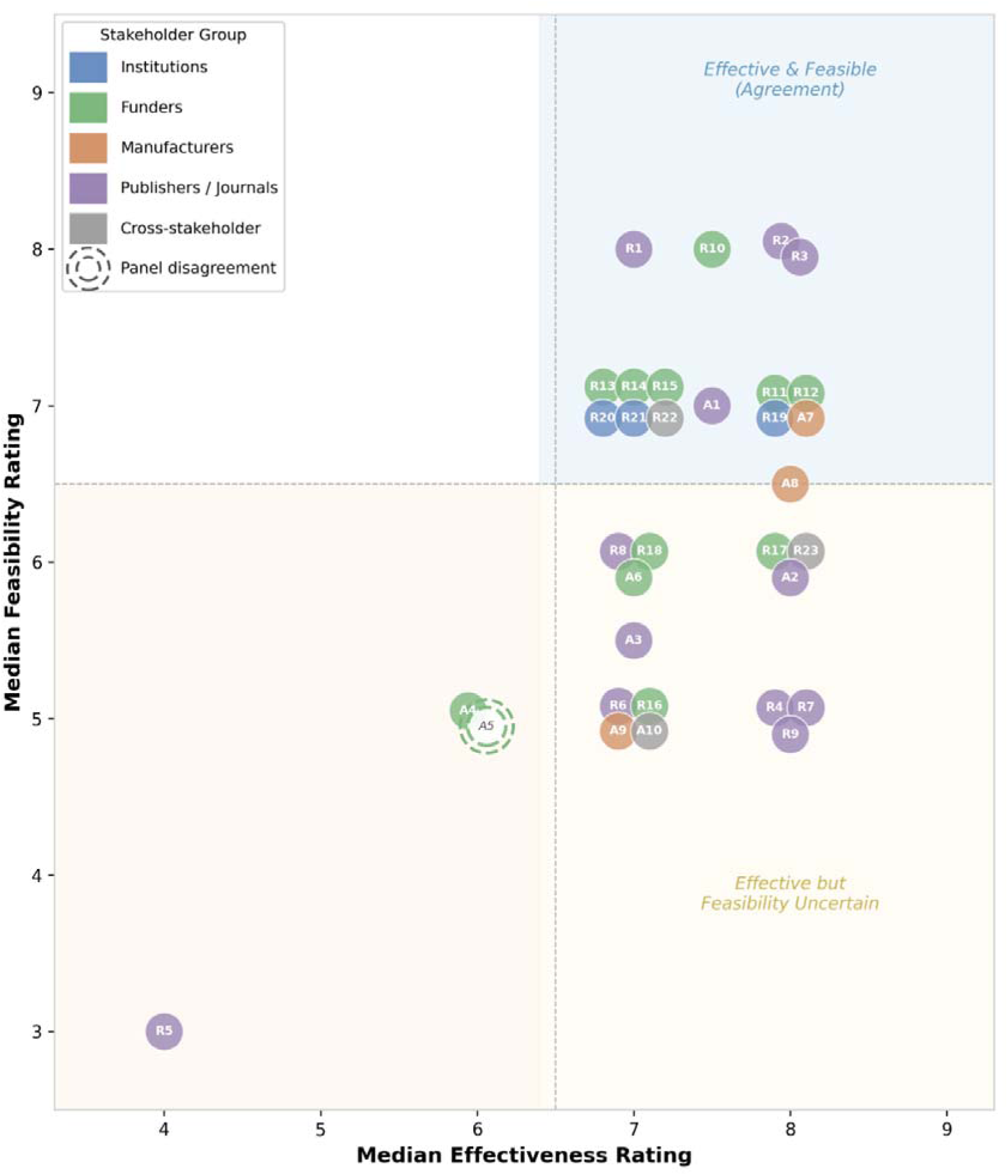
Effectiveness and feasibility ratings for all 33 proposed actions. Each point represents one proposed action, plotted by median effectiveness (x-axis) and median feasibility (y-axis) ratings on 9-point scales. Points are colour-coded by stakeholder group. Dashed lines indicate the boundary between ratings classified as effective/feasible (median ≥7) and uncertain (median 4–6), following our RAND/UCLA round-down rule for boundary medians of 6.5. The upper-right quadrant contains items rated as both effective and feasible with agreement (15 items). The lower-right quadrant contains items rated as effective but with uncertain feasibility (15 items). Item with panel disagreement (A5, IPR > IPRAS) is shown as open circle with dashed borders; all other items achieved agreement. One item (R5) was rejected by the panel (uncertain effectiveness, infeasible). Points at identical coordinates have been slightly offset for readability.

**TABLE 2:**
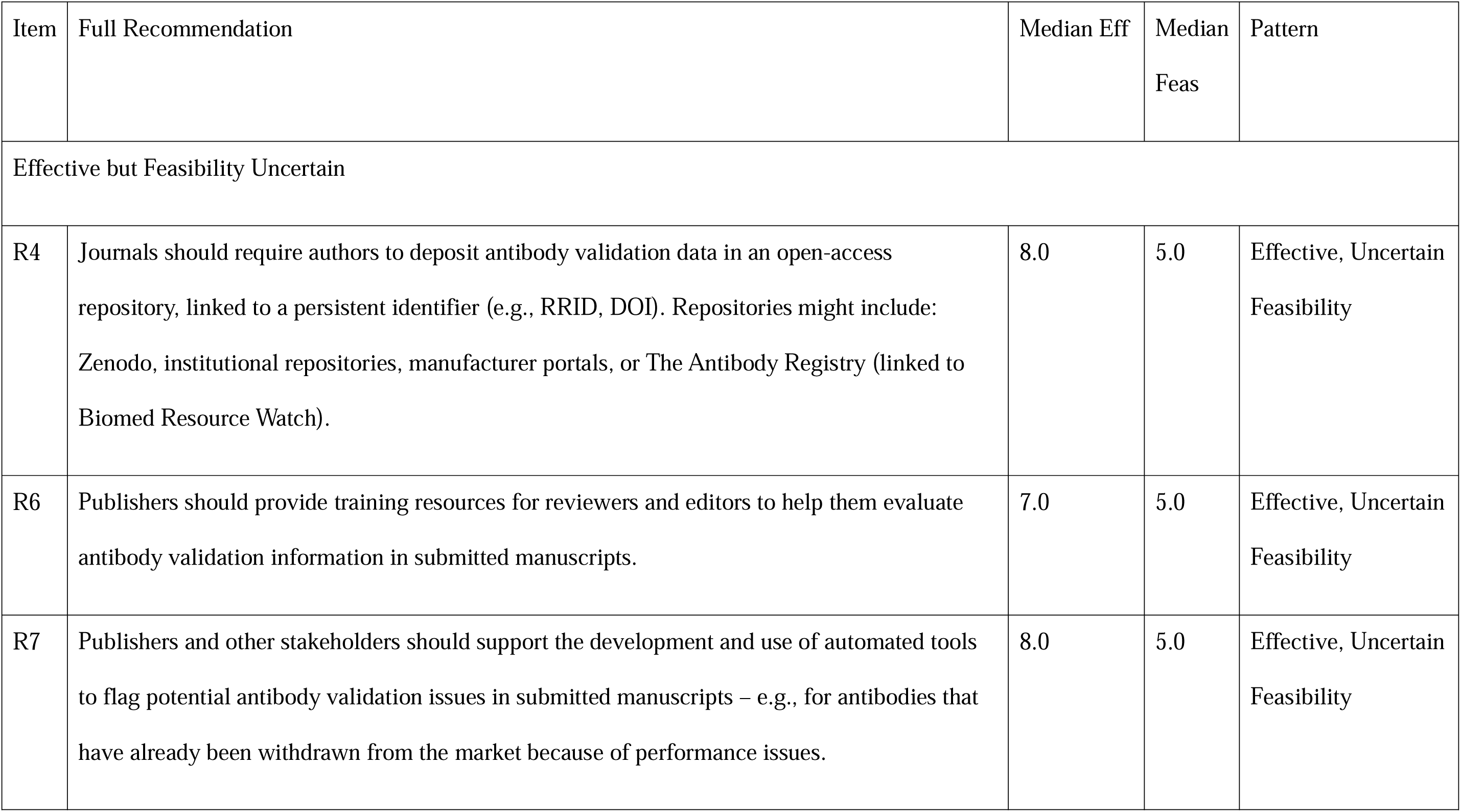
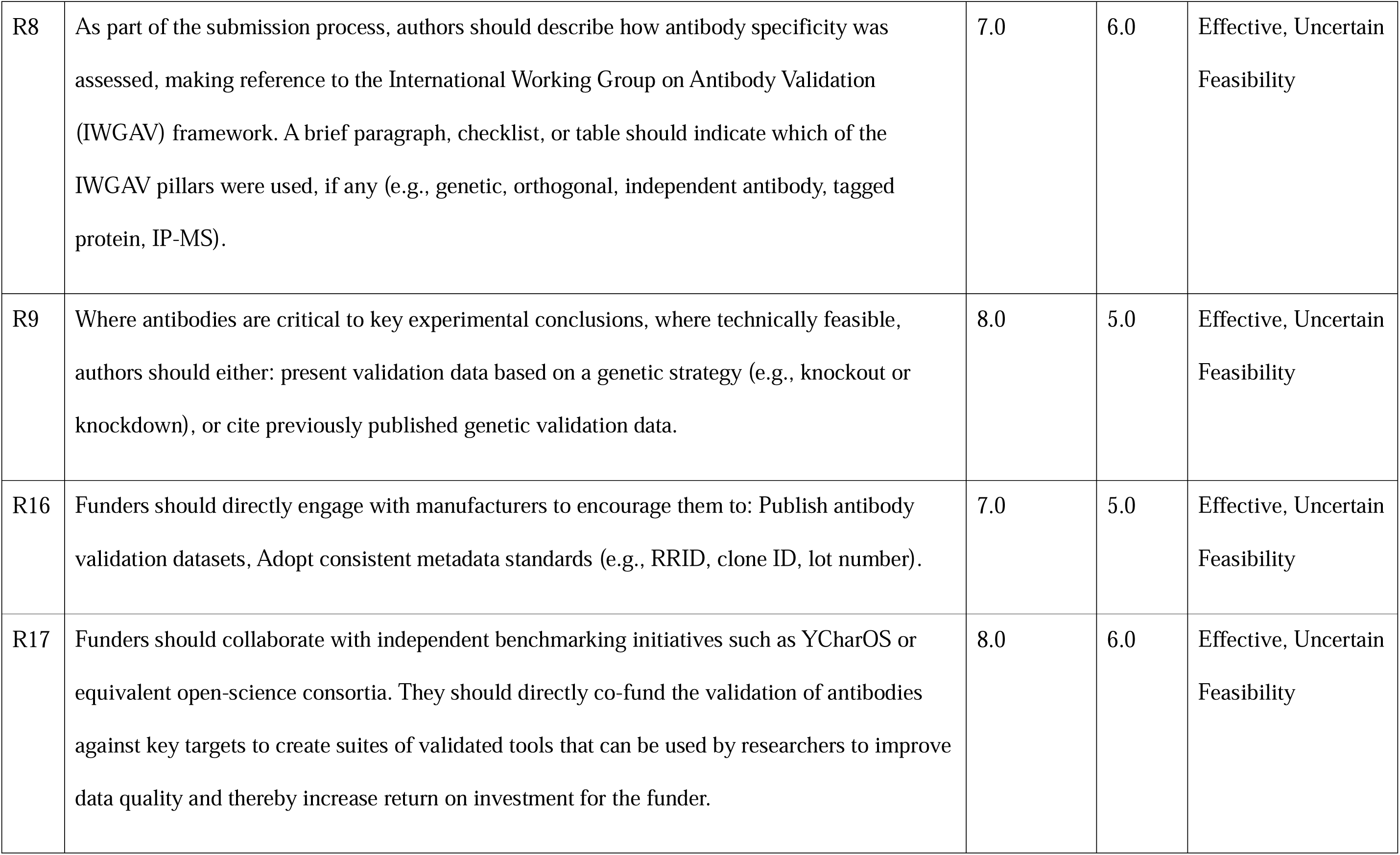
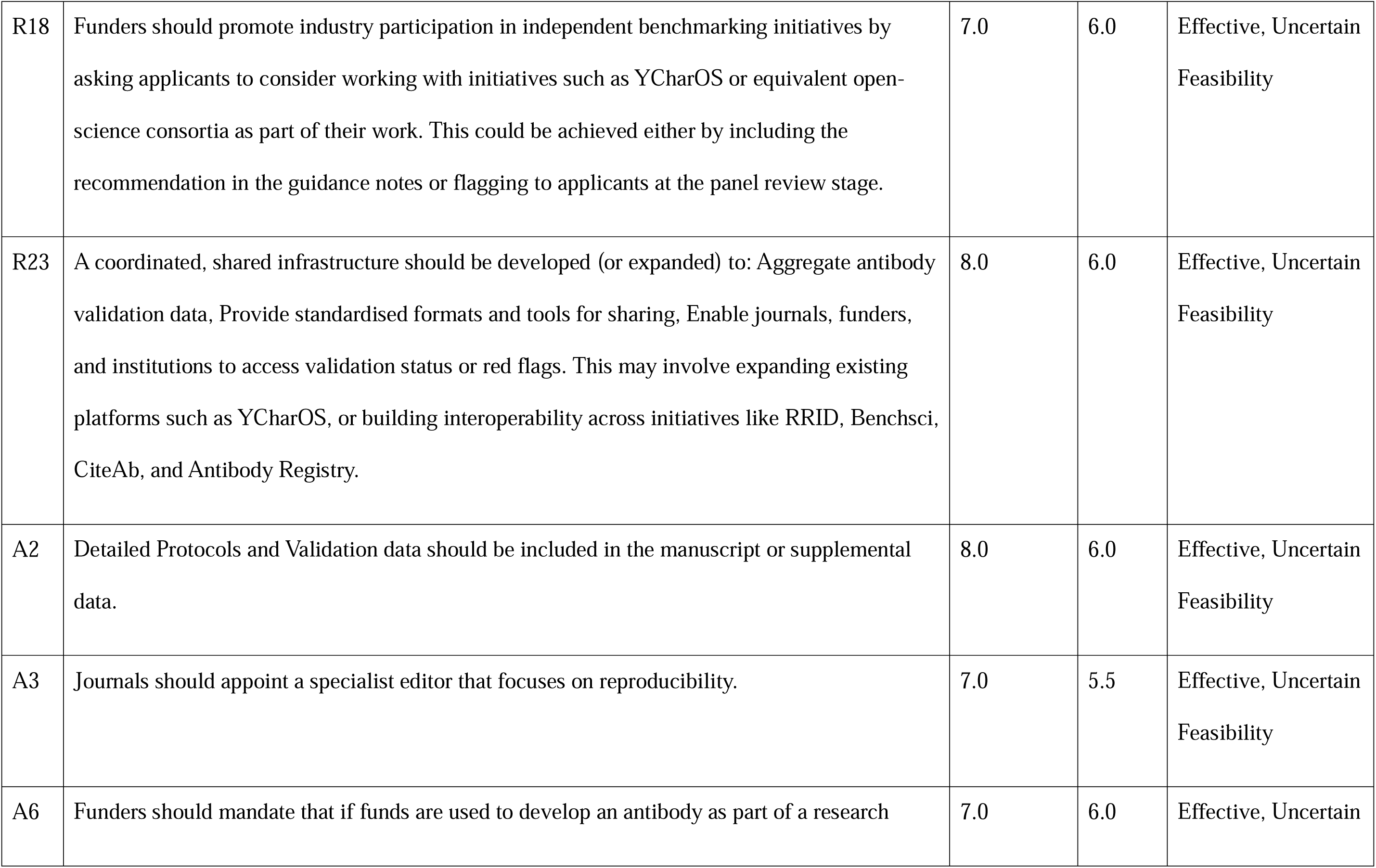
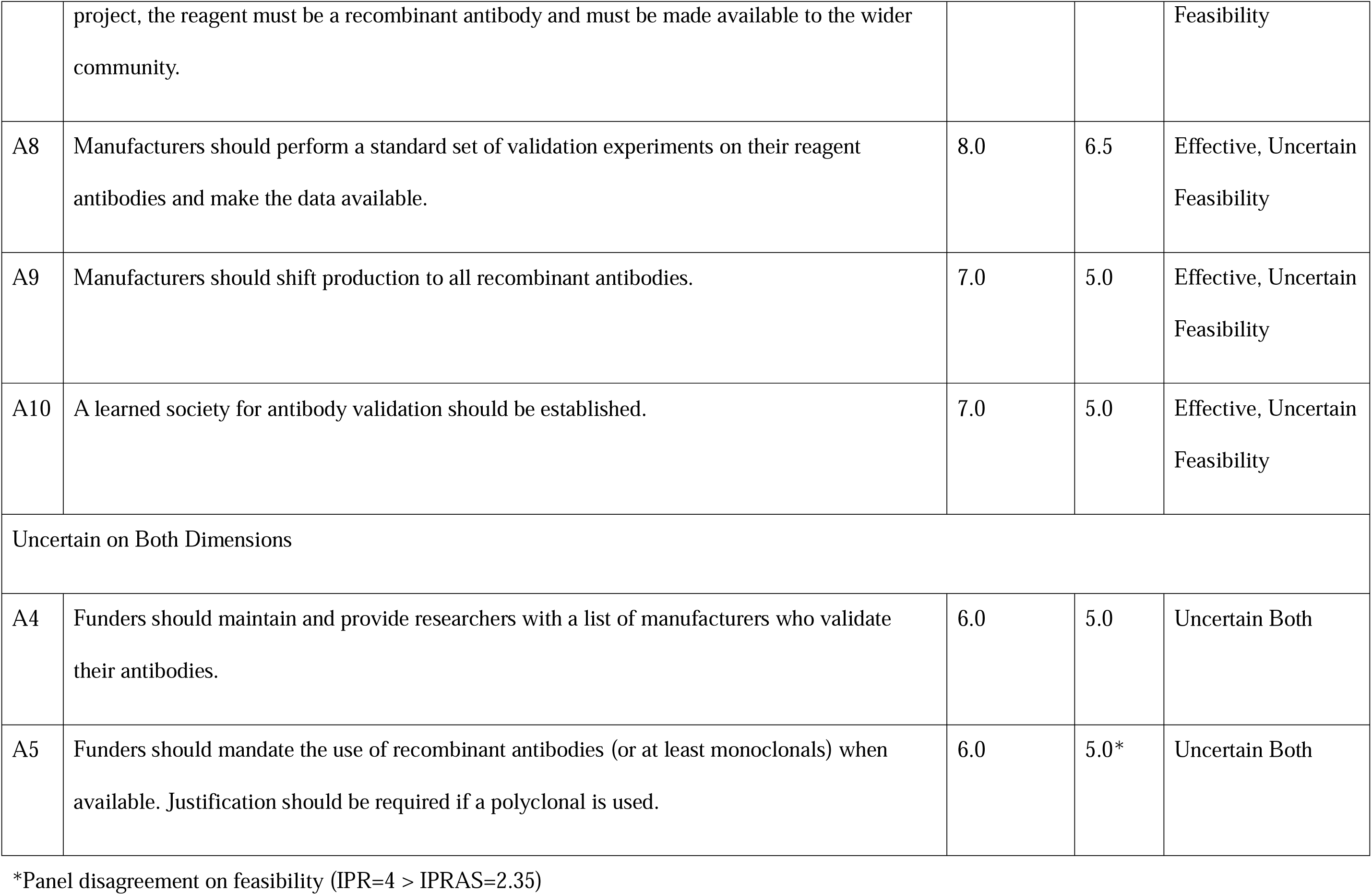
Items with Agreed Uncertain Effectiveness and/or Feasibility.

**TABLE 3:**
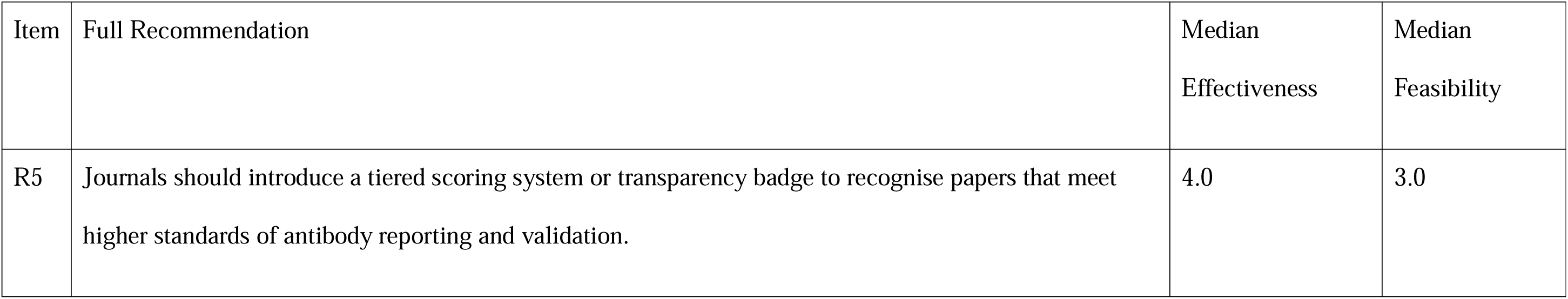
Rejected Recommendation - Item Rated Uncertain effectiveness, and Infeasible, with agreemen.

| Item | Full Recommendation | Median Effectiveness | Median Feasibility |
| --- | --- | --- | --- |
| R5 | Journals should introduce a tiered scoring system or transparency badge to recognise papers that meet higher standards of antibody reporting and validation. | 4.0 | 3.0 |

Consistent with the criteria set out in advance for progressing between rounds, the research team reviewed the Round 2 results to determine whether a further round was warranted. This decision was based on three convergent factors. First, a high proportion of items (15 of 33, 45.5%) had achieved clear consensus as both effective and feasible across the two rounds. Second, disagreement remained minimal, with disagreement being present on only one item in Round 2. Third, rating stability between rounds was high: among the 12 items re-rated in Round 2, median effectiveness ratings remained identical for 9 items (75.0%) and median feasibility ratings remained identical for 10 items (83.3%), with no item shifting by more than 1 point on either scale. The minimal rating changes between rounds indicated that further iteration would be unlikely to resolve the persistent pattern of items rated as effective but with uncertain feasibility. Full quantitative results for all items across both rounds are presented in S2 and S3 Tables, with complete qualitative commentary provided in S7 and S8 Texts.

### Stakeholder-Specific Recommendations

The following sections present the consensus recommendations, predicted to be both effective and feasible, and items predicted to be effective but with uncertain feasibility for implementation by 2030, organised by stakeholder group. For each recommendation, we provide the panel’s quantitative assessment alongside key themes from the qualitative feedback that contextualise the ratings and inform implementation considerations. Full qualitative analysis for the 15 consensus recommendations, is presented in S1 Text, and detailed stakeholder consultation drafts including implementation options are provided in S2– 5 Texts.

### Key Recommendations for Institutions

##### Summary of Institutional Recommendations

***Consensus recommendations (effective and feasible by 2030):***

- R19: Training in antibody validation in relevant bioscience courses and programmes (Eff 8.0 / Feas 7.0)
- R20: Incorporate antibody validation into research integrity and ethics review frameworks (Eff 7.0 / Feas 7.0)
- R21: Recognise and support local champions or experts in antibody validation (Eff 7.0 / Feas 7.0)

*All three institutional items achieved full consensus (ie rated effective and feasible without disagreement). No items with uncertain feasibility. This is unique among stakeholder groups.*

Median ratings: Eff = Effectiveness; Feas = Feasibility; rated on 9-point scales.

The panel viewed institutional actions as foundational to the broader reform effort, with one panellist noting that “without laying such a groundwork of training and advocacy, antibody validation will never become the norm.“

Training in antibody validation received strong endorsement (median effectiveness 8.0, feasibility 7.0; R19). The panel recommended that training should be offered in relevant bioscience courses and programmes, including undergraduate education in fields such as immunology, neuroscience, and molecular biology, postgraduate and doctoral training, and continuing professional development. Importantly, the panel emphasised that training should be field-appropriate rather than universally mandated across all domains. Panel members highlighted the need to address laboratory culture alongside formal training, noting that “*in many cases the emphasis on validation and proper controls comes down to lab culture. If a student is trained to validate antibodies but the PI/senior lab members do not agree this is a priority, it will not be done*.” The need to train established researchers as well as trainees was a recurring theme: “*Need to train the senior staff as well!“*

The panel endorsed incorporating antibody validation expectations into institutional research integrity and ethics review frameworks (median effectiveness 7.0, feasibility 7.0; R20). This could include asking researchers to consider antibody validation as part of research ethics approval processes and providing guidance or templates for documenting validation plans.

However, panel members debated the appropriate framing, with some questioning whether antibody validation constitutes an ethical issue in the traditional sense: “*I don’t really see this as an ethical issue. Lumping it into ethics with big issues like image manipulation, data fabrication, animal use, etc doesn’t seem appropriate*.” Others noted that the waste of animal and human samples resulting from poor antibody validation carries clear ethical implications. Panel members also noted that “*a lot of research does not go through ethics committees,*” emphasising the importance of complementary mechanisms beyond ethics review alone.

The third institutional recommendation called for universities, research institutions, and learned societies to recognise and support local champions or experts in antibody validation (median effectiveness 7.0, feasibility 7.0; R21). These individuals could serve as points of contact for advice on antibody selection and validation design, trainers or facilitators in reproducibility workshops, advocates for good practice within departments or core facilities, and contributors to cross-institutional reproducibility networks. Support could include protected time, acknowledgement in performance reviews, or funding for training and outreach. The panel viewed this as foundational, for example noting that “*all of the questions in this section are not going to make antibody research more reproducible per se but will lay the foundation for increased awareness and adoption of best practice*.” However, concerns about institutional buy-in and sustainability were raised, with panellists questioning how to maintain expertise networks given high turnover of research staff and limited institutional awareness of the problem.

### Key Recommendations for Funders

##### Summary of Funder Recommendations

***Consensus recommendations (effective and feasible by 2030):***

- R10: Require antibody validation plans in grant applications (Eff 7.5 / Feas 8.0)
- R11: Dedicated budget line for antibody validation in funded projects (Eff 8.0 / Feas 7.0)
- R12: Create or expand targeted tool development and validation schemes (Eff 8.0 / Feas 7.0)
- R13: Signal importance of antibody validation in applicant guidance (Eff 7.0 / Feas 7.0)
- R14: Encourage deposition of validation data in open-access repositories (Eff 7.0 / Feas 7.0)
- R15: Formally endorse community-developed reporting standards (e.g., IWGAV, MDAR) (Eff 7.0 / Feas 7.0)

***Effective but feasibility uncertain (4 items):***

- R17: Co-fund independent benchmarking initiatives (Eff 8.0 / Feas 6.0)
- R18: Promote researcher participation in benchmarking (Eff 7.0 / Feas 6.0)
- R16: Directly engage manufacturers on transparency (Eff 7.0 / Feas 5.0)
- A6: Mandate funded antibodies be recombinant and publicly available (Eff 7.0 / Feas 6.0)

Median ratings: Eff = Effectiveness; Feas = Feasibility; rated on 9-point scales.

The panel viewed funders as occupying a critical position in the research lifecycle: before experiments are conducted, when validation planning and budgeting can be embedded from the outset (full qualitative comments in S7 and S8 Texts).

The panel reached consensus that funders should provide dedicated financial support for antibody validation. This encompasses two complementary actions: applicants to biomedical funding schemes should include a specific budget line item for resources to conduct robust antibody validation where relevant to their proposal (median effectiveness 8.0, feasibility 7.0; R11), and funders should create or expand targeted schemes to support the development and validation of critical antibody-based research tools (median effectiveness 8.0, feasibility 7.0; R12). Both items received the highest effectiveness ratings among funder recommendations. Panel members noted, however, that budget constraints could pose challenges, with one participant observing that *“research proposals typically employ multiple antibodies (5–10+). Would the applicant be required to validate all antibodies prior to use? This would result in a pretty significant budget increase.“*

The panel also endorsed embedding antibody validation expectations into grant application processes through escalating approaches. At a minimum, funders should include information in applicant guidance that antibody performance is an important limitation in many experimental methods and encourage applicants to address this in methodological sections (median effectiveness 7.0, feasibility 7.0; R13). More substantively, funders should include a section in application forms where applicants must detail the steps they will take to validate the antibodies used (median effectiveness 7.5, feasibility 8.0; R10). Panel members nevertheless cautioned against box-ticking: *“Another box ticking exercise for the applicant, which is filled with little care and using generic text each time.“* Panel members cautioned that effectiveness would depend on whether validation plans are genuinely evaluated during review.

Two further consensus recommendations addressed community standards and data sharing. Funders should formally endorse community-developed reporting standards that promote antibody transparency and validation, such as those from the IWGAV and MDAR frameworks (median effectiveness 7.0, feasibility 7.0; R15). Funders should also encourage grantees to deposit antibody validation data in open-access repositories, ideally linked to RRIDs or antibody registry entries (median effectiveness 7.0, feasibility 7.0; R14). Panel members noted that endorsement alone, while a *“feasible and low lift“* action, may be insufficient without accompanying enforcement mechanisms, and expressed concern about the quality of deposited data without robust review processes.

Beyond the consensus recommendations, co-funding of independent benchmarking initiatives such as YCharOS were deemed effective (median effectiveness 8.0), but feasibility remained uncertain (median 6.0; R17). Panel members agreed that creating suites of validated tools through independent benchmarking would substantially improve data quality and increase return on investment, but expressed concern about alignment with funder mandates and funding mechanism constraints. Several additional funder recommendations were rated as effective but with feasibility concerns, including promoting researcher participation in benchmarking (median effectiveness 7.0, feasibility 6.0; R18), mandating that funded antibodies be recombinant and publicly available (median effectiveness 7.0, feasibility 6.0; A6), and engaging manufacturers directly on transparency (median effectiveness 7.0, feasibility 5.0; R16).

### Key Recommendations for Manufacturers

##### Summary of Manufacturer Recommendations

***Consensus recommendation (effective and feasible by 2030):***

- A7: Assign RRIDs to products at source (Eff 8.0 / Feas 7.0)

***Effective but feasibility uncertain (2 items):***

- A8: Perform standard validation experiments and make data available (Eff 8.0 / Feas 6.5)
- A9: Shift production toward recombinant antibodies (Eff 7.0 / Feas 5.0)

*One consensus item; feasibility barriers for remaining items reflect commercial viability and market dynamics. S5 Text for consultation draft.*

Median ratings: Eff = Effectiveness; Feas = Feasibility; rated on 9-point scales.

The manufacturer landscape presented a distinctive pattern: one recommendation achieved full consensus, while the remaining items were rated as effective but with significant feasibility barriers reflecting the commercial context in which manufacturers operate. Panel members, including manufacturer representatives, repeatedly acknowledged that leading manufacturers already exemplify good practice, while others lag behind.

The panel reached consensus that antibody manufacturers should assign Research Resource Identifiers (RRIDs) to their products at source (median effectiveness 8.0, feasibility 7.0; A7). This was the only manufacturer recommendation to achieve full consensus, and the response was notably enthusiastic, with some panellists describing it as *“really, an easy one“* and noting it *“would be fantastic.“* RRIDs assigned at source would enable researchers to uniquely identify and cite specific antibodies, facilitating systematic tracking of antibody performance across the literature and supporting automated screening of submissions by journals. Panel members acknowledged, however, that some suppliers may not cooperate: *“There are some suppliers that will ignore this, in the same way they have ignored the calls to improve validation.“*

Manufacturers performing a standard set of validation experiments and making the data available was rated as effective (median effectiveness 8.0) but with uncertain feasibility (median 6.5; A8). The tension between rigorous context-specific validation and commercial viability was central to panel deliberations. Some panellists were emphatic that this was unrealistic, e.g. with one noting “*Not going to happen. Too expensive*”, while others noted that “*reputable manufacturers are already doing this*” and that “*validation is a continual/on-going process for every antibody*.” A shift toward recombinant antibody production was similarly rated as effective (median 7.0) but with uncertain feasibility (median 5.0; A9), with the panel noting that manufacturer production decisions are ultimately driven by customer demand.

### Key Recommendations for Publishers and Journals

##### Summary of Publisher and Journal Recommendations

***Consensus recommendations (effective and feasible by 2030):***

- R1: Authors include RRIDs for each antibody used (Eff 7.0 / Feas 8.0)
- R2: Authors provide complete antibody metadata for unambiguous identification (Eff 8.0 / Feas 8.0)
- R3: Authors report dilution ratio and protein concentration (Eff 8.0 / Feas 8.0)
- A1: Journals establish clear antibody validation and reporting standards (Eff 7.5 / Feas 7.0)

***Effective but feasibility uncertain (4 items, all Eff ≥7.0):***

- R4: Require validation data in open-access repositories (Eff 8.0 / Feas 5.0)
- A2: Include validation protocols in manuscripts or supplements (Eff 8.0 / Feas 6.0)
- R7: Automated tools to flag antibody validation issues (Eff 8.0 / Feas 5.0)
- R9: Genetic validation for critical antibodies (Eff 8.0 / Feas 5.0)
- R6: Train reviewers and editors on validation assessment (Eff 7.0 / Feas 5.0)
- A3: Appoint specialist reproducibility editors (Eff 7.0 / Feas 5.5)

*Four consensus items centred on reporting requirements. Other effective items face barriers around editorial resources, cost, and infrastructure. See S1 Text for full analysis of consensus items; S2 Text for consultation draft.*

Median ratings: Eff = Effectiveness; Feas = Feasibility; rated on 9-point scales.

Publisher recommendations reflected a common pattern: the panel agreed that several interventions would substantially improve research quality but identified practical barriers around editorial resources, cost, and infrastructure. Four items achieved full consensus, while four additional items deemed effective had uncertain feasibility.

The panel reached consensus on a package of reporting requirements that together enable unambiguous antibody identification and experimental replication. Authors should include Research Resource Identifiers for each antibody used where available (median effectiveness 7.0, feasibility 8.0; R1), provide sufficient metadata to allow unambiguous identification of each antibody including clone ID, catalogue number, lot number, and vendor (median effectiveness 8.0, feasibility 8.0; R2), and report the dilution ratio and protein concentration where possible for each antibody used (median effectiveness 8.0, feasibility 8.0; R3). These three items work synergistically: complete metadata enables antibody identification, while RRIDs provide persistent identifiers that link to characterisation databases and the broader literature. The panel also reached consensus that journals should establish clear standards for antibody validation and reporting that authors must meet (median effectiveness 7.5, feasibility 7.0; A1), noting that this would only be maximally effective *“if there were community standards already in place or consensus across most journals.“*

The panel rated both the deposition of validation data in open-access repositories (median effectiveness 8.0, feasibility 5.0; R4) and the inclusion of detailed validation protocols in manuscripts or supplementary data (median effectiveness 8.0, feasibility 6.0; A2) as effective, but with uncertain feasibility. Similarly, the development and use of automated tools to flag potential antibody validation issues in submitted manuscripts were predicted to be effective, but again uncertain feasibility (median effectiveness 8.0, feasibility 5.0; R7). Panel members viewed automated tools as potentially transformative but questioned who would fund their development and maintenance, and noted that *“not all publishers/journals have the financial means“* to implement them. The requirement that authors present genetic validation data for critical antibodies (median effectiveness 8.0, feasibility 5.0; R9) was rated as effective but faced feasibility barriers around technical limitations (essential genes cannot be knocked out), cost, and the difficulty of defining which antibodies are “critical” to a study’s conclusions.

Building editorial capacity for validation assessment was rated as effective but with feasibility concerns. Both training reviewers and editors to evaluate antibody validation information (median effectiveness 7.0, feasibility 5.0; R6) and appointing specialist reproducibility editors (median effectiveness 7.0, feasibility 5.5; A3) faced scepticism about reviewer engagement and publisher resources. Panellists doubted that reviewers would invest time in additional training given already onerous review obligations, and many journals were seen as lacking resources for specialist editor positions. The only item rejected by the panel across both rounds was the introduction of a tiered scoring system or transparency badge for antibody reporting (R5), which declined from uncertain to infeasible between rounds, reflecting scepticism about badge-based incentive schemes in this context.

### Cross-Stakeholder Consensus Items

##### Summary of Cross-Stakeholder Items

***Consensus recommendation:***

- R22: Develop a shared roadmap for improving antibody validation practices by 2030 (Eff 7.0 / Feas 7.0)

***Effective but feasibility uncertain:***

- R23: Coordinated shared infrastructure for aggregating antibody validation data (Eff 8.0 / Feas 6.0)
- A10: Establish a learned society for antibody validation (Eff 7.0 / Feas 5.0)

Median ratings: Eff = Effectiveness; Feas = Feasibility; rated on 9-point scales.

Two cross-stakeholder items merit specific attention. The panel reached consensus that stakeholders should work together to develop a shared roadmap for improving antibody validation practices by 2030 (median effectiveness 7.0, feasibility 7.0; R22). This roadmap would outline stakeholder-specific actions and responsibilities, target timelines and milestones, and mechanisms for accountability and feedback.

The development of coordinated, shared infrastructure for aggregating antibody validation data was deemed potentially effective (median effectiveness 8.0) but uncertain feasibility (median 6.0; R23). This infrastructure would aggregate validation data, provide standardised formats and tools for sharing, and enable journals, funders, and institutions to access validation status or identify known problem antibodies. Panel members viewed this as potentially transformative, supporting several other recommendations, from automated manuscript screening to informed grant review, but recognised the substantial coordination and resource challenges involved. Several recommendations across stakeholder groups would benefit significantly from such shared infrastructure, suggesting that its development could unlock a cascade of downstream improvements.

The establishment of a learned society for antibody validation was rated as effective (median 7.0) but with uncertain feasibility (median 5.0; A10). While the panel recognised the potential value of a professional body to coordinate standards, training, and advocacy, concerns were raised about sustainability and the risk of duplicating existing networks.

## Discussion

This Delphi study engaged 32 international experts from across the research ecosystem to develop a formally prioritised, consensus-based set of recommendations for addressing antibody validation failures in biomedical research. The panel achieved consensus on 15 items as both effective and feasible for implementation by 2030, spanning institutional, funder, manufacturer, and publisher actions as well as cross-stakeholder coordination. An additional 15 items were rated as effective but with uncertain feasibility, reflecting a consistent pattern in which experts were confident that proposed interventions would improve antibody practices but less certain about realistic implementation within the target timeframe. These findings complement existing technical solutions [1], data-sharing initiatives [17, 18], and emerging stakeholder coordination efforts [12, 18]. They represent the prioritised judgement of a single, predefined expert panel convened across the relevant areas of expertise, rather than a definitive or globally generalisable standard, and are offered to inform stakeholder decisions. Because feasibility in particular depends on local policy, infrastructure and funding environments, readers working in differently configured research systems are best served by treating the recommendations as expert-prioritised options to weigh against their own conditions rather than as conclusions established for their setting.

### Barriers to Implementation

From the panel’s free-text comments, we interpret four interconnected barriers that help explain both the persistence of the antibody validation problem and the challenges of implementing solutions. These are offered as our interpretation of the qualitative material rather than as formally coded findings.

First, diffuse ownership enables collective inaction. Antibody validation failures impose costs across the research system — wasted funding, unreliable literature, consumed biological materials — but no single stakeholder bears the full cost or has authority to mandate change unilaterally. Researchers lack incentives to validate when publication and funding systems do not reward the practice; publishers hesitate to impose requirements that could divert submissions to less demanding competitors; funders may view validation as outside their mandate; and manufacturers competing on price and catalogue breadth face limited incentives to invest in validation when researchers continue to purchase on the basis of previous use and supplier reputation [14]. Panel members described a pattern of mutual deflection, with responsibility for validation perceived as belonging to other stakeholder groups rather than one’s own (S7 Text, R22). Several characterised the broader landscape as too fragmented for meaningful alignment to feel achievable. However, the panel’s endorsement of a shared roadmap (R22) and the interdependencies across recommendations point toward a countervailing dynamic: participants noted that journal requirements would create *de facto* mandates for manufacturers, that researcher demand would shift manufacturer behaviour, and that funder investment in infrastructure would enable practices that other stakeholders could then require.

Second, market dynamics create structural tensions. The research antibody market operates predominantly under a Research Use Only model that prioritises catalogue breadth and low unit cost over rigorous validation. Manufacturer representatives acknowledged significant economic barriers to comprehensive validation, describing the costs as prohibitive and warning that extensive validation programmes would reduce revenue without a corresponding market incentive (S7 Text, A8). At the same time, one manufacturer described their company’s ongoing commitment to validation, illustrating that the barrier is structural rather than universal. Yet the apparent low cost of inadequately validated antibodies is illusory when downstream costs are considered: failed experiments, repeated purchasing, wasted samples, and unreliable findings impose far greater costs on the research system than better validation would. Creating market conditions in which quality is rewarded — through purchasing decisions informed by independent characterisation data, journal requirements that make validation data visible, and funder signals that prioritise quality — is necessary to shift the equilibrium.

Third, resource justification challenges persist despite clear return on investment. Participants noted competing demands on limited resources, with some questioning whether funders would prioritise antibody validation given the breadth of competing demands on their budgets, even while acknowledging the value of the proposed actions (S8 Text, A10). The economic case is compelling at a system level — the estimated $1 billion annually in direct waste in the US alone [12, 13], plus far greater opportunity costs — but the return is diffuse and difficult for individual stakeholders to capture. A funder investing in validation infrastructure benefits the entire community, not just its own grantees; an institution investing in training produces graduates who carry improved practices elsewhere. The benefits are real but distributed across time and across the ecosystem, and compounded by the difficulty of demonstrating negative outcomes that were prevented.

Fourth, coordination challenges reflect the scale of the problem. With more than 22,000 human protein targets, millions of antibody products, and a global research community, the coordination challenge is immense. No single authority can mandate standards, and voluntary adoption requires alignment across actors with different incentives, timescales, and resources. The panel’s findings suggest, however, that complete alignment is not a prerequisite for progress. The 15 consensus recommendations represent actions individual stakeholders can implement without waiting for others, and participants emphasised sequencing over simultaneity: several argued that funders must first act as enablers — establishing resources, infrastructure, and training — before mandates can be effective (S8 Text, A5), while others suggested that even imperfect initial standards would allow journals to demonstrate a credible commitment to improving antibody reporting practices (S8 Text, A1).

These barriers are consistent with recent multi-stakeholder commentaries. Kahn et al. [12] noted that the origins of the antibody crisis are ‘many and varied’ and that ‘it is unrealistic to expect a complete solution in the near future’, while emphasising that significant progress is achievable through concerted action. In earlier work by members of this study team [18], we argued that behavioural and cultural change could be supported by engaging stakeholders at every level using established frameworks for research culture change. The Delphi findings complement these analyses by providing quantitative evidence of where expert agreement is strongest and where feasibility barriers remain.

### Next Steps and Implementation

The findings lend themselves to a programme of targeted stakeholder consultation. To support this, we have prepared separate documents for each stakeholder group — publishers, funders, institutions, and manufacturers — presenting consensus recommendations alongside implementation options derived from the panel’s qualitative feedback (S2–5 Texts). These present options rather than prescriptions, recognising that the optimal approach will vary across organisations, countries, and contexts. Items rated as effective with uncertain feasibility are not dismissed but treated as implementation challenges to be addressed through consultation. Interdependencies across stakeholder groups are made explicit throughout, supporting coordinated rather than siloed implementation.

Several elements of the recommended infrastructure already exist. The NC3Rs Antibody Champions programme provides a model for the local champions recommendation (R21) and delivers masterclasses for doctoral training aligned with R19. The OGA Academy offers structured online training [18], the OGA Antibody Database provides searchable characterisation data, and YCharOS continues to expand its open characterisation pipeline [1]. The eLife stakeholder roadmap [12] proposed actions that align closely with many consensus items, and the RRID initiative [17] provides identification infrastructure on which several publisher-focused recommendations depend. These resources demonstrate that the recommendations can be implemented by building on proven approaches. Notably, several recommendations are interdependent: shared infrastructure for aggregating validation data (R23) would support automated manuscript screening (R7), informed grant review (R10), and evidence-based purchasing decisions, suggesting that investment in infrastructure could unlock a cascade of downstream improvements across stakeholder groups. As an early test of how these coordinating mechanisms might work in practice, a networked antibody champions model — embedding trained local leads within institutions to deliver training and support reagent-selection decisions — is currently being piloted across UK institutions, providing an initial operationalisation of the local-champion and roadmap recommendations (R21, R22).

To illustrate how the recommendations translate into practice, consider the consensus item asking journals to establish clear standards for antibody validation and reporting that authors must meet (A1). The consensus wording is deliberately non-prescriptive, but the panel’s qualitative feedback and the accompanying publisher consultation document (S2 Text) indicate what such standards could contain. A minimal baseline would adopt the validation and reporting language already present in the MDAR framework [16], which covers RRIDs, vendor, catalogue number, clone or lot number, and dilution. Journals could strengthen this by asking authors to indicate which IWGAV-aligned evidence [15] supports each antibody-dependent conclusion, and by requiring authors to state explicitly where context-specific validation is absent. Journals could also ask authors to supply the validation data underpinning antibody-dependent conclusions, either within the manuscript or its supplementary material or by deposition in an open repository (R4, A2). Where generating new validation data is impractical, authors could be permitted to cite previously published validation evidence obtained under closely matched experimental conditions, or to reference independent characterisation resources such as the knockout-controlled datasets generated by YCharOS [1] and aggregated in the OGA Antibody Database.

### Limitations

Several limitations should be considered. Participants were identified through existing professional networks, and the panel may therefore share perspectives that do not fully represent the diversity of views across the global research community. The panel was weighted toward the United Kingdom and United States, and the biomedical research ecosystem in other countries may face different challenges. Some sectors (particularly research funders, n=4) had few representatives, meaning feasibility assessments for funder-directed recommendations may not capture the full diversity of perspectives. The high retention rate (93.8%) and low disagreement suggest a relatively cohesive panel, which may underestimate the range of views from a broader consultation.

The Delphi method captures expert opinion at a specific point in time and does not constitute empirical evidence of effectiveness. The consensus recommendations represent the panel’s assessment of what is likely to be effective and feasible, but actual effectiveness will depend on implementation context. The panel reached consensus after two rounds, which limited iterative refinement where uncertainty persisted. However, rating stability between rounds suggests that additional rounds would have been unlikely to shift results substantially. The qualitative data were synthesised narratively rather than through formal thematic analysis, consistent with RAND/UCLA practice where free-text comments inform item interpretation rather than serving as independent qualitative data. Full qualitative commentary is provided in S7 and S8 Texts.

## Conclusions

This Delphi study provides cross-stakeholder feasibility ratings with an explicit implementation horizon of 2030 and also offers prioritisation across sectors. The rating of ‘effective but uncertain feasibility’ that characterises many of the proposed actions should not be interpreted as reason for inaction. It reflects an honest assessment that the pace and sequencing of change matter as much as the direction. The 15 consensus recommendations provide an evidence-based starting point for immediate action; the remaining items rated as effective represent a second wave whose feasibility is likely to increase as early implementation builds infrastructure, shifts incentives, and establishes new norms. This staged approach — enablement to expectation to mandate — was consistently endorsed by the panel and is reflected in the consultation documents (S2–5 Texts). Together with existing technical and community initiatives, these findings provide a structured basis for coordinated multi-stakeholder action to address a persistent and costly source of unreliability in biomedical research. Despite the challenges outlined above, this study represents an important step toward building the cross-sector alignment necessary to improve antibody validation standards and research reproducibility over time.

## Data Availability

All relevant data underlying the findings reported in this study are available within the manuscript and its Supporting Information files. No external data repository was used, and no DOIs, accession numbers or other persistent identifiers apply. Full quantitative results for both Delphi rounds are provided in S2 and S3 Tables, and complete anonymised qualitative feedback is provided in S7 and S8 Texts. Individual-level raw ratings are not provided in order to protect participant anonymity, consistent with the ethical approval for this study.

## Acknowledgements

We thank the following individuals for their expert participation as Delphi panellists. Their inclusion in this list acknowledges their contribution to the consensus process and does not necessarily imply endorsement of the final manuscript or its recommendations:

Alex Ball (Senior Scientist, GeneTex); Anita Bandrowski (CEO and Co-founder, SciCrunch); Andrew Bradbury (Founder and CSO, Specifica, an IQVIA business); Andrew Chalmers (CEO, CiteAb); Alejandra Clark (Managing Editor, The Company of Biologists); Katherine Crosby (Senior Director, Antibody Applications and Validation at Cell Signaling Technology); Brian Dickie (Chief Scientist, Motor Neurone Disease Association); Carly Dix (Associate Director, Biologics Engineering, AstraZeneca); Elizabeth M. Doherty (CHDI Management/CHDI Foundation); Andrew Economou (Senior Editor, Nature Protocols); Aled Edwards (Structural Genomics Consortium); Rachel Eyre (Programme Manager, NC3Rs); Simon Goodman (Former Visiting Professor, University of Turku); Richard A. Kahn (Emeritus Professor, Emory University); Ravindran Kumaran (Head of Collaborations, Abcam Ltd.); Carl Laflamme (Head of Antibody Characterisation Laboratory, SGC/Neuro, McGill University); Jeff Lee (COO, Proteintech); Catriona J. MacCallum (Director, Future of Scholarly Research, Wiley); Malcolm Macleod (Co-Director, Edinburgh Neuroscience, University of Edinburgh); Deborah Moshinsky (Director of Antibody Characterisation and Validation, Institute for Protein Innovation); Marcus Munafò (Deputy Vice Chancellor and Provost, University of Bath); Nicole K. Polinski (Director of Research Resources, The Michael J. Fox Foundation for Parkinson’s Research); Meghan Rego (Director of Research and Product Innovation, Addgene); Muge Sarper (Scientific Coordinator, The Chemical Probes Portal, The Institute of Cancer Research, London); Srikanth Subramanian (Principal Product Scientist at Cell Signaling Technology); Wei Mun Chan (Research Integrity Manager, eLife); Dalia Williams (Associate Publisher, F1000).

The remaining panellists did not provide explicit consent to be named within the required timeframe, consistent with the principle of anonymity in Delphi studies.

## Abbreviations

COM-B: Capabilities + Opportunities + Motivation Behaviour model
IPR: inter-percentile range
IPRAS: inter-percentile range adjusted for symmetry
IWGAV: International Working Group on Antibody Validation
MDAR: Materials Design Analysis Reporting
MRC: Medical Research Council
NC3Rs: National Centre for the Replacement, Refinement and Reduction of Animals in Research
OGA: Only Good Antibodies
RRID: Research Resource Identifier
YCharOS: Antibody Characterisation by Open Science

## References

1. Ayoubi R, Ryan J, Biddle MS, Alshafie W, Fotouhi M, Bolivar SG, et al. Scaling of an antibody validation procedure enables quantification of antibody performance in major research applications. Elife. 2023;12:RP91645. 10.7554/eLife.91645

2. Elliott S, Busse L, Bass MB, Lu H, Sarosi I, Sinclair AM, et al. Anti-Epo receptor antibodies do not predict Epo receptor expression. Blood. 2006;107(5):1892–5. 10.1182/blood-2005-10-4066

3. Berglund L, Björling E, Oksvold P, Fagerberg L, Asplund A, Al-Khalili Szigyarto C, et al. A genecentric Human Protein Atlas for expression profiles based on antibodies. Mol Cell Proteomics. 2008;7(10):2019–27. 10.1074/mcp.R800013-MCP200

4. Schrohl AS, Pedersen HC, Jensen SS, Nielsen SL, Brünner N. Human epidermal growth factor receptor 2 (HER2) immunoreactivity: specificity of three pharmacodiagnostic antibodies. Histopathology. 2011;59(5):975–83. 10.1111/j.1365-2559.2011.04034.x

5. Lukinavičius G, Lavogina D, Gönczy P, Johnsson K. Commercial Cdk1 antibodies recognize the centrosomal protein Cep152. Biotechniques. 2013;55(3):111–4. 10.2144/000114074

6. Baker M. Reproducibility crisis: Blame it on the antibodies. Nature. 2015;521(7552):274–6. 10.1038/521274a

7. Andersson S, Sundberg M, Pristovsek N, Ibrahim A, Jonsson P, Katona B, et al. Insufficient antibody validation challenges oestrogen receptor beta research. Nat Commun. 2017;8:15840. 10.1038/ncomms15840

8. Laflamme C, McKeever PM, Kumar R, Schwartz J, Kolahdouzan M, Chen CX, et al. Implementation of an antibody characterization procedure and application to the major ALS/FTD disease gene C9ORF72. Elife. 2019;8:e48363. 10.7554/eLife.48363

9. Virk HS, Rekas MZ, Biddle MS, Wright AKA, Sousa J, Weston CA, et al. Validation of antibodies for the specific detection of human TRPA1. Sci Rep. 2019;9(1):18500. 10.1038/s41598-019-55133-7

10. Kwon D. The antibodies don’t work! The race to rid labs of molecules that ruin experiments. Nature. 2024;635(8037):26–8. 10.1038/d41586-024-03590-0

11. Freedman LP, Cockburn IM, Simcoe TS. The economics of reproducibility in preclinical research. PLoS Biol. 2015;13(6):e1002165. 10.1371/journal.pbio.1002165

12. Kahn RA, Virk H, Laflamme C, Houston DW, Polinski NK, Meijers R, et al. Antibody characterization is critical to enhance reproducibility in biomedical research. Elife. 2024;13:e100211. 10.7554/eLife.100211

13. Kahn RA, Virk HS, McPherson PS. Heed a decade of calls for antibody validation. Nature. 2023;620(7974):492. 10.1038/d41586-023-02566-w

14. Biddle M, Cooper J, Blades K, Ruddy D, Krockow EM, Virk H. Drivers and ethical impacts of insufficient validation of antibodies in research. bioRxiv [Preprint]. 2026 Feb 19 [cited 2026 Feb 20]. Available from: 10.64898/2026.02.19.706766

15. Uhlen M, Bandrowski A, Carr S, Edwards A, Ellenberg J, Lundberg E, et al. A proposal for validation of antibodies. Nat Methods. 2016;13(10):823–7. 10.1038/nmeth.3995

16. Macleod M, Collings AM, Graf C, Kiermer V, Mellor D, Swaminathan S, et al. The MDAR (Materials Design Analysis Reporting) Framework for transparent reporting in the life sciences. Proc Natl Acad Sci U S A. 2021;118(17):e2103238118. 10.1073/pnas.2103238118

17. Bandrowski AE, Martone ME. RRIDs: a simple step toward improving reproducibility through rigor and transparency of experimental methods. Neuron. 2016;90(3):434–6. 10.1016/j.neuron.2016.04.030

18. Biddle M, Stylianou P, Rekas M, Wright A, Sousa J, Ruddy D, et al. Improving the integrity and reproducibility of research that uses antibodies: a technical, data sharing, behavioral and policy challenge. mAbs. 2024;16(1):2323706. 10.1080/19420862.2024.2323706

19. Fitch K, Bernstein SJ, Aguilar MD, Burnand B, LaCalle JR, Lazaro P, et al. The RAND/UCLA Appropriateness Method User’s Manual. Santa Monica, CA: RAND Corporation; 2001.

20. Dalkey N. An experimental study of group opinion. Futures. 1969;1(5):408–26. 10.1016/S0016-3287(69)80025-X

21. Fink A, Kosecoff J, Chassin M, Brook RH. Consensus methods: characteristics and guidelines for use. Am J Public Health. 1984;74(9):979–83. 10.2105/ajph.74.9.979

22. Virk H, Eyre R, Holmes A, Biddle M, Krockow E. Defining the role of antibodies in improving research reproducibility: NC3Rs and Only Good Antibodies community meeting report. London: NC3Rs; 2024 [cited 2026 Feb 8]. Available from: https://nc3rs.org.uk/sites/default/files/2024-07/NC3Rs-OGA%20Meeting%20report%20%E2%80%93%20Defining%20the%20role%20of%20antibodies%20in%20improving%20research%20reproducibility.pdf

