## Supplementary material for "A multi-stakeholder Delphi consensus study proposes actionable solutions to address antibody validation failures": S1 Table

**S1 Table. Detailed participant characteristics by employment sector.**

| **Characteristic** | **n** | **%** |
| --- | --- | --- |
| ***Academic Biomedical Research (n=7)*** | | |
| Academic Leadership | 2 | 28.6 |
| Senior Academic | 3 | 42.9 |
| Post-doctoral Researcher | 1 | 14.3 |
| Early Career Academic | 1 | 14.3 |
| ***Scientific Publishing (n=6)*** | | |
| Senior Editor | 2 | 33.3 |
| Editor | 2 | 33.3 |
| Editorial Director | 1 | 16.7 |
| Executive Role | 1 | 16.7 |
| ***Antibody Manufacturing – Commercial (n=7)*** | | |
| Director | 3 | 42.9 |
| Executive Role | 1 | 14.3 |
| Senior Scientist | 1 | 14.3 |
| Principal Scientist | 1 | 14.3 |
| Group Leader | 1 | 14.3 |
| ***Research Funding Organisations (n=4)*** | | |
| Director of Funding Programmes | 2 | 50.0 |
| Executive Role | 1 | 25.0 |
| Programme Manager | 1 | 25.0 |
| ***Industrial Biomedical Research (n=4)*** | | |
| Director | 1 | 25.0 |
| Executive Role | 1 | 25.0 |
| Principal Scientist | 1 | 25.0 |
| Group Leader | 1 | 25.0 |

**Note:** Industrial biomedical research includes n=2 solely in this sector, n=1 combined with antibody manufacturing, and n=1 combined with other sector. Five additional participants reported “other” employment fields including antibody informatics, infrastructure provider, charity sector (previously industry), retired (40 years antibody manufacturing experience in academic and industrial R&D), and biotech antibody CRO founder. Participants reporting work across multiple sectors may appear in more than one category; primary sector classifications used in the main text total 32
