## Supplementary material for "A multi-stakeholder Delphi consensus study proposes actionable solutions to address antibody validation failures": S2 Table

**S2 Table. Full quantitative results for Round 1 (n=32 participants, 23 items).**

| **Item** | **Description** | **Med Eff** | **Med Feas** | **IPR Eff** | **IPRAS Eff** | **IPR Feas** | **IPRAS Feas** | **Rating Eff** | **Rating Feas** | **Disagree Eff** | **Disagree Feas** | **Number of ratings Eff (%)** | **Number of ratings Feas (%)** |
| --- | --- | --- | --- | --- | --- | --- | --- | --- | --- | --- | --- | --- | --- |
| ***Journal and Publisher Policies*** | | | | | | | | | | | | | |
| **R1** | Authors include RRIDs for each antibody used | 7.0 | 8.0 | 1 | 6.10 | 2 | 6.85 | Effective | Feasible | No | No | 96.9 | 96.9 |
| **R2** | Authors provide sufficient metadata for unambiguous antibody identification | 8.0 | 8.0 | 2 | 6.85 | 3 | 6.10 | Effective | Feasible | No | No | 100.0 | 100.0 |
| **R3** | Authors report antibody concentration or dilution | 8.0 | 8.0 | 1 | 6.10 | 1 | 7.60 | Effective | Feasible | No | No | 100.0 | 100.0 |
| **R4** | Journals require validation data in open-access repository | 8.0 | 5.0 | 1 | 6.10 | 3 | 3.10 | Effective | Uncertain | No | No | 96.9 | 96.9 |
| **R5** | Journals introduce tiered scoring/transparency badge | 5.0 | 4.0 | 2 | 2.35 | 2 | 3.85 | Uncertain | Uncertain | No | No | 93.8 | 93.8 |
| **R6** | Publishers train reviewers/editors on validation assessment | 6.5 | 5.0 | 3 | 4.60 | 4 | 2.35 | Uncertain | Uncertain | No | Yes | 100.0 | 100.0 |
| **R7** | Publishers support automated validation flagging tools | 8.0 | 5.0 | 1 | 6.10 | 2 | 3.85 | Effective | Uncertain | No | No | 96.9 | 90.6 |
| **R8** | Authors report validation using IWGAV framework | 7.0 | 6.0 | 1 | 5.35 | 3 | 3.10 | Effective | Uncertain | No | No | 93.8 | 90.6 |
| **R9** | Authors provide genetic validation for critical antibodies | 8.0 | 5.0 | 2 | 6.85 | 3 | 3.10 | Effective | Uncertain | No | No | 93.8 | 90.6 |
| ***Research Funder Actions*** | | | | | | | | | | | | | |
| **R10** | Funders include validation plan section in application form | 7.5 | 8.0 | 1 | 6.10 | 2 | 6.85 | Effective | Feasible | No | No | 93.8 | 93.8 |
| **R11** | Applicants include specific budget line for antibody validation | 8.0 | 7.0 | 1 | 6.10 | 2 | 5.35 | Effective | Feasible | No | No | 96.9 | 90.6 |
| **R12** | Funders create/expand targeted tool development and validation schemes | 8.0 | 7.0 | 1 | 6.10 | 2 | 3.85 | Effective | Feasible | No | No | 100.0 | 90.6 |
| **R13** | Funders signal importance of antibody validation in applicant guidance | 7.0 | 7.0 | 1 | 4.60 | 1 | 6.10 | Effective | Feasible | No | No | 100.0 | 100.0 |
| **R14** | Funders encourage deposition of validation data in open repositories | 7.0 | 7.0 | 1 | 6.10 | 2 | 5.35 | Effective | Feasible | No | No | 90.6 | 90.6 |
| **R15** | Funders formally endorse community-developed reporting standards | 7.0 | 7.0 | 2 | 5.35 | 0 | 5.35 | Effective | Feasible | No | No | 96.9 | 93.8 |
| **R16** | Funders directly engage manufacturers on transparency | 7.0 | 5.0 | 2 | 5.35 | 4 | 2.35 | Effective | Uncertain | No | **Yes** | 100.0 | 90.6 |
| **R17** | Funders co-fund independent benchmarking initiatives | 7.5 | 6.0 | 1 | 6.10 | 2 | 3.85 | Effective | Uncertain | No | No | 93.8 | 90.6 |
| **R18** | Funders promote industry participation in benchmarking | 7.0 | 6.0 | 2 | 5.35 | 2 | 3.85 | Effective | Uncertain | No | No | 81.3 | 78.1 |
| ***Institutional and Educational Actions*** | | | | | | | | | | | | | |
| **R19** | Training in antibody validation in relevant bioscience courses | 8.0 | 7.0 | 2 | 6.85 | 1 | 6.10 | Effective | Feasible | No | No | 100.0 | 100.0 |
| **R20** | Incorporate antibody validation into research integrity/ethics frameworks | 7.0 | 7.0 | 1 | 6.10 | 1 | 6.10 | Effective | Feasible | No | No | 100.0 | 96.9 |
| **R21** | Recognise and support local champions in antibody validation | 7.0 | 6.5 | 1 | 6.10 | 2 | 3.85 | Effective | Uncertain | No | No | 100.0 | 100.0 |
| ***Cross-Stakeholder Coordination*** | | | | | | | | | | | | | |
| **R22** | Develop shared roadmap for improving antibody validation by 2030 | 7.0 | 7.0 | 1 | 6.10 | 2 | 3.85 | Effective | Feasible | No | No | 100.0 | 96.9 |
| **R23** | Develop coordinated shared infrastructure for validation data | 8.0 | 6.0 | 1 | 6.10 | 2 | 3.85 | Effective | Uncertain | No | No | 100.0 | 96.9 |

**Note:** Med = Median; Eff = Effectiveness; Feas = Feasibility; IPR = Inter-Percentile Range; IPRAS = IPR Adjusted for Symmetry. Consensus classification: Effective/Feasible (median 7–9), Uncertain (median 4–6), Ineffective/Infeasible (median 1–3). Disagreement = IPR > IPRAS (shown in bold). Number of ratings = percentage of the panel providing a rating for the item. Items achieving consensus as both effective and feasible in Round 1 (R1, R2, R10–R15, R19, R20, R22) were accepted as recommendations and not re-rated in Round 2.
