## Supplementary material for "A multi-stakeholder Delphi consensus study proposes actionable solutions to address antibody validation failures": S3 Table

**S3 Table. Full quantitative results for Round 2 (n=30 participants, 22 items: 12 re-rated from Round 1, 10 new).**

| **Item** | **Description** | **Med Eff** | **Med Feas** | **IPR Eff** | **IPRAS Eff** | **IPR Feas** | **IPRAS Feas** | **Rating Eff** | **Rating Feas** | **Disagree Eff** | **Disagree Feas** | **Number of Ratings Eff (%)** | **Number of Ratings Fea (%)** |
| --- | --- | --- | --- | --- | --- | --- | --- | --- | --- | --- | --- | --- | --- |
| ***Re-rated Items from Round 1*** | | | | | | | | | | | | | |
| **R3** | Authors report antibody dilution and concentration | 8.0 | 8.0 | 0 | 6.85 | 0 | 6.85 | Effective | Feasible | No | No | 100.0 | 96.7 |
| **R4** | Journals require validation data in open repositories | 8.0 | 5.0 | 1 | 6.10 | 2 | 2.35 | Effective | Uncertain | No | No | 100.0 | 96.7 |
| **R5** | Journals use transparency badges or tiered scoring | 4.0 | 3.0 | 2 | 3.85 | 0 | 5.35 | Uncertain | Infeasible | No | No | 96.7 | 96.7 |
| **R6** | Publishers train reviewers/editors on validation | 7.0 | 5.0 | 2 | 5.35 | 3 | 3.10 | Effective | Uncertain | No | No | 100.0 | 100.0 |
| **R7** | Publishers support automated validation flagging tools | 8.0 | 5.0 | 1 | 6.10 | 1 | 3.10 | Effective | Uncertain | No | No | 96.7 | 96.7 |
| **R8** | Authors report validation using IWGAV framework | 7.0 | 6.0 | 0 | 5.35 | 2 | 3.85 | Effective | Uncertain | No | No | 96.7 | 96.7 |
| **R9** | Authors provide genetic validation for critical antibodies | 8.0 | 5.0 | 0 | 6.85 | 2 | 3.85 | Effective | Uncertain | No | No | 93.3 | 90.0 |
| **R16** | Funders engage manufacturers on transparency | 7.0 | 5.0 | 1 | 4.60 | 2 | 3.85 | Effective | Uncertain | No | No | 100.0 | 96.7 |
| **R17** | Funders co-fund independent benchmarking initiatives | 8.0 | 6.0 | 1 | 6.10 | 2 | 3.85 | Effective | Uncertain | No | No | 96.7 | 93.3 |
| **R18** | Funders promote industry benchmarking participation | 7.0 | 6.0 | 1 | 4.60 | 2 | 3.85 | Effective | Uncertain | No | No | 93.3 | 90.0 |
| **R21** | Institutions support local validation champions | 7.0 | 7.0 | 1 | 6.10 | 1 | 4.60 | Effective | Feasible | No | No | 100.0 | 100.0 |
| **R23** | Develop shared validation data infrastructure | 8.0 | 6.0 | 0 | 6.85 | 1 | 3.10 | Effective | Uncertain | No | No | 96.7 | 96.7 |
| ***New Items Added in Round 2*** | | | | | | | | | | | | | |
| **A1** | Journals establish clear validation standards | 7.5 | 7.0 | 1 | 6.10 | 2 | 5.35 | Effective | Feasible | No | No | 100.0 | 100.0 |
| **A2** | Validation protocols included in manuscripts/supplements | 8.0 | 6.0 | 1 | 6.10 | 2 | 4.60 | Effective | Uncertain | No | No | 100.0 | 100.0 |
| **A3** | Journals appoint specialist reproducibility editors | 7.0 | 5.5 | 2 | 5.35 | 3 | 3.10 | Effective | Uncertain | No | No | 96.7 | 93.3 |
| **A4** | Funders maintain list of validating manufacturers | 6.0 | 5.0 | 2 | 3.85 | 2 | 2.35 | Uncertain | Uncertain | No | No | 96.7 | 93.3 |
| **A5** | Funders mandate recombinant antibody use | 6.0 | 5.0 | 2 | 3.85 | 4 | 2.35 | Uncertain | Uncertain | No | **Yes** | 86.7 | 86.7 |
| **A6** | Funded antibodies must be recombinant and publicly available | 7.0 | 6.0 | 1 | 6.10 | 3 | 3.10 | Effective | Uncertain | No | No | 90.0 | 83.3 |
| **A7** | Manufacturers assign RRIDs at source | 8.0 | 7.0 | 1 | 6.10 | 1 | 6.10 | Effective | Feasible | No | No | 93.3 | 80.0 |
| **A8** | Manufacturers perform standard validation experiments | 8.0 | 6.5 | 0 | 6.85 | 2 | 3.85 | Effective | Uncertain | No | No | 100.0 | 86.7 |
| **A9** | Manufacturers shift to recombinant antibody production | 7.0 | 5.0 | 1 | 6.10 | 3 | 3.10 | Effective | Uncertain | No | No | 86.7 | 73.3 |
| **A10** | Establish learned society for antibody validation | 7.0 | 5.0 | 2 | 5.35 | 2 | 3.85 | Effective | Uncertain | No | No | 100.0 | 93.3 |

**Note:** Med = Median; Eff = Effectiveness; Feas = Feasibility; IPR = Inter-Percentile Range; IPRAS = IPR Adjusted for Symmetry. Consensus classification: Effective/Feasible (median 7–9), Uncertain (median 4–6), Ineffective/Infeasible (median 1–3). Disagreement = IPR > IPRAS (shown in bold). Number of ratings = percentage of the panel providing a rating for the item. Items R3, R21, A1, and A7 achieved consensus as both effective and feasible in Round 2. Item R5 shifted to infeasible (median feasibility declined from 4.0 to 3.0). Item A5 was the only item with panel disagreement (on the feasibility dimension).
