## Supplementary material for "A multi-stakeholder Delphi consensus study proposes actionable solutions to address antibody validation failures": S4 Table

**S4 Table. Summary of items grouped by stakeholder group and rating.**

| **Item** | **Description** | **Median Effectiveness** | **Median Feasibility** | **Rating** |
| --- | --- | --- | --- | --- |
| ***Journal and Publisher Items*** | | | | |
| **R1** | Authors should include the Research Resource Identifier (RRID) for each antibody used, where available. | 7.0 | 8.0 | Effective, Feasible |
| **R2** | Authors should provide sufficient metadata to allow unambiguous identification of each antibody. This includes: Clone ID (for monoclonal antibodies), Catalogue number and lot number (for polyclonal antibodies), Vendor or source. | 8.0 | 8.0 | Effective, Feasible |
| **R3** | Authors should report the dilution ratio at which each antibody is used. In addition, authors should state the antibody protein concentration where possible. Examples include dilution ratios such as 1:1000 or 1:500, or concentrations expressed in µg/mL. | 8.0 | 8.0 | Effective, Feasible |
| **A1** | Journals should have a clearly stated standard for antibody validation and reporting. | 7.5 | 7.0 | Effective, Feasible |
| **R4** | Journals should require authors to deposit antibody validation data in an open-access repository, linked to a persistent identifier (e.g., RRID, DOI). Repositories might include: Zenodo, institutional repositories, manufacturer portals, or The Antibody Registry (linked to Biomed Resource Watch). | 8.0 | 5.0 | Effective, Uncertain Feasibility |
| **R6** | Publishers should provide training resources for reviewers and editors to help them evaluate antibody validation information in submitted manuscripts. | 7.0 | 5.0 | Effective, Uncertain Feasibility |
| **R7** | Publishers and other stakeholders should support the development and use of automated tools to flag potential antibody validation issues in submitted manuscripts – e.g., for antibodies that have already been withdrawn from the market because of performance issues. | 8.0 | 5.0 | Effective, Uncertain Feasibility |
| **R8** | As part of the submission process, authors should describe how antibody specificity was assessed, making reference to the International Working Group on Antibody Validation (IWGAV) framework. A brief paragraph, checklist, or table should indicate which of the IWGAV pillars were used, if any (e.g., genetic, orthogonal, independent antibody, tagged protein, IP-MS). | 7.0 | 6.0 | Effective, Uncertain Feasibility |
| **R9** | Where antibodies are critical to key experimental conclusions, where technically feasible, authors should either: present validation data based on a genetic strategy (e.g., knockout or knockdown), or cite previously published genetic validation data. | 8.0 | 5.0 | Effective, Uncertain Feasibility |
| **A2** | Detailed Protocols and Validation data should be included in the manuscript or supplemental data. | 8.0 | 6.0 | Effective, Uncertain Feasibility |
| **A3** | Journals should appoint a specialist editor that focuses on reproducibility. | 7.0 | 5.5 | Effective, Uncertain Feasibility |
| **R5** | Journals should introduce a tiered scoring system or transparency badge to recognise papers that meet higher standards of antibody reporting and validation. | 4.0 | 3.0 | Uncertain Effectiveness, Unfeasible; rejected |
| ***Research Funder Items*** | | | | |
| **R10** | Funders should include a section in the application form where applicants must detail the steps they will take to validate the antibodies used. | 7.5 | 8.0 | Effective, Feasible |
| **R11** | Applicants to biomedical funding schemes should include a specific line item within their budget for resources to conduct robust antibody validation (if relevant to their proposal). | 8.0 | 7.0 | Effective, Feasible |
| **R12** | Funders should create or expand targeted schemes to support the development and validation of critical antibody-based research tools. These could include, for example: Characterisation of under-studied antibodies, Generation of KO/KD cell lines, Cross-platform benchmarking of antibody performance, Replacement of animal-derived antibodies. | 8.0 | 7.0 | Effective, Feasible |
| **R13** | Funders should share information within the applicant guidance that antibody performance is an important limitation in many experimental methods. Applicants should be encouraged to address this in methodological or reproducibility sections, as this will be part of how these sections are evaluated by reviewers and panels. | 7.0 | 7.0 | Effective, Feasible |
| **R14** | Funders should encourage grantees to deposit antibody validation data in open-access repositories, ideally linked to RRIDs or registry entries. This expectation could be embedded in end-of-grant reporting or incentivised through visibility or reproducibility initiatives. | 7.0 | 7.0 | Effective, Feasible |
| **R15** | Funders should formally endorse community-developed reporting standards that promote antibody transparency and validation (e.g., IWGAV, MDAR). Such endorsements can help reinforce aligned expectations across funders and journals. | 7.0 | 7.0 | Effective, Feasible |
| **R16** | Funders should directly engage with manufacturers to encourage them to: Publish antibody validation datasets, Adopt consistent metadata standards (e.g., RRID, clone ID, lot number). | 7.0 | 5.0 | Effective, Uncertain Feasibility |
| **R17** | Funders should collaborate with independent benchmarking initiatives such as YCharOS or equivalent open-science consortia. They should directly co-fund the validation of antibodies against key targets to create suites of validated tools that can be used by researchers to improve data quality and thereby increase return on investment for the funder. | 8.0 | 6.0 | Effective, Uncertain Feasibility |
| **R18** | Funders should promote industry participation in independent benchmarking initiatives by asking applicants to consider working with initiatives such as YCharOS or equivalent open-science consortia as part of their work. This could be achieved either by including the recommendation in the guidance notes or flagging to applicants at the panel review stage. | 7.0 | 6.0 | Effective, Uncertain Feasibility |
| **A6** | Funders should mandate that if funds are used to develop an antibody as part of a research project, the reagent must be a recombinant antibody and must be made available to the wider community. | 7.0 | 6.0 | Effective, Uncertain Feasibility |
| **A4** | Funders should maintain and provide researchers with a list of manufacturers who validate their antibodies. | 6.0 | 5.0 | Uncertain Both |
| **A5** | Funders should mandate the use of recombinant antibodies (or at least monoclonals) when available. Justification should be required if a polyclonal is used. | 6.0 | 5.0* | Uncertain Both |
| ***Institutional and Educational Items*** | | | | |
| **R19** | Training in antibody validation should be offered in relevant bioscience courses and programmes. This could include: Undergraduate education (e.g., in immunology, neuroscience, or molecular biology programmes), Postgraduate or doctoral training (e.g., through doctoral training centres), Postdoctoral or continuing professional development. Training should be field-appropriate, not universally required across all domains. | 8.0 | 7.0 | Effective, Feasible |
| **R20** | Universities and research institutions should incorporate antibody validation expectations into research integrity and ethics review frameworks. This may include: Asking researchers to consider antibody validation as part of research ethics approval processes, Providing guidance or templates for documenting validation plans. | 7.0 | 7.0 | Effective, Feasible |
| **R21** | Universities, research institutions and learned societies should recognise and support the Antibody Champions Scheme. This UK-based initiative, co-funded by the Medical Research Council (MRC) and the National Centre for the replacement, refinement and reduction of animals in research (NC3Rs), will be launched in 2026. It will be a voluntary scheme of local champions or experts in antibody validation. These individuals will serve as: Points of contact for advice on antibody selection, validation design, and data interpretation; Trainers or facilitators in antibody validation best practice workshops or research methods courses; Advocates for antibody good practice within departments or core facilities. Support could include protected time, acknowledgement in performance reviews, or funding for training and outreach. | 7.0 | 7.0 | Effective, Feasible |
| **A10** | A learned society for antibody validation should be established. | 7.0 | 5.0 | Effective, Uncertain Feasibility |
| ***Manufacturer Items*** | | | | |
| **A7** | Antibody manufacturers should be encouraged to assign RRIDs to their products at source. | 8.0 | 7.0 | Effective, Feasible |
| **A8** | Manufacturers should perform a standard set of validation experiments on their reagent antibodies and make the data available. | 8.0 | 6.5 | Effective, Uncertain Feasibility |
| **A9** | Manufacturers should shift production to all recombinant antibodies. | 7.0 | 5.0 | Effective, Uncertain Feasibility |
| ***Cross-Stakeholder Items*** | | | | |
| **R22** | Stakeholders should work together to develop a shared roadmap for improving antibody validation practices by 2030. This would outline: Stakeholder-specific actions and responsibilities, Target timelines and milestones, Mechanisms for accountability and feedback. | 7.0 | 7.0 | Effective, Feasible |
| **R23** | A coordinated, shared infrastructure should be developed (or expanded) to: Aggregate antibody validation data, Provide standardised formats and tools for sharing, Enable journals, funders, and institutions to access validation status or red flags. This may involve expanding existing platforms such as YCharOS, or building interoperability across initiatives like RRID, Benchsci, CiteAb, and Antibody Registry. | 8.0 | 6.0 | Effective, Uncertain Feasibility |

**Note:** Median effectiveness and feasibility were rated on a 1–9 scale. *Panel disagreement on feasibility (IPR = 4 > IPRAS = 2.35).
