## Supplementary material for "A multi-stakeholder Delphi consensus study proposes actionable solutions to address antibody validation failures": S1 Text

**S1 TEXT. DETAILED OVERVIEW OF CONSENSUS RECOMMENDATIONS**

**1.1 JOURNAL AND PUBLISHER RECOMMENDATIONS**

**R1: Authors should include RRIDs where available**

**(Effectiveness: 7.0, Feasibility: 8.0, accepted in Round 1)**

**The Recommendation**

Authors should include the Research Resource Identifier (RRID) for each antibody used, where available.

**Panel Assessment**

This recommendation achieved consensus as both effective (median 7.0) and feasible (median 8.0), without disagreement, in Round 1. The panel agreed that RRID reporting would support reproducibility and could realistically be implemented by 2030.

**Supporting Rationale**

Panellists recognised RRIDs as a practical step toward improved reagent identification, with one noting that "effectiveness and feasibility has been demonstrated empirically." The qualifier "where available" was seen as appropriately pragmatic. One participant suggested that "antibody producers could be encouraged to assign RRIDs to their products at source. Down to individual batches if possible. That could complete the identification chain for the reagents being used."

**Implementation Considerations**

The panel identified several technical limitations of the current RRID system that should be addressed to maximize effectiveness. The most significant gap is that RRIDs do not currently capture lot-to-lot variation, which is particularly problematic for polyclonal antibodies that "remain the most used antibody product." This suggests that RRID implementation should be paired with lot number reporting (as recommended in R2) rather than viewed as a standalone solution.

Multiple participants noted that the same antibody clone sold by different vendors receives different RRIDs, which can inadvertently lead researchers to believe they are using independent antibodies for validation when they are not. This indicates a need for RRID system enhancements to cross-reference identical clones across suppliers, or for complementary reporting of clone names/IDs alongside RRIDs.

Participants suggested that upstream assignment of RRIDs—"antibody producers could be encouraged to assign RRIDs to their products at source, down to individual batches if possible"—would streamline adoption and "complete the identification chain." This points to a coordination opportunity with manufacturers (see also A7).

Implementation barriers included limited researcher awareness of how RRIDs are generated and how to use them, suggesting a need for educational resources embedded in journal author guidelines. While journal enforcement through automated screening was seen as feasible for larger publishers, smaller or scholar-led publishers may require shared infrastructure or tools to check RRID compliance cost-effectively. One participant noted that "author behaviour is shaped by incentive structures that currently do not reward efforts toward reproducibility," indicating that journal policies may be more effective when aligned with funder expectations for RRID reporting.

**R2: Authors should provide complete antibody metadata**

**Effectiveness: 8.0 | Feasibility: 8.0 | Status: Consensus - Accepted in Round 1**

**The Recommendation:**

Authors should provide sufficient metadata to allow unambiguous identification of each antibody. This includes:

- Clone ID (for monoclonal antibodies)
- Catalogue number and lot number (for polyclonal antibodies)
- Vendor or source

**Panel Assessment:**

This recommendation achieved strong ratings on both effectiveness (median 8.0) and feasibility (median 8.0). The panel judged that providing complete antibody metadata would meaningfully improve research reproducibility and that this standard could realistically be adopted by journals by 2030. No statistical disagreement was detected, indicating consistent support across the diverse stakeholder panel.

**Supporting Rationale:**

Panellists viewed complete metadata reporting as foundational to antibody identification and a natural complement to RRID reporting (R1). One participant noted that "metadata would be best used in combination with RRIDs," highlighting how these recommendations work synergistically. The straightforward nature of this requirement—asking authors to report information they should already have when purchasing and using antibodies—contributed to high feasibility ratings. Participants emphasized that this addresses a basic reproducibility need: readers must be able to identify precisely which antibody was used to attempt replication or interpret results.

However, panellists also noted that effectiveness depends on implementation mechanisms. As one participant observed, "This would require buy-in and enforcement from journals and vendors," underscoring that journal policies alone are insufficient without verification processes and vendor cooperation in providing clear, consistent metadata on product packaging and datasheets.

**Implementation Considerations:**

The primary implementation challenge identified was enforcement. While adding metadata requirements to author guidelines is straightforward, several participants questioned how journals would verify compliance, particularly for journals with limited editorial resources. Panellists noted that "authors don't know which items to add without a clear unambiguous guideline and with a guideline they often partially comply," suggesting that structured templates or data entry fields may be more effective than narrative instructions.

Space constraints in manuscripts were identified as a practical barrier, pointing to the need for publisher accommodation through supplementary materials sections or structured metadata fields. Panellists also noted complications when companies rename clones or when the same antibody is sold by multiple vendors under different catalogue numbers, indicating this works best when combined with RRIDs (R1) or persistent clone IDs.

A cultural challenge was identified around data capture practices: "There would need to be a change of culture in organisations at the stage of data capture, as this might be overlooked and result in inaccurate reporting." One participant noted that "some vendors do not provide unambiguous production lot numbers," highlighting a need for manufacturer engagement. Panelists also highlighted that standardisation across journals would strengthen effectiveness, suggesting value in coordinating metadata standards through existing journal initiatives or discipline-specific reporting standards.

**R3: Authors should report antibody dilution and concentration**

**Effectiveness: 8.0 | Feasibility: 8.0 | Status: Consensus - Accepted in Round 2 (refined from Round 1)**

**The Recommendation:**

Authors should report the dilution ratio at which each antibody is used. In addition, authors should state the antibody protein concentration where possible. Examples include dilution ratios such as 1:1000 or 1:500, or concentrations expressed in μg/mL.

**Panel Assessment:**

This recommendation maintained strong consensus across both rounds (Round 1: 8.0/8.0; Round 2: 8.0/8.0), with wording refined between rounds to prioritize dilution ratio as the minimum reportable information while encouraging protein concentration reporting where available. The panel judged this information essential for reproducibility, as antibody performance is concentration-dependent.

**Supporting Rationale:**

Panelists emphasized that absolute protein concentration is scientifically preferable to dilution ratios alone, as identical dilution ratios can yield different working concentrations depending on stock concentration. As one participant stated: "Protein concentration should be reported as much as possible as dilution ratios alone may not be sufficient information for others to reproduce an experiment accurately." However, participants recognized that protein concentration is not always available from manufacturers, particularly for polyclonal antibodies. The refined wording establishing dilution ratio as the minimum standard while encouraging concentration reporting where available addressed this practical constraint.

**Implementation Considerations:**

The primary challenge identified was the variability in what concentration information manufacturers provide. While some participants argued that responsible vendors should always report protein concentration, others noted this remains inconsistent in practice, particularly for polyclonal antibodies where concentration may be variable or unknown. Panellists also raised practical issues around serial dilutions performed in laboratories (e.g., 1:1 with glycerol for storage) that are "rarely included in calculations/reporting by lab staff" and whether total dilution from the commercially obtained source should be reported.

Participants noted that concentration reporting in µg/mL is particularly important for monoclonals where "different sources will have different concentrations and dilutions could be confusing," while dilution ratios may be acceptable for polyclonals. The key implementation question is whether journals can verify compliance without placing excessive burden on editors with limited resources, and whether coordination across journals would strengthen adoption.

**A1: Journals should have clearly stated validation standards**

**(Effectiveness: 7.5, Feasibility: 7.0, introduced and accepted in Round 2)**

**The Recommendation**

Journals should have a clearly stated standard for antibody validation and reporting.

**Panel Assessment**

This recommendation achieved consensus as both effective (median 7.5) and feasible (median 7.0) without disagreement. The panel agreed that establishing clear journal standards would meaningfully improve antibody practices and could realistically be implemented by 2030.

**Supporting Rationale**

Panelists emphasized that clear standards would provide much-needed clarity for authors and signal journals' commitment to reproducibility. One participant noted that "having a clearly stated standard even if the policy is not fully aligned with the gold standard is a start and a means for journals to signal they are on the right trajectory." The suggestion of tiered approaches—similar to the evolution of data sharing policies from "encourages" to "expects" or "mandates"—was seen as providing a feasible transitional path.

**Implementation Considerations**

The central tension identified by the panel concerned the relationship between individual journal standards and broader community consensus. Multiple participants emphasized that journal standards "would only be effective if there were community standards already in place or consensus across most journals." The challenge of achieving uniform standards across journals was repeatedly noted, with concerns that differing standards "would complexify submission processes" and create risks that "authors would publish elsewhere" if requirements were perceived as too burdensome.

Importantly, external standards frameworks already exist which journals could reference. The International Working Group for Antibody Validation (IWGAV) five-pillar framework provides established guidance for context-specific validation approaches (genetic strategies, independent antibodies, tagged expression, immunocapture mass spectrometry, and antibody-independent orthogonal methods). The YCharOS platform offers complementary standardized characterization data on antibody performance across major research applications (Western blot, immunoprecipitation, immunofluorescence), which can inform—though not replace—context-specific validation requirements.

Participants suggested that coordination bodies like "YCharOS / OGA acted as an interface to set up an agreed standard reporting template, and aligned it with the major publishers." The practical challenge of validation diversity was raised: "a checklist may not be feasible depending on the specific target(s)" and "there are multiple hallmarks of validation; no one validation method is better than another, but the cumulative effect of several validation methods that is most powerful." One participant noted the temporal challenge that standards "would require a long lead time to make researchers aware of standards prior to starting work."

**1.2 Funder Recommendations**

**R10: Funders should require validation plans in applications**

**(Effectiveness: 7.5, Feasibility: 8.0, accepted in Round 1)**

**The Recommendation**

Funders should include a section in the application form where applicants must detail the steps they will take to validate the antibodies used.

**Panel Assessment**

This recommendation achieved consensus as both effective (median 7.5) and highly feasible (median 8.0) without disagreement. The panel agreed that requiring validation plans in grant applications would improve antibody practices and could realistically be implemented by 2030.

**Supporting Rationale**

Panelists viewed this as a straightforward and highly feasible intervention: "Easy for funders to implement" and could be "implemented in application templates across funding organizations." One participant noted that NIH already has a related requirement through its "key biological resources supplemental file," demonstrating that such policies exist. The act of requiring applicants to articulate validation plans was seen as valuable even if not strictly enforced: "it will likely become a copy-paste scenario where applicants pay lip service to the principles of antibody validation but in practice skip this step to save on time/budget. The benefit of this would be solely in serving as a reminder and soft encouragement."

**Implementation Considerations**

The central implementation challenge identified was the risk of this becoming "another box ticking exercise for the applicant, which is filled with little care and using generic text each time." Multiple participants emphasized that effectiveness depends entirely on how the information is used in grant evaluation: "Effectiveness depends on how essential the information is (i.e. will the project be scored based on this information, or is it an optional extra?)."

Panelists questioned whether funders and review panels have the capacity to meaningfully evaluate validation plans: "I marked 6 for feasibility as I am not sure how easy it would be for funders and grant committees to evaluate this extra information." The suggestion was made that "some of this type of assessment could be done in conjunction with AI tools."

A fundamental challenge was identified around timing: "Would you know this information prior to starting the study?" and "Detailing the steps is different from taking them." This points to a tension between prospective planning (at application stage) and actual validation conduct (during the funded research). One participant noted: "The main issue here is follow through. In many cases, once the grant is funded there is no over-site to ensure follow through. If more grants required meeting milestones for the subsequent years it would be more effective."

Panelists also questioned whether applicants would be expected to validate all antibodies themselves or could rely on existing data: "Will researchers always validate their own abs or utilize other sources (suppliers websites, previous publications)?" This relates to the broader question of what constitutes adequate validation evidence in the context of a proposed study.

**R11: Funders should include antibody validation budget line**

**(Effectiveness: 8.0, Feasibility: 7.0, accepted in Round 1)**

**The Recommendation**

A budget for antibody validation should also be costed for in the application. Applicants to biomedical funding schemes should include a specific line item within their budget for resources to conduct robust antibody validation (if relevant to their proposal).

**Panel Assessment**

This recommendation achieved consensus as both effective (median 8.0) and feasible (median 7.0) without disagreement. The panel judged that explicit budget allocation for validation would meaningfully improve antibody practices and could realistically be implemented by funders by 2030.

**Supporting Rationale**

Panelists recognized that dedicated budget allocation addresses a fundamental barrier to validation: the perception that validation is an unfunded add-on. As one participant noted: "Antibody validation is expensive but setting aside a budget ahead of time would help researchers to plan ahead." Another emphasized: "If specific funding is provided authors will complete the required experiments." The principle of "you get what you pay for" was invoked multiple times, with one participant arguing that validation budget allocation would "likely increase [funders'] ROI" by improving research quality.

The importance of this recommendation was seen as proportional to the criticality of the antibody to the research: "the importance and utility of having it rises very sharply depending on how critical the antibody is to the research project in question."

**Implementation Considerations**

The primary challenge identified was budget constraint in competitive funding environments. Panelists noted tensions between validation costs and research scope: "research proposals typically employ multiple antibodies in a study, sometimes upwards of 5-10. Would the applicant be required to validate all antibodies prior to use? Would they need to validate them in their specific experimental cell/tissue system? This would result in a pretty significant budget increase which would come at the cost of reducing the scope of the hypothesis-driven research study or cutting down the number of applications a funder can fund for a program."

The feasibility concern centered on whether additional validation budget would come from existing grant budgets or represent new funding. As one participant noted: "research is expensive and the funding situation (at least in the US) is precarious right now," and "funding levels are tight and getting money assigned to validate reagents will be difficult."

Panelists also raised questions about whether researchers would need to budget for validating every antibody themselves or could rely on existing validation data from manufacturers or published literature: "This assumes researchers will have to validate every antibody without using other sources." The recommendation does not specify the extent of validation required, leaving open whether applicants should budget for comprehensive in-house validation or verification experiments.

An implementation question was raised about accountability: "Unclear whether these funds would actually be aligned to this," suggesting that even when budgeted, validation funds might be reallocated to other purposes during project execution.

### R12: Funders should support antibody tool development schemes

### (Effectiveness: 8.0, Feasibility: 7.0, accepted in Round 1)

**The Recommendation**

Funders should create or expand targeted schemes to support the development and validation of critical antibody-based research tools. These could include, for example: Characterisation of under-studied antibodies, Generation of KO/KD cell lines, Cross-platform benchmarking of antibody performance, Replacement of animal-derived antibodies.

**Panel Assessment**

This recommendation achieved consensus as both effective (median 8.0) and feasible (median 7.0) without disagreement. The panel agreed that dedicated funding schemes for antibody tool development would meaningfully improve research infrastructure and could realistically be implemented by 2030.

**Supporting Rationale**

Panelists strongly endorsed the principle of funders supporting antibody tool infrastructure. One participant emphasized urgency: "There is no reason why these steps should not be implemented asap. In a related issue, there is currently no repository for cell lines KO'd for specific targets and such a resource would be a valuable resource for several uses." The creation of shared resources—validated antibodies, knockout cell lines, benchmarking data—was recognized as addressing infrastructure gaps that individual researchers cannot fill: "Validated recombinant antibodies where sequences are publicly available would be the best solutions."

However, panelists also noted that tool creation alone is insufficient without promotion and education: "There needs to be a larger initiative by funders to not only create validated tools but also to promote them. Currently researchers want to use cited antibodies even if they are poor. They are resistant to change even if a new well validated tool comes out. In addition to providing well validated antibodies there needs to be a strong initiative to educate researchers about the new reagents and push them to use them."

**Implementation Considerations**

The primary implementation challenge identified was scale relative to need. Multiple participants noted: "This would potentially help some of the targets, but there are 22000 proteins" and "The vast number of antibodies available would make it challenging to create the resources described above." This points to the need for strategic prioritization of which targets or antibodies to support, as comprehensive coverage is infeasible.

Funding constraints were raised repeatedly, particularly for disease-specific funders: "This may be a challenge in the orphan disease arena, where funding is limited" and "could be hard to justify/implement as it is not something that is easy to explain or attractive to patients/donors for fundraising efforts." One participant questioned whether resources might be better allocated: "Difficult to know if this is feasible - which depends on the funders have sufficient money to implement these schemes. It's possible that the money might be better spent on education?"

The case for investment needs strengthening: "It would depend on a cost-benefit analysis. Otherwise funders might be very reluctant 'what for? There are loads of antibodies (and they work)' type of resistance. So, a picture of the actual current costs of bad antibody validation in terms of wasted funds (and research years, peoples careers' etc. etc. as documented in the various validation commentaries in the literature) would be helpful to stimulate interest." Participants noted: "Would need to make the case to funders on the amount of wasted funds in order to get their buy-in (more accurate costing needed)."

Technical challenges with specific tool types were identified: "Much of this could depend on the nature of the target. If there are no consistently reliable cell models that would make studying the protein of interest difficult. KO models could be lethal depending on the target and KD may not show a big difference if endogenous levels of a given protein are very high." This suggests that schemes need flexibility to accommodate target-specific constraints.

Regarding animal-derived antibody replacement, one participant cautioned: "If by 'animal derived' antibodies, you mean moving from polyclonals and hybridoma-derived toward recombinant, I would generally agree. Although I think that there will always be a role for polyclonals in early research and for certain applications. It is hard to make the case that research cannot be conducted until you are assured of a rigorously validated, recombinant monoclonal."

One opportunity identified was collaborative funding: "This may be a challenge in the orphan disease arena, where funding is limited. However, it can offer an opportunity for funders to collaborate (for example, as had occurred with the amyotrophic lateral sclerosis community in the past) in supporting antibody production and validation."

**R13: Funders should signal validation importance in guidance**

**(Effectiveness: 7.0, Feasibility: 7.0, accepted in Round 1)**

**The Recommendation**

Funders should share information within the applicant guidance that antibody performance is an important limitation in many experimental methods. Applicants should be encouraged to address this in methodological or reproducibility sections, as this will be part of how these sections are evaluated by reviewers and panels.

**Panel Assessment**

This recommendation achieved consensus as both effective (median 7.0) and feasible (median 7.0) without disagreement. The panel agreed that guidance-based signaling could influence practices and could be implemented by 2030, though with more reservation about effectiveness compared to stronger interventions.

**Supporting Rationale**

Panelists viewed this as a "low lift" intervention that could "push the needle and raise awareness" without creating hard requirements. As one participant noted: "If I understand this correctly, it would involve simply stating in the instructions for applications that antibody validation strategies and data should be included to support data interpretation and reproducibility. Yes, this is an easy and useful strategy." Another characterized it as "a nudge, and thinly veiled threat. But certainly a wake-up for people who STILL don't understand the issues."

However, several participants expressed skepticism about effectiveness without enforcement: "Doubtful that this would be strong enough to drive best practice" and "'Signalling Value Without Mandating' seems not much different than the current situation."

**Implementation Considerations**

The primary implementation challenge is that effectiveness depends entirely on how seriously reviewers treat the guidance: "Effectiveness depends on how strongly funders insist on the inclusion of this information and Panel training in the assessment of the information." One participant noted that reviewers "themselves probably require educating."

Panelists raised concerns that without consequences, this could be ineffective: "If it's not a mandate then incentive for applicants to do this is not there and therefore will not be effective" and "I'm not sure that sharing information alone is that effective without some consequences/reward for those authors that do not or do take heed of this."

The risk of generic compliance was identified: "Similar to R10, I think this will just end up in a copy/paste situation where applicants are paying lip service to the principle without affecting behavior." One participant noted: "This seems possible, but perhaps not strong enough to drive real change in researchers who have a range of other priorities. It might get overlooked."

The implementation question is whether funders can ensure that validation considerations genuinely factor into peer review scoring rather than becoming ignored boilerplate in applications.

**R14: Funders should encourage validation data deposition**

**(Effectiveness: 7.0, Feasibility: 7.0, accepted in Round 1)**

**The Recommendation**

Funders should encourage grantees to deposit antibody validation data in open-access repositories, ideally linked to RRIDs or registry entries. This expectation could be embedded in end-of-grant reporting or incentivised through visibility or reproducibility initiatives.

**Panel Assessment**

This recommendation achieved consensus as both effective (median 7.0) and feasible (median 7.0) without disagreement. The panel agreed that encouraging data deposition could improve practices and could be implemented by 2030.

**Supporting Rationale**

Panelists recognized the value of making validation data publicly accessible, though the recommendation uses "encourage" rather than "require," reflecting a softer approach. One participant suggested removing the word "ideally" from linking to RRIDs: "I think you can leave the 'ideally' out of the question. Unequivocal identification using for example RRID is easy and a simple step to getting antibodies validated."

**Implementation Considerations**

The primary challenges identified concerned infrastructure, quality control, and enforcement. Panelists noted the need for accessible repositories: "We will need a place to share this data that is relatively easy to use and well known." The quality of deposited data was a significant concern: "The issue here is the quality of the 'antibody validation data'" and "I have a similar concern about who will be checking the validation data to ensure robustness. I worry it will be a massive, meaningless data drop just to fulfill [sic] the requirement. It should be a reviewed/curated process but who would do this work?"

Enforcement at the end-of-grant stage was identified as particularly challenging: "It can be a challenge to robustly enforce end of grant reporting. Follow up requires staff resource which may not be available. Charitable funders in particular tend to focus resource on new studies (and funding commitments) out of necessity, in order to generate funds from the public." One participant was blunt: "The most you could do would be to 'encourage'. I don't think it's feasible to mandate or track compliance."

A coordination opportunity was identified: "Feasibility comes down to whether there is also education/training for authors about where and how to deposit the data and also the willingness of funders to monitor and enforce deposit. Note it would be easier for publishers to enforce at publication if there is an expectation that funders will also be monitoring and enforcing this at an earlier stage in the research lifecycle." This suggests that funder encouragement works best when aligned with journal requirements at the publication stage.

One participant noted a practical constraint: "Highly feasible, but only if authors already have the validation data," highlighting that encouragement to deposit presupposes that validation has been conducted.

**R15: Funders should endorse reporting standards**

**(Effectiveness: 7.0, Feasibility: 7.0, accepted in Round 1)**

**The Recommendation**

Funders should formally endorse community-developed reporting standards that promote antibody transparency and validation (e.g., IWGAV, MDAR). Such endorsements can help reinforce aligned expectations across funders and journals.

**Panel Assessment**

This recommendation achieved consensus as both effective (median 7.0) and feasible (median 7.0) without disagreement. The panel agreed that formal endorsement of existing standards could support better practices and could be implemented by 2030.

**Supporting Rationale**

Panelists viewed formal endorsement as a relatively low-effort intervention with potential for impact. One participant was optimistic: "I am marking this as a 9 on effectiveness as I feel that this would provide the type of incentive needed to drive a change in research practice/reporting. I expect 'endorsement' to be relatively feasible by funders. Endorsement could be in the form of a request to applicants to demonstrate past adherence to community-developed reporting standards as a potential pre-requisite for consideration."

The value of alignment between funders and publishers was emphasized: "If funders and publishers collaborate on the standards and funders then mandate those standards, publishers will generally be much more willing to ensure there is compliance. Endorsement alone will help but funders also need to require that standards are met to be really effective."

**Implementation Considerations**

The primary implementation challenge is that endorsement alone may be insufficient to change behavior. Several participants were skeptical: "Feasible and a low lift. But will most likely be ignored" and "This seems possible, but perhaps not strong enough to drive real change in researchers who have a range of other priorities. It might get overlooked."

Consistency across funders was identified as important: "Needs to be consistent across funders wrt which reporting standards are endorsed, unless all are held in equally in high regard." This suggests value in coordination between funders to endorse the same standards rather than fragmenting the landscape with different funder-specific requirements.

Institutional barriers were noted: "It is difficult to get funders to do things like this. We have tried, but they are frequently locked into various difficult to change rules." This points to potential bureaucratic or policy constraints that may slow adoption even of straightforward endorsements.

The key implementation question is whether endorsement can be meaningfully linked to grant evaluation (as suggested by one participant's proposal to make past adherence a "pre-requisite for consideration") or whether endorsements will remain aspirational statements with limited practical impact.

**1.3 Institutional and Educational Recommendations**

**R19: Training in antibody validation**

**(Effectiveness: 8.0, Feasibility: 7.0, accepted in Round 1)**

**The Recommendation**

Training in antibody validation should be offered in relevant bioscience courses and programmes. This could include: Undergraduate education (e.g., in immunology, neuroscience, or molecular biology programmes), Postgraduate or doctoral training (e.g., through doctoral training centres), Postdoctoral or continuing professional development. Training should be field-appropriate, not universally required across all domains.

**Panel Assessment**

This recommendation achieved consensus as both effective (median 8.0) and feasible (median 7.0) without disagreement. The panel agreed that educational interventions would meaningfully improve antibody practices and could realistically be implemented by 2030.

**Supporting Rationale**

Panelists strongly endorsed education as a foundational intervention: "I think education and training will be key to ultimately improving antibody validation - so that it just becomes the norm. Requirements and enforcement by funders or publishers will only be effective if researchers understand why it is so important and are trained appropriately (it's just too late by the time a study gets to publication)." The value of early-career training was emphasized: "It is so important that the young bioscientists see what a chaos bad antibodies cause, and much easier to teach them than the oldies. Start at the roots."

One participant argued for disciplinary recognition: "Antibody validation should be recognized as a research discipline and incorporated into cell biology curricula."

**Implementation Considerations**

The primary implementation challenges concerned reach, standardization, and resources. Multiple participants questioned: "Difficult to know how to influence this across a broad range of institutions?" and noted that "it would be difficult to get all institutions on board. Different institutions, countries, will have different priorities and policies."

The level at which training should be provided generated discussion: "Not necessary for undergraduates but should be part of PhD programs. Practical training would be challenging and may have cost barriers but a lecture on best practices for using antibodies and other reagents should be feasible." This points to potential trade-offs between breadth of coverage and depth of training.

A critical implementation gap was identified: "Need to train the senior staff as well!" This recognizes that "in many cases the emphasis on validation and proper controls comes down to lab culture. If a student is trained to validate antibodies but the PI/senior lab members do not agree this is a priority, it will not be done." Training students without addressing PI-level culture may be insufficient.

Resource questions were raised repeatedly: "How will training be funded? Who will deliver?" and "Better training is required, the questions that need to be addressed are: who funds this, who is responsible for the training, who sets the standards, etc." One participant acknowledged uncertainty about implementation mechanisms: "I have said that this should be very feasible in principle to implement but bear in mind that I'm not really sure how easily such training can be introduced systematically into institutions that do not have this and/or are unaware of the importance."

**R20: Institutional research integrity frameworks**

**(Effectiveness: 7.0, Feasibility: 7.0, accepted in Round 1)**

**The Recommendation**

Universities and research institutions should incorporate antibody validation expectations into research integrity and ethics review frameworks. This may include: Asking researchers to consider antibody validation as part of research ethics approval processes, Providing guidance or templates for documenting validation plans.

**Panel Assessment**

This recommendation achieved consensus as both effective (median 7.0) and feasible (median 7.0) without disagreement. The panel agreed that integrating antibody validation into existing institutional frameworks could improve practices and could be implemented by 2030.

**Supporting Rationale**

Panelists recognized the value of framing antibody validation as an integrity issue: "Incorporating training into existing ethics and integrity frameworks is a hugely important signal to researchers that this is not just good research practice but there is a moral imperative to ensure it is done correctly because of the immense damage done without appropriate validation. Such frameworks could include the consequences (economic, health, equity etc). Such training will hopefully help make researchers think twice before skipping a step or help incentivise researchers to do the extra steps required for validation."

The recommendation could work "as long as the guidance is clear enough, and ideally uniform across institutions."

**Implementation Considerations**

The most significant implementation challenge identified was whether antibody validation appropriately fits within ethics frameworks. One participant was skeptical: "I don't really see this as an ethical issue. Lumping it into ethics with big issues like image manipulation, data fabrication, animal use, etc doesn't seem appropriate to me, personally. Also, if a lab publishes without antibody validation data, could ethics complaints be lodged against them?"

Coverage limitations were noted: "A lot of research does not go through ethics committees," suggesting that ethics-based frameworks may not reach all relevant research.

Leadership awareness was identified as a barrier: "While such strategies should absolutely be in place today, the main barrier to doing so is ignorance of the issues by the leadership at all levels of academia." This points to the need for senior-level education before institutional frameworks can be effectively implemented.

Prioritization challenges were raised: "There is probably a long list of things that should be done, and getting something to the top of the list to implement is challenging. Demonstration of what is lost to future scientists by omitting this data would provide value." This suggests that making the case for antibody validation as a priority requires evidence of impact.

**R21: Local champions and expertise networks**

**(Effectiveness: 7.0, Feasibility: 7.0, accepted in Round 1)**

**The Recommendation**

Universities, research institutions and learned societies should recognise and support local champions or experts in antibody validation. These individuals may serve as: Points of contact for advice on antibody selection, validation design, and data interpretation; Trainers or facilitators in reproducibility workshops or research methods courses; Advocates for good practice within departments or core facilities; Contributors to institutional or cross-institutional reproducibility networks. Support could include protected time, acknowledgement in performance reviews, or funding for training and outreach.

**Panel Assessment**

This recommendation achieved consensus as both effective (median 7.0) and feasible (median 7.0) without disagreement. The panel agreed that supporting local expertise could improve practices and could be implemented by 2030.

**Supporting Rationale**

Panelists appreciated the flexibility of this approach: "I like this idea. It has the flexibility/opportunity to educate multiple levels of the system (students, postdocs, PIs) and provide a direct connection to someone for further discussion, insight, and support." One participant noted it "has the opportunity to educate multiple levels" and could be integrated into existing structures by "being rolled into outreach training."

The recommendation was recognized as foundational: "All of the questions in this section are not going to make antibody research more reproducible per se but will lay the foundation for increased awareness and adoption of best practice for researchers and disciplinary communities. Without laying such a groundwork of training and advocacy (as this specific question promotes), antibody validation will never become the norm."

**Implementation Considerations**

The primary implementation challenges concerned sustainability and institutional buy-in. One participant was pessimistic: "While I believe this is a much needed item, I don't believe this issue has much likelihood of rising to a sufficient level of awareness that it will be supported by an institute at this time." Experience with similar initiatives suggested challenges: "We tried this at our university, maybe it was not the right time, but we could not get a reproducibiliTea [sic] group to launch properly."

Staffing and sustainability questions were raised: "High turnover of research staff - how to maintain long term champions?" and "Would these roles be a full-time job or would this be someone doing it in addition to their research? It would require a lot of time but if it is a full time job who would fund it?" The tension between volunteer-based and funded positions remains unresolved.

Engagement was identified as uncertain: "Of course, it won't work if the users don't use the facility..." This points to the challenge that creating expertise resources does not guarantee uptake without accompanying cultural change or requirements that drive researchers to seek advice.

**1.4 Cross-Stakeholder and Manufacturer Recommendations**

**R22: Shared roadmap for stakeholder coordination**

**(Effectiveness: 7.0, Feasibility: 7.0, accepted in Round 1)**

**The Recommendation**

Stakeholders should work together to develop a shared roadmap for improving antibody validation practices by 2030. This would outline: Stakeholder-specific actions and responsibilities, Target timelines and milestones, Mechanisms for accountability and feedback.

**Panel Assessment**

This recommendation achieved consensus as both effective (median 7.0) and feasible (median 7.0) without disagreement. The panel agreed that cross-stakeholder coordination could improve practices and could be implemented by 2030, though with recognition of significant coordination challenges.

**Supporting Rationale**

Panelists endorsed the principle of coordinated action: "Organizing meetings of stakeholders at funding agencies (e.g., NIH), national/society meetings, and others should be encouraged and enacted asap" and "Engages various groups for informed recommendations across sectors." The need for coordination was recognized as addressing fragmentation: "This effort, broadly, is too fragmented. Alignment feels difficult to achieve across the various initiatives."

One participant noted the sustained nature of the problem: "This is a conversation that's been going on for about 30 years. I published a widely cited (my most cited) commentary in 2015 making the case for recombinant antibodies with published sequences. Since then there has been some improvement in antibody providers striving to sell recombinant antibodies, rather than polyclonals, but progress has been so so so slow. Everyone points fingers at everyone else to indicate where the responsibility should lie."

**Implementation Considerations**

The primary implementation challenges concerned consensus-building, leadership, and sustainability. Historical attempts at coordination have struggled: "There have been at least 2 initiatives to implement a framework for antibody validation across stakeholder groups that have started but were not able to develop a final product. The reasons for failing were different (one was funding related and the other was inability to reach a consensus) but they are illustrative of how difficult a problem this is to tackle."

Consensus-building challenges were noted: "It may be difficult to reach consensus, as shown by prior work within the IWGAV group, which may result from different motivations and drivers for different individuals" and "I could see these types of cross-stakeholder efforts getting bogged down. Everybody thinks they know best." One participant questioned: "How is this different from the IWGAV?" suggesting potential overlap with existing coordination bodies.

Leadership and accountability gaps were identified: "Scored low on feasibility as there will be challenges achieving cross sector collaboration and leadership." The question of who would drive coordination remained open.

Resource requirements were emphasized: "Time and funding are always major factors" and "Funding is required to get the right people together to create a roadmap and continued funding is required to get it implemented. These things traditionally take years or decades to achieve across all the various stakeholders but you have to start somewhere and gather momentum."

Timeline concerns were raised: "Rating 7 only because 2030 is not so far away and roadmap development involving multiple stakeholders can be a lengthy process." This points to tension between the 2030 target date and the typical timescales for multi-stakeholder coordination.

**A7: Manufacturers should assign RRIDs at source**

**(Effectiveness: 8.0, Feasibility: 7.0, introduced and accepted in Round 2)**

**The Recommendation**

Antibody manufacturers should be encouraged to assign RRIDs to their products at source.

**Panel Assessment**

This recommendation achieved consensus as both effective (median 8.0) and feasible (median 7.0) without disagreement. The panel agreed that upstream RRID assignment would meaningfully improve antibody identification and could realistically be implemented by 2030.

**Supporting Rationale**

Panelists responded enthusiastically to this recommendation, with one noting: "Really, an easy one" and another stating "this would be fantastic." The simplicity and directness of the intervention contributed to strong support.

One participant suggested that existing practices may already be sufficient: "Listing the antibody's full name, including clone and possibly Lot# along with the manufacturer's name is more than sufficient," though this comment may reflect uncertainty about the added value of RRIDs beyond traditional identifiers.

**Implementation Considerations**

The primary implementation challenge identified was manufacturer willingness to participate. One participant noted: "There are some suppliers that will ignore this, in the same way they have ignored the calls to improve validation." This points to variation in manufacturer responsiveness to community standards.

The mechanism of influence was questioned: "This should be feasible - but it requires manufacturers to be willing and I don't know whether manufacturers take heed of funders (in the same way that some funders have mandates around open access and metadata, which are then ignored by some publishers)." This suggests that "encouragement" alone may be insufficient, and that consideration should be given to what incentives or pressures would motivate manufacturer participation.

The recommendation specifies "encouragement" rather than mandates, leaving open the question of who would encourage manufacturers and through what mechanisms. Potential avenues could include funder requirements (researchers must use antibodies with RRIDs), journal policies (manuscripts must report RRIDs), or market-based incentives (researchers preferentially purchase products with RRIDs). The connection to R1 (authors include RRIDs) and R2 (complete metadata reporting) is apparent—if researchers are required to report RRIDs, demand for manufacturer-assigned RRIDs would increase.
