## Supplementary material for "A multi-stakeholder Delphi consensus study proposes actionable solutions to address antibody validation failures": S2 Text

**[DRAFT FOR CONSULTATION]**

**Recommendations for Publishers and Journals**

*Improving Antibody Validation in Biomedical Research*

Based on findings from an MRC-funded Delphi consensus study

### Executive Summary

A significant proportion of antibodies used in published biomedical research are not fit for the specific purpose for which they are used. This leads to unreliable findings, economic waste, and the unnecessary use of patient and animal samples. Technical solutions and data-sharing initiatives exist, but coordinated stakeholder action is needed to embed better practices across the research ecosystem.

Through an NC3Rs-convened stakeholder meeting and an MRC-funded Delphi study, a panel of 32 international experts identified interventions to improve antibody validation that are both effective and feasible for implementation by 2030. Publishers and journals operate at the point where research enters the public record, making them a natural place to establish expectations for antibody identification and validation.

This document asks publishers to act on **two priority recommendations ready for implementation now**: establishing a complete antibody reporting package and adopting clear antibody validation standards. Four further actions were rated as effective but face implementation barriers — this document outlines how publishers can begin to address those.

This is part of a coordinated strategy with parallel consultation documents for research funders, institutions and educational bodies, and antibody manufacturers. We welcome your feedback and invite participation in a proposed working group to develop practical implementation guidance.

### About This Document

#### Background

This document is one of four stakeholder-specific consultation documents developed from an MRC-funded Delphi consensus study on antibody validation in biomedical research. A panel of 32 experts participated in two rounds of structured assessment, rating proposed interventions on effectiveness (ability to improve antibody validation in published research, scale 1–9) and feasibility (realistic implementation across the field by 2030, scale 1–9). Items achieving a median of ≥7 on both dimensions without panel disagreement are classified as consensus recommendations. Where multiple Delphi items addressed related issues with similar ratings, they have been merged for clarity; original item codes (e.g., R1, R2) are provided for cross-referencing.

Full study methods and results are published in the accompanying manuscript, with complete qualitative commentary from panellists available in S7 and S8 Texts. Parallel consultation documents have been prepared for research funders, institutions and educational bodies, and antibody manufacturers.

#### Key Terminology

**Validation** refers to experimental evidence that an antibody is performing as claimed in a specific experiment — that it is interacting selectively with its intended target in the specific assay, tissue, or sample type used. Validation is context-specific: an antibody validated for Western blot in one cell type is not validated for immunofluorescence in another.

**Characterisation** refers to systematic experiments showcasing the performance qualities (or limitations) of an antibody across standardised conditions. Characterisation data (such as that generated by YCharOS or displayed on the OGA Antibody Database) can help researchers assess whether an antibody is likely to perform well, but does not replace the need for context-specific validation where results depend on antibody specificity.

### Priority Actions for Publishers

The following recommendations achieved consensus as both effective and feasible, representing actions suitable for immediate implementation.

#### Complete Antibody Reporting Package

**Recommendation:** Authors should provide complete antibody information for each antibody used, including: Research Resource Identifier (RRID) where available, clone ID (for monoclonal antibodies) or lot number (for polyclonal antibodies), catalogue number, vendor/source, dilution ratio used, and protein concentration where possible.

This merges three related consensus items:

| **Item** | **Description** | **Effectiveness** | **Feasibility** |
| --- | --- | --- | --- |
| **R1** | Authors include RRIDs for each antibody used | 7.0 | 8.0 |
| **R2** | Authors provide complete antibody metadata (clone ID, catalogue number, lot number, vendor/source) | 8.0 | 8.0 |
| **R3** | Authors report dilution ratio and protein concentration where possible | 8.0 | 8.0 |

These requirements work synergistically: RRIDs enable unique identification and tracking across the literature; metadata ensures the specific product can be identified even without an RRID; and dilution/concentration data supports experimental replication. Together, they create the foundation on which other recommendations in this document build — enabling antibodies to be checked against characterisation databases, linked to validation data, and screened for known problems.

##### Implementation Options

1. **Adopt the MDAR (Materials Design Analysis Reporting) framework.** MDAR already covers RRID, vendor, catalogue number, clone/lot number, and dilution. It would require only the addition of protein concentration as an optional field.
2. **Implement automated compliance checking.** Tools such as SciScore can flag missing information at submission. Cost-sharing models across publishers could make this accessible to smaller journals.
3. **Create structured submission templates.** Required fields in manuscript submission systems capture information at the point of entry, avoiding post-hoc checking.

#### Clear Antibody Validation Standards

**Recommendation:** Journals should establish clear standards for antibody validation reporting that authors must meet.

| **Item** | **Description** | **Effectiveness** | **Feasibility** |
| --- | --- | --- | --- |
| **A1** | Journals establish clear antibody validation and reporting standards | 7.5 | 7.0 |

Clear standards give practical meaning to the reporting requirements above. Without them, authors may report antibody identifiers while lacking evidence that the antibody performed as claimed in their specific experiment. The MDAR framework already includes validation reporting requirements, and the IWGAV (International Working Group for Antibody Validation) five-pillar framework provides specific validation approaches that could strengthen what “validation” means in practice. Because validation is context-specific (see Key Terminology), standards should reflect this rather than implying a binary validated/not-validated status.

##### Implementation Options

1. **Adopt existing MDAR validation language** as a baseline requirement.
2. **Strengthen MDAR by explicitly referencing the IWGAV framework.** Ask authors to indicate which IWGAV-aligned evidence supports their specific antibody-dependent data. The panel separately assessed IWGAV referencing (R8: Effectiveness 7.0, Feasibility 6.0), suggesting phased introduction may be needed.
3. **Require transparency about validation gaps.** Where context-specific validation is lacking, authors should acknowledge this limitation.
4. **Allow reference to published validation data** where experimental conditions closely match (same application, similar sample type).

### Longer-Term Actions: Building Toward Greater Impact

The following four recommendations were all rated as effective (median ≥7) but face implementation barriers around cost, infrastructure, editorial resources, or technical constraints. For each, we suggest practical ways publishers can begin working toward implementation. Connections to parallel stakeholder actions (funder, institutional, manufacturer) are noted in the Support section.

#### Making Validation Data Accessible

**Recommendation:** Authors should make antibody validation data accessible through deposition in open-access repositories linked to persistent identifiers (e.g., RRID, DOI) or through manuscript supplementary materials.

This merges two related items:

| **Item** | **Description** | **Effectiveness** | **Feasibility** |
| --- | --- | --- | --- |
| **A2** | Include validation protocols/data in manuscripts or supplements | 8.0 | 6.0 |
| **R4** | Require validation data deposition in open-access repositories | 8.0 | 5.0 |

Once deposited and linked to RRIDs, validation data becomes a shared resource — available to future researchers and reviewers, reducing duplication.

##### How Publishers Can Begin

1. **Apply requirements to critical antibodies only.** Focus on antibodies whose specificity is necessary to interpret the main result, not all antibodies in a paper.
2. **Provide clear author guidance on repository options.** Several repositories exist (Biomed Resource Watch, Zenodo, F1000 data gateways) but are not well known. Clear instructions with examples would reduce the barrier.
3. **Allow citation of existing validation data** where experimental conditions closely match, as an alternative to generating new data.

#### Automated Screening Tools for Antibody Information

**Recommendation:** Journals should adopt automated tools to screen manuscripts for incomplete antibody metadata or flag antibodies withdrawn from the market due to performance issues.

| **Item** | **Description** | **Effectiveness** | **Feasibility** |
| --- | --- | --- | --- |
| **R7** | Automated tools to screen manuscripts and flag antibody issues | 8.0 | 5.0 |

The panel agreed this would be highly effective but expressed uncertainty about implementation by 2030, primarily due to cost and technical integration challenges that individual publishers — particularly smaller ones — may not be able to sustain alone.

##### How Publishers Can Begin

1. **Pilot with existing tools.** Test available tools (e.g., SciScore for RRID checking) to establish evidence of effectiveness before broader investment.
2. **Develop a consortium model for cost-sharing.** Multiple publishers pooling resources to develop or license shared screening infrastructure reduces per-publisher costs significantly.
3. **Adopt a tiered rollout.** Larger publishers adopt first, demonstrating value and developing shared tools that smaller publishers can later access.
4. **Support community-developed open-source solutions.** Freely available screening scripts that integrate across editorial platforms could make automated checking accessible to all publishers.

#### Genetic Validation for Critical Antibodies

**Recommendation:** Where antibodies are critical to key experimental conclusions and technically feasible, authors should either present validation data based on a genetic strategy (e.g., knockout or knockdown) or cite previously published genetic validation data.

| **Item** | **Description** | **Effectiveness** | **Feasibility** |
| --- | --- | --- | --- |
| **R9** | Genetic validation for antibodies critical to key conclusions | 8.0 | 5.0 |

Genetic validation represents the strongest form of antibody specificity evidence, but it is not universally applicable. Essential genes cannot be knocked out, some targets are technically intractable, and the cost can be substantial. The IWGAV framework provides alternative validation approaches for such targets.

##### How Publishers Can Begin

1. **Encourage rather than mandate.** Frame as best practice, acknowledging technical constraints for certain targets.
2. **Accept alternative IWGAV validation methods** (independent antibodies, orthogonal methods) when genetic strategies are not feasible.
3. **Provide objective criteria for “critical.”** Develop guidance on which antibody uses qualify (e.g., novel target identification, therapeutic target validation, biomarker discovery).
4. **Allow citation of published genetic validation** where experimental conditions are comparable.

#### Building Editorial Capacity for Validation Assessment

**Recommendation:** Journals should build capacity to evaluate antibody validation through training reviewers and editors, or by appointing specialist editors focused on reproducibility.

This merges two related items:

| **Item** | **Description** | **Effectiveness** | **Feasibility** |
| --- | --- | --- | --- |
| **R6** | Train reviewers and editors on antibody validation assessment | 7.0 | 5.0 |
| **A3** | Appoint specialist reproducibility editors | 7.0 | 5.5 |

##### How Publishers Can Begin

1. **Provide lightweight reviewer checklists.** Integrate antibody validation prompts into existing review forms rather than requiring extensive separate training.
2. **Develop shared training resources across publishers.** Collaborate on reusable educational materials, drawing on existing resources such as OGA Academy modules.
3. **Appoint cross-journal reproducibility editors.** Specialist editors operating across a journal portfolio makes this feasible for larger publishers while building a model others could adopt.
4. **Offer expert review services for complex cases.** Provide access to antibody validation specialists when needed, as piloted by Lifecycle journal.

### Support, Resources, and Next Steps

#### Stakeholder Coordination

These recommendations are part of a coordinated strategy. The Delphi panel endorsed a shared roadmap for stakeholder coordination (R22: Effectiveness 7.0, Feasibility 7.0), recognising that sustained improvement requires aligned action. Each stakeholder acting independently strengthens the conditions for others: as publishers require reporting, researchers demand RRIDs from manufacturers; as funders signal the importance of validation, authors arrive at submission better prepared; as institutions train researchers, the reviewer pool with relevant expertise grows.

#### Shared Infrastructure

The panel assessed coordinated infrastructure for aggregating antibody validation data (R23: Effectiveness 8.0, Feasibility 6.0). While not achieving consensus on feasibility, this received a high effectiveness rating, reflecting the view that shared data infrastructure would be transformative. Several longer-term recommendations in this document (automated screening, accessible validation data) would benefit significantly from such infrastructure.

#### Resources Available

- **YCharOS** (<https://zenodo.org/communities/ycharos>): Open antibody characterisation data generated through independent benchmarking using knockout cell lines. The primary data source underlying many of the recommendations in this document.
- **OGA Antibody Database** (onlygoodantibodies.co.uk): Curated, searchable interface for antibody characterisation data across Western Blot, immunoprecipitation, ICC/IF, and flow cytometry — designed to reduce the work involved in making informed antibody decisions. Can support editorial assessment and reviewer guidance.
- **OGA Academy** (onlygoodantibodies.co.uk/academy): Four free eLearning modules covering antibody selection and validation.
- **MDAR framework**: Established reporting checklist already adopted by some journals.

#### Proposed Next Steps

We propose forming a working group to develop practical implementation guidance, including template language for author guidelines, reviewer checklists, and guidance on adopting MDAR with IWGAV-aligned validation standards. We will be working with NC3Rs and other partners to convene this.

We welcome your feedback on which implementation options are most feasible for your context, barriers we have not adequately addressed, interest in pilot implementations, and willingness to join the proposed working group.

#### Contact

Dr Harvinder Virk

University of Leicester
