## Supplementary material for "A multi-stakeholder Delphi consensus study proposes actionable solutions to address antibody validation failures": S3 Text

**[DRAFT FOR CONSULTATION]**

**Recommendations for Research Funders**

*Improving Antibody Validation in Biomedical Research*

Based on findings from an MRC-funded Delphi consensus study

### Executive Summary

A significant proportion of antibodies used in published biomedical research are not fit for the specific purpose for which they are used. This leads to unreliable findings, economic waste, and the unnecessary use of patient and animal samples. Technical solutions and data-sharing initiatives exist, but coordinated stakeholder action is needed to embed better practices across the research ecosystem.

Through an NC3Rs-convened stakeholder meeting and an MRC-funded Delphi study, a panel of 32 international experts identified interventions to improve antibody validation that are both effective and feasible for implementation by 2030. Funders shape researcher behaviour through application requirements, resource allocation, and signalling priorities. They operate at a critical point in the research lifecycle: before experiments are conducted, when validation planning and budgeting can be embedded from the outset.

This document asks funders to act on **three priority recommendations ready for implementation now**: providing financial support for antibody validation (dedicated budget lines and tool development schemes), addressing antibody validation in grant applications (through guidance and/or requirements), and supporting community standards and data sharing. Four further actions were rated as effective but face implementation barriers — this document outlines how funders can begin to address those.

### Priority Actions for Funders

The following recommendations achieved consensus as both effective and feasible, representing actions suitable for immediate implementation. Six individual Delphi items are organised into three thematic recommendations.

#### Funding Support for Validation

**Recommendation:** Funders should provide financial support for antibody validation through dedicated budget lines in grant applications and targeted tool development schemes.

- **Dedicated budget lines:** Applicants to biomedical funding schemes should include a specific line item for antibody validation resources (where relevant to their proposal).
- **Targeted tool development schemes:** Create or expand schemes to support development and validation of critical antibody-based research tools, including characterisation of under-studied antibodies, generation of knockout/knockdown cell lines, cross-platform benchmarking, and replacement of animal-derived antibodies.

| **Item** | **Description** | **Effectiveness** | **Feasibility** |
| --- | --- | --- | --- |
| **R11** | Budget line for antibody validation in grant applications | 8.0 | 7.0 |
| **R12** | Targeted tool development schemes for antibody validation | 8.0 | 7.0 |

Budget lines support validation within individual projects; tool development schemes create shared infrastructure benefiting the wider community.

##### Implementation Options

***For budget lines:***

1. **Tiered requirement based on antibody criticality.** Require validation budgets only when antibodies are central to main activities, not for every antibody in a proposal.
2. **Provide budget guidance.** Develop guidance on reasonable validation costs to help applicants budget appropriately.
3. **Phased implementation.** Pilot with specific funding schemes before broader rollout.

***For tool development schemes:***

1. **Strategic prioritisation.** Focus on high-impact targets (common use, poor current options, critical disease areas) rather than attempting comprehensive coverage.
2. **Collaborative funding models.** Partner across funders to share costs, particularly valuable in the orphan disease arena where individual funder budgets are limited.
3. **Build on existing initiatives.** Expand proven models (like the YCharOS consortium) rather than creating new infrastructure.

#### Grant Application Requirements

**Recommendation:** Funders should address antibody validation in grant applications through escalating approaches:

- **Signal importance in guidance (lower threshold):** Include information in applicant guidance that antibody performance is an important limitation in many experimental methods. Encourage applicants to address this in methodological sections, noting it will be part of evaluation.
- **Require validation plans (stronger requirement):** Include a section in application forms where applicants must detail the steps they will take to validate the antibodies they will use.

| **Item** | **Description** | **Effectiveness** | **Feasibility** |
| --- | --- | --- | --- |
| **R13** | Signal importance of antibody validation in applicant guidance | 7.0 | 7.0 |
| **R10** | Require antibody validation plans in grant applications | 7.5 | 8.0 |

R10 had higher effectiveness, reflecting the view that explicit requirements drive more change than soft encouragement. Funders can start with R13 to raise awareness, then move to R10 once the groundwork is established.

##### Implementation Options

***For signalling importance (R13):***

1. **Add to existing guidance sections.** Incorporate antibody validation into established reproducibility or methods guidance.
2. **Link to training resources.** Point applicants to resources on validation best practices (IWGAV framework, OGA Academy).
3. **Combine with reviewer guidance.** Ensure review criteria explicitly mention antibody validation.

***For requiring validation plans (R10):***

1. **Provide structured templates based on existing models.** Adapt proven formats (e.g., NIH “key biological resources” file) with specific prompts to prevent generic responses.
2. **Focus review on validation of preliminary data.** Reviewers assess validation evidence supporting antibody-dependent data already presented in applications. This addresses the timing challenge — rather than asking applicants to predict future validation, it evaluates the reliability of evidence already generated.
3. **Integrate into milestone reporting.** Link validation expectations to progress reporting to ensure follow-through.

#### Community Standards and Data Sharing

**Recommendation:** Funders should support community-wide antibody validation practices through endorsing reporting standards and encouraging validation data deposition.

- **Endorse reporting standards:** Formally endorse community-developed standards that promote antibody transparency and validation (e.g., IWGAV, MDAR).
- **Encourage validation data deposition:** Encourage grantees to deposit antibody validation data in open-access repositories, ideally linked to RRIDs or registry entries.

| **Item** | **Description** | **Effectiveness** | **Feasibility** |
| --- | --- | --- | --- |
| **R15** | Endorse community-developed antibody reporting standards | 7.0 | 7.0 |
| **R14** | Encourage grantees to deposit antibody validation data | 7.0 | 7.0 |

##### Implementation Options

***For endorsing standards (R15):***

1. **Align with publisher policies.** Coordinate endorsement with major journals so applicants see consistent expectations across the research lifecycle.
2. **Incorporate into application requirements.** Move beyond endorsement to requiring demonstration of adherence in applications.

***For encouraging data deposition (R14):***

1. **Provide infrastructure guidance.** Clear information on available repositories with examples of good practice.
2. **Align with publication stage.** Coordinate with journal requirements for data deposition at publication, which is easier to enforce than post-grant follow-up.
3. **Incentivise through visibility.** Feature grantees who share validation data in communications or reproducibility showcases.

### Longer-Term Actions: Building Toward Greater Impact

The following four recommendations were all rated as effective (median ≥7) but face implementation barriers around resource alignment, funder mandates, or cross-sector coordination. For each, we suggest practical ways funders can begin working toward implementation. Connections to parallel stakeholder actions are noted in the Support section.

#### Co-Fund Benchmarking Initiatives

**Recommendation:** Funders should collaborate with independent benchmarking initiatives such as YCharOS or equivalent open-science consortia, directly co-funding the validation of antibodies against key targets to create suites of validated tools.

| **Item** | **Description** | **Effectiveness** | **Feasibility** |
| --- | --- | --- | --- |
| **R17** | Co-fund independent antibody benchmarking initiatives | 8.0 | 6.0 |

The panel rated this as the highest-effectiveness funder action. Concerns centred on whether this type of infrastructure funding falls within the scope of disease-specific funders and whether existing funding mechanisms can accommodate it.

##### How Funders Can Begin

1. **Multi-funder consortium model.** Pool resources across funders. Disease-specific funders partner with infrastructure-focused funders to co-fund target-specific characterisation.
2. **Align target selection with funder priorities.** Focus benchmarking on antibodies critical to specific disease areas, enabling disease-specific funders to justify investment as supporting their mission.
3. **Phased funding commitment.** Start with pilot funding to demonstrate impact on research quality before seeking long-term commitment.
4. **Integrate with tool development schemes.** Benchmarking initiatives could be one type of project supported under the consensus recommendation for tool development funding.

#### Promote Researcher Participation in Benchmarking

**Recommendation:** Funders should promote researcher participation in independent benchmarking initiatives by asking applicants to consider working with initiatives such as YCharOS or equivalent open-science consortia. This could be included in guidance notes or flagged at the panel review stage.

| **Item** | **Description** | **Effectiveness** | **Feasibility** |
| --- | --- | --- | --- |
| **R18** | Promote researcher participation in benchmarking initiatives | 7.0 | 6.0 |

##### How Funders Can Begin

1. **Include in guidance as suggestion.** Low-lift awareness-raising that helps normalise engagement with benchmarking initiatives.
2. **Connect to budget line recommendation.** Allow applicants to budget for benchmarking participation under the consensus recommendation for validation budget lines.
3. **Target antibody development grants specifically.** Focus on applicants developing new antibodies or tools, where benchmarking is a natural part of the validation process.

#### Funded Antibodies Must Be Recombinant and Publicly Available

**Recommendation:** Funders should mandate that if funds are used to develop an antibody as part of a research project, the reagent must be a recombinant antibody and must be made available to the wider community.

| **Item** | **Description** | **Effectiveness** | **Feasibility** |
| --- | --- | --- | --- |
| **A6** | Funded antibodies must be recombinant and publicly available | 7.0 | 6.0 |

The panel endorsed the principle but identified practical barriers around grant timelines (recombinant development takes 12+ months), enforcement challenges with institutional technology transfer offices, and the need for priority use periods.

##### How Funders Can Begin

1. **Prospective requirement for new grants.** Setting expectations before work begins is more feasible than retrospective enforcement.
2. **Allow a priority use period.** Permit the generating lab a defined period of exclusive use before requiring deposit or distribution.
3. **Require reporting on availability.** Require grantees to report how funded antibodies will be made available, even if enforcement of deposit remains challenging.

#### Engage Antibody Manufacturers

**Recommendation:** Funders should directly engage with manufacturers to encourage them to publish antibody validation datasets and adopt consistent metadata standards (e.g., RRID, clone ID, lot number).

| **Item** | **Description** | **Effectiveness** | **Feasibility** |
| --- | --- | --- | --- |
| **R16** | Funders engage manufacturers on validation data and metadata standards | 7.0 | 5.0 |

The panel agreed this could improve quality but questioned whether funders have appropriate leverage over commercial entities. The feasibility score reflects this uncertainty about the funder–manufacturer relationship.

##### How Funders Can Begin

1. **Market-based incentive approach.** Maintain and share lists of manufacturers meeting validation and metadata standards, creating competitive pressure.
2. **Coordinate with journal requirements.** Leverage publisher policies that require metadata and RRIDs to create market pressure on manufacturers.
3. **Support third-party curation.** Instead of engaging each manufacturer individually, support infrastructure that aggregates and makes visible which manufacturers meet standards. The parallel manufacturer consultation document addresses these recommendations from the manufacturer perspective.

### Support, Resources, and Next Steps

#### Stakeholder Coordination

These recommendations are part of a coordinated strategy. The Delphi panel endorsed a shared roadmap for stakeholder coordination (R22: Effectiveness 7.0, Feasibility 7.0), recognising that sustained improvement requires aligned action. Each stakeholder acting independently strengthens the conditions for others: as funders require validation plans, researchers seek training from institutions; as funders endorse reporting standards, publishers can align their requirements; as both create expectations, manufacturers face market pressure to provide better-identified, better-characterised products.

#### Shared Infrastructure

The panel assessed coordinated infrastructure for aggregating antibody validation data (R23: Effectiveness 8.0, Feasibility 6.0). While not achieving consensus on feasibility, this received a high effectiveness rating, reflecting the view that shared data infrastructure would be transformative. Several recommendations in this document (tool development schemes, benchmarking co-funding, data deposition) would benefit from such infrastructure.

#### Resources Available

- **YCharOS** (<https://zenodo.org/communities/ycharos>): Open antibody characterisation data generated through independent benchmarking using knockout cell lines. Can inform grant review — reviewers can check whether antibodies proposed or used in preliminary data have independent characterisation support.
- **OGA Antibody Database** (onlygoodantibodies.co.uk): Curated, searchable interface for antibody characterisation data — designed to reduce the work involved in making informed antibody decisions.
- **OGA Academy** (onlygoodantibodies.co.uk/academy): Four free eLearning modules covering antibody selection and validation, appropriate for early career researchers and beyond.
- **Template language for grant application forms and reviewer guidance** (in development).

#### Proposed Next Steps

We propose forming a working group to develop practical implementation guidance for the consensus recommendations, including template language for application forms, reviewer guidance, and budget estimation tools. We will be working with NC3Rs and other partners to convene this.

We welcome your feedback on which implementation options are most feasible for your context, barriers we have not adequately addressed, interest in pilot implementations, and willingness to join the proposed working group.

#### Contact

Dr Harvinder Virk

University of Leicester
