## Supplementary material for "A multi-stakeholder Delphi consensus study proposes actionable solutions to address antibody validation failures": S4 Text

**[DRAFT FOR CONSULTATION]**

**Recommendations for Universities, Research Institutions, and Educational Bodies**

*Improving Antibody Validation in Biomedical Research*

Based on findings from an MRC-funded Delphi consensus study

### Executive Summary

A significant proportion of antibodies used in published biomedical research are not fit for the specific purpose for which they are used. This leads to unreliable findings, economic waste, and the unnecessary use of patient and animal samples. Technical solutions and data-sharing initiatives exist, but coordinated stakeholder action is needed to embed better practices across the research ecosystem.

Through an NC3Rs-convened stakeholder meeting and an MRC-funded Delphi study, a panel of 32 international experts identified interventions to improve antibody validation that are both effective and feasible for implementation by 2030. Universities and research institutions shape the culture, training, and frameworks within which antibody use occurs. The panel identified institutional actions as foundational: without training and local expertise, downstream interventions by publishers and funders are less likely to achieve lasting change.

This document asks institutions to act on **three recommendations, all of which achieved full consensus**: embedding antibody validation training in relevant bioscience courses, incorporating validation expectations into research integrity frameworks, and supporting local champions and expertise networks. Unlike several publisher and funder recommendations where feasibility barriers were identified, the panel expressed no reservations about the feasibility of institutional actions.

This is part of a coordinated strategy with parallel consultation documents for publishers and journals, research funders, and antibody manufacturers. We welcome your feedback and invite participation in a proposed working group to develop practical implementation guidance.

### About This Document

#### Background

This document is one of four stakeholder-specific consultation documents developed from an MRC-funded Delphi consensus study on antibody validation in biomedical research. A panel of 32 experts participated in two rounds of structured assessment, rating proposed interventions on effectiveness (ability to improve antibody validation in published research, scale 1–9) and feasibility (realistic implementation across the field by 2030, scale 1–9). Items achieving a median of ≥7 on both dimensions without panel disagreement are classified as consensus recommendations.

Full study methods and results are published in the accompanying manuscript, with complete qualitative commentary from panellists available in S7 and S8 Texts. Parallel consultation documents have been prepared for publishers and journals, research funders, and antibody manufacturers.

#### Key Terminology

**Validation** refers to experimental evidence that an antibody is performing as claimed in a specific experiment — that it is interacting selectively with its intended target in the specific assay, tissue, or sample type used. Validation is context-specific: an antibody validated for Western blot in one cell type is not necessarily validated for immunofluorescence in another.

**Characterisation** refers to systematic experiments showcasing the performance qualities (or limitations) of an antibody across standardised conditions. Characterisation data (such as that generated by YCharOS or displayed on the OGA Antibody Database) can help researchers assess whether an antibody is likely to perform well, but does not replace the need for context-specific validation where results depend on antibody specificity.

### Priority Actions for Institutions

All three institutional recommendations achieved full consensus as both effective and feasible, representing actions suitable for immediate implementation. The three recommendations are complementary: training builds knowledge, integrity frameworks embed expectations, and local champions provide ongoing support and culture change.

#### Training in Antibody Validation

**Recommendation:** Training in antibody validation should be offered in relevant bioscience courses and programmes, including undergraduate education (e.g., immunology, neuroscience, molecular biology), postgraduate and doctoral training (e.g., through doctoral training centres), and continuing professional development. Training should be tailored to the research field and experience level of participants.

| **Item** | **Description** | **Effectiveness** | **Feasibility** |
| --- | --- | --- | --- |
| **R19** | Training in antibody validation across relevant bioscience programmes | 8.0 | 7.0 |

This was the highest-rated institutional recommendation on effectiveness.

##### Implementation Options

1. **Embed in doctoral training programmes.** Integrate antibody validation into existing research methods modules. OGA already delivers masterclasses for doctoral training programmes that are ready for wider rollout, providing a tested model institutions can adopt.
2. **Use existing training resources.** The OGA Academy provides four free eLearning modules covering antibody selection and validation strategies. These can supplement institutional teaching without requiring content development from scratch.
3. **Include continuing professional development for established researchers.** Address the gap in senior staff awareness through seminar series, workshop programmes, or integration into institutional research integrity training. Framing validation as a research quality issue may increase engagement from established PIs.

#### Institutional Research Integrity Frameworks

**Recommendation:** Universities and research institutions should incorporate antibody validation expectations into research integrity and ethics review frameworks. This may include asking researchers to consider antibody validation as part of research ethics approval processes, and providing guidance or templates for documenting validation plans.

| **Item** | **Description** | **Effectiveness** | **Feasibility** |
| --- | --- | --- | --- |
| **R20** | Incorporate antibody validation into institutional research integrity frameworks | 7.0 | 7.0 |

##### Implementation Options

1. **Integrate into research integrity training rather than ethics review.** Many institutions already have mandatory research integrity training — adding an antibody validation component addresses coverage concerns while reaching a wider audience than ethics committees alone.
2. **Provide institutional validation plan templates.** Standard templates help researchers document their antibody validation plans, aligned with emerging funder expectations for validation information in grant applications.
3. **Frame around waste reduction and research quality.** To secure leadership buy-in, present antibody validation as a cost-saving and quality measure rather than a compliance burden. Evidence on wasted resources and animal/patient sample waste provides a compelling institutional case.
4. **Align with emerging external expectations.** Institutional frameworks that embed validation from the outset position researchers well as requirements across the sector evolve.

#### Local Champions and Expertise Networks

**Recommendation:** Universities, research institutions, and learned societies should recognise and support local champions or experts in antibody validation. These individuals may serve as points of contact for advice on antibody selection and validation design, trainers in reproducibility workshops, advocates for good practice within departments or core facilities, and contributors to institutional or cross-institutional networks. Support could include protected time, acknowledgement in performance reviews, or funding for training and outreach.

| **Item** | **Description** | **Effectiveness** | **Feasibility** |
| --- | --- | --- | --- |
| **R21** | Support local champions and expertise networks for antibody validation | 7.0 | 7.0 |

##### Implementation Options

1. **Build on the Only Good Antibodies–NC3Rs Antibody Champions scheme.** This 12-month programme recruits early career researchers to drive antibody validation and best practice within their institutions. Institutions can nominate and support participants, providing protected time and recognising the role in performance reviews.
2. **Integrate into core facility structures.** Embed antibody validation expertise within existing core facilities (imaging, flow cytometry, proteomics) where staff already advise on reagent use. This addresses sustainability by linking to funded positions.
3. **Create cross-institutional networks.** Where individual institutions cannot sustain dedicated champions, regional or disciplinary networks can share expertise. Learned societies could coordinate discipline-specific networks.
4. **Connect champions to training delivery.** Champions serve as local trainers, linking the expertise role with the training recommendation. The same individuals deliver training and provide ongoing consultation, making both more sustainable.

### Support, Resources, and Next Steps

#### Stakeholder Coordination

These recommendations are part of a coordinated strategy. The Delphi panel endorsed a shared roadmap for stakeholder coordination (R22: Effectiveness 7.0, Feasibility 7.0), recognising that sustained improvement requires aligned action. Institutional actions are foundational to the wider strategy: as institutions train researchers and embed validation expectations, those researchers are better placed to meet funder requirements for validation plans and publisher expectations for validation reporting, which in turn creates market pressure on manufacturers to provide better-identified, better-characterised products.

#### Shared Infrastructure

The panel assessed coordinated infrastructure for aggregating antibody validation data (R23: Effectiveness 8.0, Feasibility 6.0). While not achieving consensus on feasibility, this received a high effectiveness rating. Shared data infrastructure would support institutional training by providing accessible, curated examples of validation and characterisation data for teaching purposes.

#### Resources Available

- **OGA Academy** (onlygoodantibodies.co.uk/academy): Four free eLearning modules covering antibody selection and validation. Ready for integration into institutional teaching.
- **OGA Antibody Database** (onlygoodantibodies.co.uk): Curated, searchable interface for antibody characterisation data across Western Blot, immunoprecipitation, ICC/IF, and flow cytometry — designed to reduce the work involved in making informed antibody decisions.
- **YCharOS** (<https://zenodo.org/communities/ycharos>): Open antibody characterisation data generated through independent benchmarking — the primary data source curated in the OGA Database.
- **Only Good Antibodies–NC3Rs Antibody Champions scheme** (<https://nc3rs.org.uk/only-good-antibodies-nc3rs-antibody-champions-scheme>): A 12-month programme recruiting early career researchers to drive antibody validation and best practice within their institutions.

#### Proposed Next Steps

We propose forming a working group to develop practical implementation guidance, including model training curricula, institutional policy templates, and guidance on establishing or supporting local champions. We will be working with NC3Rs and other partners to convene this.

We welcome your feedback on which implementation options are most feasible for your context, barriers we have not adequately addressed, interest in pilot implementations, and willingness to join the proposed working group.

#### Contact

Dr Harvinder Virk

University of Leicester
