## Supplementary material for "A multi-stakeholder Delphi consensus study proposes actionable solutions to address antibody validation failures": S5 Text

**[DRAFT FOR CONSULTATION]**

**Recommendations for Antibody Manufacturers**

*Improving Antibody Validation in Biomedical Research*

Based on findings from an MRC-funded Delphi consensus study

### Executive Summary

A significant proportion of antibodies used in published biomedical research are not fit for the specific purpose for which they are used. This leads to unreliable findings, economic waste, and the unnecessary use of patient and animal samples. Technical solutions and data-sharing initiatives exist, but coordinated stakeholder action is needed to embed better practices across the research ecosystem.

Through an NC3Rs-convened stakeholder meeting and an MRC-funded Delphi study, a panel of 32 international experts — including manufacturer representatives — identified interventions to improve antibody validation that are both effective and feasible for implementation by 2030. Antibody manufacturers sit at a critical point in the research supply chain: the quality, consistency, and identifiability of reagents at source shapes what is possible downstream.

This document asks manufacturers to act on **one priority recommendation ready for implementation now**: assigning Research Resource Identifiers (RRIDs) to products at source. Two further actions — performing standard validation experiments and shifting production toward recombinant antibodies — were rated as highly effective but face implementation barriers. This document outlines how manufacturers can begin to address those.

The antibody manufacturer landscape is diverse, from large companies with extensive R&D resources to smaller suppliers and distributors. This is part of a coordinated strategy with parallel consultation documents for publishers and journals, research funders, and institutions and educational bodies. We welcome your feedback.

### About This Document

#### Background

This document is one of four stakeholder-specific consultation documents developed from an MRC-funded Delphi consensus study on antibody validation in biomedical research. A panel of 32 experts participated in two rounds of structured assessment, rating proposed interventions on effectiveness (ability to improve antibody validation in published research, scale 1–9) and feasibility (realistic implementation across the field by 2030, scale 1–9). Items achieving a median of ≥7 on both dimensions without panel disagreement are classified as consensus recommendations.

Full study methods and results are published in the accompanying manuscript, with complete qualitative commentary from panellists available in S7 and S8 Texts. Parallel consultation documents have been prepared for publishers and journals, research funders, and institutions and educational bodies.

#### Key Terminology

**Validation** refers to experimental evidence that an antibody is performing as claimed in a specific experiment — that it is interacting selectively with its intended target in the specific assay, tissue, or sample type used. Validation is context-specific: an antibody validated for Western blot in one cell type is not necessarily validated for immunofluorescence in another.

**Characterisation** refers to systematic experiments showcasing the performance qualities (or limitations) of an antibody across standardised conditions. Characterisation data (such as that generated by YCharOS or displayed on the OGA Antibody Database) can help researchers assess whether an antibody is likely to perform well, but does not replace the need for context-specific validation where results depend on antibody specificity.

### Priority Action for Manufacturers

#### Assign RRIDs to Products at Source

**Recommendation:** Antibody manufacturers should assign Research Resource Identifiers (RRIDs) to their products at source.

| **Item** | **Description** | **Effectiveness** | **Feasibility** |
| --- | --- | --- | --- |
| **A7** | Manufacturers assign RRIDs to products at source | 8.0 | 7.0 |

This achieved full consensus with high ratings on both dimensions. RRIDs enable unique identification of antibody products across the literature, supporting reproducibility and systematic tracking of antibody performance. Providing RRIDs at source removes friction for customers who increasingly need them for publication.

##### Implementation Options

1. **Self-registration with the Antibody Registry.** Manufacturers can register products directly with the Antibody Registry (antibodyregistry.org) to obtain RRIDs. Batch registration processes are available for large catalogues.
2. **Include RRIDs on product packaging and datasheets.** Printing RRIDs on product labels, datasheets, and website listings makes them immediately accessible to researchers at the point of use.
3. **Cross-reference identical clones.** Identical antibody clones sold by different vendors currently receive different RRIDs. Manufacturers could support efforts to cross-reference identical clones across suppliers, improving transparency for researchers.

### Longer-Term Actions: Building Toward Greater Impact

The following two recommendations were both rated as highly effective but face implementation barriers around cost, commercial viability, and technical constraints. For each, we suggest practical ways manufacturers can begin.

#### Perform Standard Validation Experiments

**Recommendation:** Manufacturers should perform a standard set of validation experiments on their reagent antibodies and make the data available.

| **Item** | **Description** | **Effectiveness** | **Feasibility** |
| --- | --- | --- | --- |
| **A8** | Perform standard validation experiments and make data available | 8.0 | 6.5 |

The panel agreed this would meaningfully improve antibody quality but expressed reservations about feasibility by 2030, citing cost and the challenge of defining what “standard” validation should include across diverse products. The panel also noted wide variation in current practice: some manufacturers already provide comprehensive application-specific validation data, while others provide minimal information.

##### How Manufacturers Can Begin

1. **Tiered approach.** A minimum tier (e.g., application-specific positive control data) could be feasible for most manufacturers, while enhanced validation (e.g., knockout-validated, multi-application tested) represents a premium offering. This mirrors approaches leading manufacturers already take.
2. **Reference the IWGAV framework.** Rather than creating new protocols, indicate which IWGAV-aligned validation evidence supports each product, for which applications.
3. **Make existing data more accessible.** Much validation data already exists but is not easily discoverable. Structured web pages and downloadable datasets can address this without requiring new experiments.
4. **Engage with independent benchmarking.** Initiatives such as YCharOS generate standardised characterisation data. Manufacturers can support independent testing of their products and link to the resulting data.

#### Shift Production Toward Recombinant Antibodies

**Recommendation:** Manufacturers should shift production toward recombinant antibodies.

| **Item** | **Description** | **Effectiveness** | **Feasibility** |
| --- | --- | --- | --- |
| **A9** | Shift production to recombinant antibodies | 7.0 | 5.0 |

The panel endorsed the principle — recombinant antibodies offer greater consistency, reproducibility, and lot-to-lot reliability — but identified substantial barriers to universal implementation by 2030. These include cost, the demand-driven nature of the market, and the recognition that recombinant production alone does not guarantee quality without proper characterisation.

##### How Manufacturers Can Begin

1. **Market-led transition with transparency.** Create transparency about antibody format (recombinant vs. hybridoma-derived vs. polyclonal) in product metadata. As researcher awareness grows through training, market demand will increasingly favour recombinant products.
2. **Prioritise high-impact targets.** Focus recombinant development on targets with highest research usage and where lot-to-lot variation causes the most reproducibility problems.
3. **Sequence disclosure or escrow.** Panellists suggested an escrow service that curates and stores antibody sequences, providing accession numbers so researchers can identify which products are identical — balancing transparency with commercial protection.
4. **Maintain quality standards for polyclonals.** Where polyclonal antibodies continue to be used, lot-specific characterisation and validation data should be provided. The shift toward recombinant production should complement, not replace, quality standards for existing product types.

### Support, Resources, and Next Steps

#### Stakeholder Coordination

These recommendations are part of a coordinated strategy. The Delphi panel endorsed a shared roadmap for stakeholder coordination (R22: Effectiveness 7.0, Feasibility 7.0). As publishers require RRIDs and metadata, funders endorse reporting standards, and researchers receive training, demand for well-identified, well-characterised antibodies increases. Manufacturer actions should be understood in this ecosystem context: the recommendations here are reinforced by parallel changes across the research system.

#### Recognising Good Practice

The panel repeatedly acknowledged that many leading manufacturers already meet or exceed these recommendations. We welcome engagement from manufacturers who can share their experience of implementing validation standards, RRID assignment, and recombinant antibody production. Their expertise can inform practical guidance for broader adoption across the sector.

#### Resources Available

- **Antibody Registry** (antibodyregistry.org): For RRID registration, including batch registration for large catalogues.
- **YCharOS** (<https://zenodo.org/communities/ycharos>): Open antibody characterisation data generated through independent benchmarking using knockout cell lines. Manufacturers can see how their products perform and engage with standardised, independent characterisation.
- **OGA Antibody Database** (onlygoodantibodies.co.uk): Curated, searchable interface displaying characterisation data across applications — designed to help researchers make informed antibody decisions.

#### Proposed Next Steps

We propose forming a working group to develop practical implementation guidance, including RRID registration workflows, tiered validation frameworks, and guidance on data accessibility standards. We will be working with NC3Rs and other partners to convene this.

We welcome your feedback on which implementation options are commercially realistic, barriers we have not adequately addressed, and willingness to contribute to cross-stakeholder coordination.

#### Contact

Dr Harvinder Virk

University of Leicester
