## Supplementary material for "A multi-stakeholder Delphi consensus study proposes actionable solutions to address antibody validation failures": S6 Text

**Round 1**

**Section 1: Journal and Publisher Policy**

**Reporting and Identification of Antibodies**

The following three questions explore how antibody reagents should be reported in scientific publications to support reproducibility and good practice.

Please consider the **needs and constraints of all stakeholders** (authors, reviewers, editors, publishers, readers) when assessing feasibility.

**R1 RRID Reporting**

Authors should include the **Research Resource Identifier (RRID)** for each antibody used, where available.

**R2 Metadata Reporting**

Authors should provide sufficient metadata to allow unambiguous identification of each antibody. This includes:

- Clone ID (for monoclonal antibodies)
- Catalogue number and lot number (for polyclonal antibodies)
- Vendor or source

**R3 Dilution or Concentration Reporting**

Authors should report the **concentration or dilution** at which each antibody is used. Examples include dilution ratios such as 1:1000 or 1:500, or concentrations expressed in μg/mL.

**Validation and Transparency Requirements**

The following four questions explore how publishers and journals might support more transparent, rigorous, and standardised antibody reporting and validation practices. Actions include changes to author guidelines, reviewer/editor training, validation evidence presentation, and infrastructure for linking and surfacing data quality.

Please assess these proposals from the perspective of **their ability to influence the broader research culture** while remaining **practically adoptable** by a range of journals.

**R4 Deposition of Validation Data in Open Repositories**

Journals should require authors to deposit antibody validation data in an open-access repository, linked to a persistent identifier (e.g., RRID, DOI).
Repositories might include: Zenodo, institutional repositories, manufacturer portals, or The Antibody Registry (linked to Biomed Resource Watch).

**Incentives and Visibility Measures**

**R5 Transparency Badges or Tiered Scoring**

Journals should introduce a tiered scoring system or transparency badge to recognise papers that meet higher standards of antibody reporting and validation.

**Reviewer and Editorial Capacity Building**

**R6 Reviewer and Editor Training in Antibody Validation**

Publishers should provide training resources for reviewers and editors to help them evaluate antibody validation information in submitted manuscripts.

**Technological Infrastructure**

**R7 Automated Tools to Flag Validation Concerns**

Publishers and other stakeholders should support the development and use of automated tools to flag potential antibody validation issues in submitted manuscripts – e.g., for antibodies that have already been withdrawn from the market because of performance issues.

**Validation and Specificity Evidence**

The following two questions explore how authors should report antibody validation to support confidence in the specificity, selectivity, and performance of antibody-based methods.

Please consider the diversity of experimental contexts and the varying access to validation tools across disciplines and sectors. You are encouraged to consider the role of frameworks such as the IWGAV pillars and the importance of transparency.

**R8 Validation Strategy Reporting Using IWGAV**

Authors should describe how antibody specificity was assessed, making reference to the **International Working Group on Antibody Validation (IWGAV)** framework.
A brief paragraph, checklist, or table should indicate which of the IWGAV pillars were used, if any (e.g., genetic, orthogonal, independent antibody, tagged protein, IP-MS).

**R9 Genetic Validation Evidence for Critical Applications**

Where antibodies are **critical to key experimental conclusions**, authors should either:

- present validation data based on a **genetic strategy** (e.g., knockout or knockdown), or
- cite previously published genetic validation data.

*Note: This is based on evidence from Ayoubi et al. (2023), which found that genetic validation strategies were the most predictive of antibody performance.*

**Section 2: Funders' Role**

The following nine questions consider the various ways in which research funders can promote better practices in antibody selection, validation, and reporting. This includes direct mechanisms (e.g., funding of tools and infrastructure) as well as indirect influence (e.g., signalling expectations or encouraging broader ecosystem change).

Respondents are encouraged to consider how feasible these actions would be for a typical funding body to adopt, and what impact they might have on researchers, institutions, publishers, and industry.

**R10 Funding projects that have included antibody validation**

Funders should include a section in the application form where applicants must detail the steps they will take to validate the antibodies used.

**R11 A budget for antibody validation should also be costed for in the application**

Applicants to biomedical funding schemes should include a specific line item within their budget for resources to conduct robust antibody validation (if relevant to their proposal)

**R12 Funding Development of Critical Antibody Tools**

Funders should create or expand targeted schemes to support the **development and validation of critical antibody-based research tools**. These could include, for example:

- Characterisation of under-studied antibodies
- Generation of KO/KD cell lines
- Cross-platform benchmarking of antibody performance
- Replacement of animal-derived antibodies

**R13 Signalling Value Without Mandating**

Funders should share information within the applicant guidance that antibody performance is an important limitation in many experimental methods.
Applicants should be encouraged to address this in methodological or reproducibility sections, as this will be part of how these sections are evaluated by reviewers and panels.
*Note: This would influence assessment criteria but not be a strict eligibility requirement.*

**R14 Encouraging Open Sharing of Validation Data**

Funders should encourage grantees to **deposit antibody validation data in open-access repositories**, ideally linked to RRIDs or registry entries.
This expectation could be embedded in **end-of-grant reporting** or incentivised through visibility or reproducibility initiatives.

**R15 Endorsing Journal or Community Guidelines**

Funders should formally endorse **community-developed reporting standards** that promote antibody transparency and validation (e.g., IWGAV, MDAR).
Such endorsements can help reinforce aligned expectations across funders and journals.

**R16 Encouraging Manufacturer Transparency**

Although funders do not directly regulate manufacturers, they are key drivers of demand through the research they fund. Their grant policies and collaborative initiatives can shape incentives, norms, and expectations in the commercial marketplace.

Funders should directly engage with manufacturers to encourage them to:

- Publish antibody validation datasets
- Adopt consistent metadata standards (e.g., RRID, clone ID, lot number)

**R17 Promoting Participation in Independent Benchmarking**

Funders should co-fund independent benchmarking initiatives such as YCharOS or equivalent open-science consortia by nominating key targets for their research area and contributing resources for that work.

**R18 Promoting Participation in Independent Benchmarking**

Funders should promote industry participation in independent benchmarking initiatives by asking applicants to consider working with initiatives such as YCharOS or equivalent open-science consortia as part of their work.

**Section 3: Institutions and Education**

This section explores how universities, research institutions, learned societies and training programmes can play a role in improving antibody validation and use. Possible actions include embedding training in relevant programmes, institutional policies for reproducibility, and establishing local champions.

Please consider the feasibility of implementation across different disciplines and institutional types.

**R19 Tailored Education and Training on Antibody Validation**

Training in antibody validation should be offered in relevant bioscience courses and programmes. This could include:

- Undergraduate education (e.g., in immunology, neuroscience, or molecular biology programmes)
- Postgraduate or doctoral training (e.g., through doctoral training centres)
- Postdoctoral or continuing professional development

Training should be field-appropriate, not universally required across all domains.

**R20 Research Integrity and Ethics Frameworks**

Universities and research institutions should incorporate antibody validation expectations into research integrity and ethics review frameworks. This may include:

- Asking researchers to consider antibody validation as part of research ethics approval processes
- Providing guidance or templates for documenting validation plans

**R21 Support for Local Champions of Best Practice**

Universities, research institutions and learned societies should recognise and support local champions or experts in antibody validation. These individuals may serve as:

- Points of contact for advice on antibody selection, validation design, and data interpretation
- Trainers or facilitators in reproducibility workshops or research methods courses
- Advocates for good practice within departments or core facilities
- Contributors to institutional or cross-institutional reproducibility networks

Support could include protected time, acknowledgement in performance reviews, or funding for training and outreach.

**Section 4: Cross-Stakeholder Coordination and Infrastructure**
This section addresses how coordination across stakeholder groups — including journals, funders, institutions, learned societies, manufacturers, and researchers — could enable better antibody practices at scale.

OGAs and YCharOS have been proposed as platforms for cross-sector coordination, but other initiatives or consortia may also play important roles. The intention is not to specify a single model, but to test support for shared infrastructure, road-mapping, and alignment across stakeholder types.

**R22 Establishing a Coordinated Roadmap**

Stakeholders should work together to develop a **shared roadmap** for improving antibody validation practices by 2030. This would outline:

- Stakeholder-specific actions and responsibilities
- Target timelines and milestones
- Mechanisms for accountability and feedback

**R23 Shared Infrastructure for Validation Data and Standards**

A coordinated, shared infrastructure should be developed (or expanded) to:

- Aggregate antibody validation data
- Provide standardised formats and tools for sharing
- Enable journals, funders, and institutions to access validation status or red flags

This may involve expanding existing platforms such as YCharOS, or building interoperability across initiatives like RRID, Benchsci, CiteAb, and Antibody Registry.

**Final Reflections**

**R24 Are there any other solutions to promote better antibody practices that have not been covered in the items above that you think should be included in the next round of review?**

**R25 Please use this space to share any additional thoughts, recommendations, or reflections.**

**Round 2**

Re-presented items from round 1:

**Section 1: Journal and Publisher Policy**

The following question explores how antibody reagents should be reported in scientific publications to support reproducibility and good practice.

Please consider the **needs and constraints of all stakeholders** (authors, reviewers, editors, publishers, readers) when assessing feasibility.

**R3 Dilution or Concentration Reporting**

Original wording of item R3:

‘Authors should report the **concentration or dilution** at which each antibody is used. Examples include dilution ratios such as 1:1000 or 1:500, or concentrations expressed in μg/mL.’

Revised wording of item R3:

‘Authors should report the dilution ratio at which each antibody is used. In addition, authors should state the antibody protein concentration where possible. Examples include dilution ratios such as 1:1000 or 1:500, or concentrations expressed in μg/mL.’

**Commentary:** The wording of this item has been changed to reflect comments made in round one that the dilution ratio should be the minimum level of information provided. Antibody protein concentration is very useful but is not always available.

**Validation and Transparency Requirements**

The following four questions explore how publishers and journals might support more transparent, rigorous, and standardised antibody reporting and validation practices. Actions include changes to author guidelines, reviewer/editor training, validation evidence presentation, and infrastructure for linking and surfacing data quality.

Please assess these proposals from the perspective of **their ability to influence the broader research culture** while remaining **practically adoptable** by a range of journals.

**R4 Deposition of Validation Data in Open Repositories**

Original wording of item R4:

‘Journals should require authors to deposit antibody validation data in an open-access repository, linked to a persistent identifier (e.g., RRID, DOI).
Repositories might include: Zenodo, institutional repositories, manufacturer portals, or The Antibody Registry (linked to Biomed Resource Watch).’

**Commentary:** We ask you to re-rate this item because the aggregate feasibility rating was ‘Unsure’.

In round one participants made the following comments regarding this item:

- Online depositories can be confusing, time-consuming, or challenging
- For the data to be useful, someone would need to define what validation data was required
- For this to be useful, someone would need to verify the quality of the validation data that was deposited
- This item may be a barrier to authors at the submission stage of manuscript publication.

Several participants commented that validation data would be better stored elsewhere. Two options were suggested; in/with the publication or on manufacturers websites. We have added separate items for these suggestions below. Please rate this item independently of whether validation data is included in the publication and/or on the manufacturers’ website.

**Incentives and Visibility Measures**

**R5 Transparency Badges or Tiered Scoring**

Original wording of item R5:

‘Journals should introduce a tiered scoring system or transparency badge to recognise papers that meet higher standards of antibody reporting and validation.’

**Commentary:** This item is being included in round two without amendment. In round one participants made the following comments regarding a tiered scoring system:

- The journal should set the standard for all manuscripts accepted; no need for an additional badge
- To be effective it would need standard criteria across journals, which may be difficult to implement
- Who would administer it?
- Validation requirements are different according to the experimental situation, so scoring would be complicated
- A tiered approach may be too light and not influential enough
- Other badging schemes (eg Open Science Indicators) have had limited success
- There is no incentive to participate in the current research climate

Taking these into consideration, please re-rate the item.

**Reviewer and Editorial Capacity Building**

**R6 Reviewer and Editor Training in Antibody Validation**

Original wording of item R6:

‘Publishers should provide training resources for reviewers and editors to help them evaluate antibody validation information in submitted manuscripts.’

**Commentary:** Participants are asked to re-rate this item because there was disagreement amongst participants in round one. To provide further context on this item, validation training resources are being prepared on antibody validation by Only Good Antibodies and will be made freely available.

**Technological Infrastructure**

**R7 Automated Tools to Flag Validation Concerns**

Original wording of item R7:

‘Publishers and other stakeholders should support the development and use of automated tools to flag potential antibody validation issues in submitted manuscripts – e.g., for antibodies that have already been withdrawn from the market because of performance issues.’

**Commentary:** This item is being included in round two without amendment. In round one participants made the following comments:

- Some participants rated this highly for feasibility because similar tools (e.g., SciScore) already exist
- Cost may be a limiting factor for some publishers.
- This may only be feasible if RRIDs were applied at a supplier level.
- This would need someone to administer it and keep it up to date
- Images would need to be checked by a human
- How would a bad antibody be linked back to prior publications and would they need to be retracted?

**Validation and Specificity Evidence**

The following two questions explore how authors should report antibody validation to support confidence in the specificity, selectivity, and performance of antibody-based methods.

Please consider the diversity of experimental contexts and the varying access to validation tools across disciplines and sectors. You are encouraged to consider the role of frameworks such as the IWGAV pillars and the importance of transparency.

**R8 Validation Strategy Reporting Using IWGAV**

Original wording of item R8:

‘Authors should describe how antibody specificity was assessed, making reference to the **International Working Group on Antibody Validation (IWGAV)** framework.
A brief paragraph, checklist, or table should indicate which of the IWGAV pillars were used, if any (e.g., genetic, orthogonal, independent antibody, tagged protein, IP-MS).’

Revised wording of item R8:

As part of the submission process, authors should describe how antibody specificity was assessed, making reference to the **International Working Group on Antibody Validation (IWGAV)** framework.
A brief paragraph, checklist, or table should indicate which of the IWGAV pillars were used, if any (e.g., genetic, orthogonal, independent antibody, tagged protein, IP-MS).

**Commentary:** This item has been rephrased to clarify that this would be part of the submission process, not included in either the body of the manuscript or a supplementary file (there has been an additional item included in this round to address inclusion of data in publications).

In addition, participants made the following comments in round one:

- Authors may not understand what is being asked of them, either in how to complete the check list or how to complete the experiments
- Author compliance could be an issue
- Is implementation at the point of publication too late – authors won’t be aware and won’t go back to complete the validation if they haven’t already done it
- Are the IWGAV recommendations universally accepted?
- Most labs aren’t equipped to complete validations. This may need a consortium approach

**R9 Genetic Validation Evidence for Critical Applications**

Original wording of item R9:

‘Where antibodies are **critical to key experimental conclusions**, authors should either:

- present validation data based on a **genetic strategy** (e.g., knockout or knockdown), or
- cite previously published genetic validation data.’

Revised wording of item R9:

‘Where antibodies are **critical to key experimental conclusions**, where technically feasible, authors should either:

- present validation data based on a **genetic strategy** (e.g., knockout or knockdown), or
- cite previously published genetic validation data.’

**Commentary:** This item has been rephrased in light of participant comments from round one highlighting that genetic strategies may not be feasible in certain model systems or for certain epitopes.

**Section 2: Funders' Role**

The following questions consider the various ways in which research funders can promote better practices in antibody selection, validation, and reporting. This includes direct mechanisms (e.g., funding of tools and infrastructure) as well as indirect influence (e.g., signalling expectations or encouraging broader ecosystem change).

Respondents are encouraged to consider how feasible these actions would be for a typical funding body to adopt, and what impact they might have on researchers, institutions, publishers, and industry.

**R16 Encouraging Manufacturer Transparency**

Original wording of item R16:

‘Although funders do not directly regulate manufacturers, they are key drivers of demand through the research they fund. Their grant policies and collaborative initiatives can shape incentives, norms, and expectations in the commercial marketplace.

Funders should directly engage with manufacturers to encourage them to:

- Publish antibody validation datasets
- Adopt consistent metadata standards (e.g., RRID, clone ID, lot number)’

**Commentary:** Participants are asked to re-rate this item because there was disagreement amongst participants in round one. Several comments were made on the ability of manufacturers to provide validation data and Item A8 has been included in round two to address this. Therefore, when rating this item, please only consider this from the point of view of the Funder.

In addition, participants made the following comments:

- Funders don’t have time for this
- The funding situation (at least in the US) would make this difficult for federal agencies
- This is not in the scope of most funders work
- It would be difficult for funders to find points of contact with the manufacturers
- Why would the manufacturer be influenced by the funder?

**R17 Promoting Participation in Independent Benchmarking**

Original wording of item R17:

‘Funders should co-fund independent benchmarking initiatives such as YCharOS or equivalent open-science consortia by nominating key targets for their research area and contributing resources for that work.’

Revised wording of item R17:

‘Funders should collaborate with independent benchmarking initiatives such as YCharOS or equivalent open-science consortia. They should directly co-fund the validation of antibodies against key targets to create suites of validated tools that can be used by researchers to improve data quality and thereby increase return on investment for the funder.’

**Commentary:** This item has been rephrased for clarity.

**R18 Promoting Participation in Independent Benchmarking**

Original wording of item R18:

‘Funders should promote industry participation in independent benchmarking initiatives by asking applicants to consider working with initiatives such as YCharOS or equivalent open-science consortia as part of their work.’

Revised wording of item R18:

‘Funders should promote industry participation in independent benchmarking initiatives by asking applicants to consider working with initiatives such as YCharOS or equivalent open-science consortia as part of their work. This could be achieved either by including the recommendation in the guidance notes or flagging to applicants at the panel review stage.’

**Commentary:** This item has been reworded to include a more detailed description of how this may work in practice.

**Section 3: Institutions and Education**

This section explores how universities, research institutions, learned societies and training programmes can play a role in improving antibody validation and use. Possible actions include embedding training in relevant programmes, institutional policies for reproducibility, and establishing local champions.

Please consider the feasibility of implementation across different disciplines and institutional types.

**R21 Support for Local Champions of Best Practice**

Original wording of item R21:

Universities, research institutions and learned societies should recognise and support local champions or experts in antibody validation. These individuals may serve as:

- Points of contact for advice on antibody selection, validation design, and data interpretation
- Trainers or facilitators in reproducibility workshops or research methods courses
- Advocates for good practice within departments or core facilities
- Contributors to institutional or cross-institutional reproducibility networks

Support could include protected time, acknowledgement in performance reviews, or funding for training and outreach.

Revised wording of item R21:

Universities, research institutions and learned societies should recognise and support the Antibody Champions Scheme. This UK-based initiative, co-funded by the Medical Research Council (MRC) and the National Centre for the replacement, refinement and reduction of animals in research (NC3Rs), will be launched in 2026. It will be a voluntary scheme of local champions or experts in antibody validation. These individuals will serve as:

- Points of contact for advice on antibody selection, validation design, and data interpretation
- Trainers or facilitators in antibody validation best practice workshops or research methods courses
  - Advocates for antibody good practice within departments or core facilities

Support could include protected time, acknowledgement in performance reviews, or funding for training and outreach.

**Commentary:** This item has been rephrased for clarity.

**Section 4: Cross-Stakeholder Coordination and Infrastructure**

This section addresses how coordination across stakeholder groups — including journals, funders, institutions, learned societies, manufacturers, and researchers — could enable better antibody practices at scale.

OGAs and YCharOS have been proposed as platforms for cross-sector coordination, but other initiatives or consortia may also play important roles. The intention is not to specify a single model, but to test support for shared infrastructure, road-mapping, and alignment across stakeholder types.

**R23 Shared Infrastructure for Validation Data and Standards- share comments and ask for suggested solutions**

‘A coordinated, shared infrastructure should be developed (or expanded) to:

- Aggregate antibody validation data
- Provide standardised formats and tools for sharing
- Enable journals, funders, and institutions to access validation status or red flags

This may involve expanding existing platforms such as YCharOS, or building interoperability across initiatives like RRID, Benchsci, CiteAb, and Antibody Registry.’

**Commentary:** This item is being included in round two without amendment. In round one participants made the following comments:

- This would be a huge undertaking
- This would need sustainable funding
- This would need a system of community-based governance to be implemented

**Additional Items**

The following items were suggested by participants in round one. Please rate them for effectiveness and feasibility according to the same criteria as the previous items.

**A1** Journals should have a clearly stated standard for antibody validation and reporting

**A2** Detailed Protocols and Validation data should be included in the manuscript or supplemental data

**A3** Journals should appoint a specialist editor that focuses on reproducibility

**A4** Funders should maintain and provide researchers with a list of manufacturers who validate their antibodies

**A5** Funders should mandate the use of recombinant antibodies (or at least monoclonals) when available. Justification should be required if a polyclonal is used.

**A6** Funders should mandate that if funds are used to develop an antibody as part of a research project, the reagent must be a recombinant antibody and must be made available to the wider community.

**A7** Antibody manufacturers should be encouraged to assign RRIDs to their products at source

**A8** Manufacturers should perform a standard set of validation experiments on their reagent antibodies and make the data available

**A9** Manufacturers should shift production to all recombinant antibodies.**A10** A learned society for antibody validation should be established. This could perform several functions, for example:

a. Promote antibody validation
b. Provide training and guidance on antibody validation strategies
c. Provide standard protocols for validation approaches
d. Provide independent assessment of antibody validation data
e. Oversee a tiered scoring/badge system for publications containing antibody data (see Q5)
f. Work with manufacturers to improve antibody transparency and validation
