## Supplementary material for "A multi-stakeholder Delphi consensus study proposes actionable solutions to address antibody validation failures": S7 Text

**S7 Text. Round 1 qualitative feedback (participant comments).**

*Free-text comments provided by Delphi panellists during Round 1 (n = 32 participants, 23 items). Comments are presented verbatim and organised by item. Comments are numbered sequentially within each item; numbering restarts for each item and does not track individual participants across items.*

**Item R1**

*RRID Reporting. Authors should include the Research Resource Identifier (RRID) for each antibody used, where available.*

**Participant Comments**

**[Comment 1]**

Effectiveness and feasibility has been demonstrated empirically

**[Comment 2]**

Barriers are understanding how RRIDs are generated, but is should be feasible to explain this at the journal interface. Not sure if this is problematic if there are issues with batch to batch variation for antibodies with the same RRID - how would this apply to polyclonal antibodies?

**[Comment 3]**

Could be effective in making results reproducible, but those results won't necessarily be valid.

**[Comment 4]**

I believe that the reporting of RRIDs by authors would improve research reproducibility. I interpret feasibility as the extent to which journals can effectively implement this policy.

In my view, feasibility is constrained by several factors: author behavior (which often does not prioritise reproducibility); technological limitations on the publisher's side, (particularly regarding cost-effective compliance checks); lack of standardisation of this policy across most journals.

By 2030, I anticipate that technological advancements will make it possible for journals to implement RRID reporting as a mandatory requirement, although standarisation by most journals will also be key. I am assigning a feasibility score of 6, as I believe the primary barrier is author behaviour, as this is shaped by incentive structures that currently do not reward efforts (ie. by funders) toward reproducibility or transparent reporting.

**[Comment 5]**

I rated the effectiveness as 7 because RRIDs are not perfect. The same antibody sold by different providers can have different RRIDs leading to researchers to falsely believe 2 antibodies are different when they are, in fact, the same. This is problematic as researchers often use "different" antibodies to cross validate each other and could in theory be comparing an antibody to itself for validation purposes. This approach should be completely feasible and not a huge hurdle for researchers.

**[Comment 6]**

RRID allows you to assign an identifier if one is not available. Plus, antibody producers could be encouraged to assign RRIDs to their products at source. Down to individual batches if possible. That could complete the identification chain for the reagents being used.

**[Comment 7]**

An issue is that an RRID entry might only list one commercial supplier of the antibody, so accurate tracking of reagents across multiple suppliers is lost. Also, while citing RRIDs in publications would hopefully provide more clarity on the reagent used, oftentimes a clone number/name can be a more useful identifying feature and also help with cross-referencing other publications and commercial supplier data. And RRID does not speak to antibody validation or quality, or variation in quality across companies that use different production/purification methods.

**[Comment 8]**

The problem with RRIDs is that they take no account of 1) lots and 2) identical antibodies sold by different companies. In my opinion this makes them pretty useless, unless these issues are addressed. Addressing both of these issues would make RRIDs much much more valuable.

**[Comment 9]**

Of course, RRIDs work well for monoclonal antibodies, but are not effective for polys (batch to batch variation)... and polys remain the most used antibody product.

**[Comment 10]**

RRIDs do not contemplate the same clone sold by multiple vendors. There is a cost associated with the bulk assignment of RRIDs which is a barrier to implementation broadly. How does the RRID differ from the combination of vendor name and product number? Either could work as an identifier if the requirements around reagent identification were standardized and enforced by the journals. RRIDs create another layer of bureaucracy.

**[Comment 11]**

Publishing mandates are effective but often hard to police. Also, they can feel like unnecessary bureaucracy for authors so providing incentives and case studies for the benefits would assist

**[Comment 12]**

Although this is feasible in theory, enforcement by journals would require that the article is checked to ensure the RRIDs are there. This could potentially be automated but the automation would need to be put into the screening checks andd cost for smaller or scholar-led publishers may be prohibitive.

**[Comment 13]**

I have never used RRID as a resource therefore cannot rate the effectiveness

**[Comment 14]**

RRIDs are company and product form specific, so no guarantee that the mAb with the exact RRID will be available in the future

**[Comment 15]**

RRIDs are theoretically good at identifying unique antibodies from suppliers but there are some caveats: 1. they do not take lot numbers into account and 2. they rely on the suppliers keep the same catalog numbers with the same antibodies. RRIDs should work better for recombinant antibodies however.

**Item R2**

*Metadata Reporting. Authors should provide sufficient metadata to allow unambiguous identification of each antibody. This includes: clone ID (for monoclonal antibodies), catalogue number and supplier name, lot/batch number (where available), host species and clonality.*

**Participant Comments**

**[Comment 1]**

Providing the information listed should be the minimal standard, with use of RRID and any other validation data even better.

**[Comment 2]**

Authors don't know which items to add without a clear unambiguous guideline and with a guideline they often partially comply

**[Comment 3]**

Whilst providing metadata is good, it's not so clearcut how easy it is for readers to identify the resource described. Using some recognised standard identifiers would help e.g. RRIDs.

**[Comment 4]**

There would need to be a change of culture in organisations at the stage of data capture, as this might be overlooked and result in inaccurate reporting (i.e. pick an unrelated batch # if you don't know the real one?). What about studies where multiple batches have been used - the authors may not want to disclose this?

**[Comment 5]**

As above.

**[Comment 6]**

In my opinion, this combined with RRIDs is one step better. You could run into the same issue as above though if a company acquired an antibody and then changed its name/clone number. I'm not sure how often that actually occurs but unless antibody providers become completely open about the history of their antibodies (original source, changes in name, changes in format etc.) or provide antibody sequence users don't know exactly what they are using. From a feasibility perspective this is completely feasible.

**[Comment 7]**

Some producers rename the clones. And some clone names are duplicated in wholly different antibodies. So they may be inadvertently "soft" identifiers, and ambiguous. This measure is likely only effective if associated with a hard identifier like an RRID. I am an absolutist, and very resistant to the use of polyclonals in research, given the current excellent alternatives. But, in short, even with extensive metadata on a polyclonal, so what? The next batch will be different. The likelyhood of the next batch being different sinks when using a monoclonal, and is a lot lower for recombinants.

**[Comment 8]**

This should be a minimum reporting requirement. Allows tracking of clones across suppliers (via clone ID) for monoclonal antibodies and accounts for differences in material between suppliers and lots. Everyone should have this information already in their lab notebooks or can look at through purchasing history.

**[Comment 9]**

The challenge with the reporting vednors is there are multiple antibodies that are the same but have different catalog numbers/suppliers due to the OEM from different suppliers.

**[Comment 10]**

Will require buy-in and enforcement from journals.

**[Comment 11]**

Space to include this information in manuscripts and grant applications can be limited, so this would need to addressed with publishers and funders to enable this change

**[Comment 12]**

Metadata reporting would, in theory, help improve reproducibility but such detailed reporting could also represent an additional burden for authors. Additionally, if antibodies have been developed in-house there may be IP related barriers to providing very granular information.

**[Comment 13]**

Some vendors do not provide unambiguous production lot numbers

**[Comment 14]**

form of antibody should also be reported (supernatant, purified, concentrated, affinity purified, etc..)

**Item R3**

*Dilution or Concentration Reporting. Authors should report the concentration or dilution at which each antibody is used. Examples include dilution ratios such as 1:1000 or 1:500, or concentrations expressed in µg/mL.*

**Participant Comments**

**[Comment 1]**

If protein concentration is available it should definitely be included. Experience tells me that use of dilution can be misleading or erroneous as my lab typically dilutes incoming commerical Abs 1:1 with glycerol for storage at -20C but not all lab members take this into account in listing dilutions. Also, it is quite common to not know the protein concentration of an Ab, e.g., if using media in which hybridoma cells were grown as a source.

**[Comment 2]**

Methods should be described a fully a possible to allow interested researchers reproduce the work.

**[Comment 3]**

Scored highly because this will be effective, but has some limitations. This will only be truly effective if the actual antibody concentration (rather than dilution) is specified. In my opinion, the supplier should always specify the actual concentration, and suggest working concentrations rather than dilutions. Suggesting dilutions leads to non reproducibility due to changes in the batch concentration that the end user may miss by mistake

**[Comment 4]**

As above.

**[Comment 5]**

Concentration alone, while helpful, is not enough. Full detailed protocols should be provided including processing steps, buffer recipes, incubation times, temps etc. If researchers started providing more details I think it would be hugely beneficial for reproducibility efforts but I also think that there will be resistance. It will take some time to pull the information together especially when the person submitting the manuscript (Professor) is not the individual who performed the experiment.

**[Comment 6]**

If you are forced to use dilutions as a measure, then you do not know how much antibody is actually present in your reagent. Of course, for polyclonals there is no good alternative. But producers should be encouraged to state unequivocally the antibody protein concentration in the reagents they sell. And the concentraion (or dilution) data must be stated by users for every application in a study.

**[Comment 7]**

Again, this should be information stored in lab notebooks/protocols. Easy to incorporate and informs future use and interpretation of results.

**[Comment 8]**

Antibodies should be described in terms of concentration - µg/ml. Dilutions from unknown starting concentrations (except polyclonals) is useless

**[Comment 9]**

Following the manufacturers recommended dilution ratio will help optimize performance of the antibody; concentrations can vary and it would be incorrect to assume that a low concentration would result in low activity.

**[Comment 10]**

Dilutions can be meaningful when tied to a specific lot of antibody, but a dilution in the absence of a reported lot number may not be helpful. Additionally, optimal concentrations are helpful, of course, but can be influenced by various experimental parameters. Laboratories should be determining optimal working conditions themselves and not simply using an antibody blindly.

**[Comment 11]**

Again, requiring such things at the stage of publication may be too late. Also limits on pages numbers of submission can cause author to remove details to fit constraints. Ideally these requirements should be made further upstream to become community norms

**[Comment 12]**

Concentration proffered over dilution. If dilution given, then antibody concentration should be listed somewhere.

**[Comment 13]**

Dilutions should reported along with concentration in ug/mL or other units. Concentration of Ab should be independently confirmed upon receipt. This may not be feasible when unpurified, provided in ascites, or with additives in the formulation that interfere with BCA or UV analysis.

**[Comment 14]**

monoclonal antibodies really should always be reported as concentrations expressed in μg/mL owing to different preparations by different manufacturers

**Item R4**

*Deposition of Validation Data in Open Repositories. Journals should require authors to deposit antibody validation data in an open-access repository, linked to a persistent identifier (e.g., RRID, DOI).*

**Participant Comments**

**[Comment 1]**

I have found that use of some of the online resources can be confusing, time consuming or challenging, which is why my score for feasibility is lower than it might be otherwise

**[Comment 2]**

Does not seem feasible, because such repositories are unknown. People can use supplemental files giving them a DOI, they can use zenodo, but those are options without guidance for what to report and how to do it.

**[Comment 3]**

How do we judge the reliability of antibody validation data?

**[Comment 4]**

While it could be straight forwards to mandate that an identifier is linked to each cited antibody, and could potentially be an effective strategy, it is dependent on the quality of the validation data. If this needs to be check by editorial staff, then I think it will be infeasible. Also, data of this sort should ideally be reviewed by experts (ie researchers).

**[Comment 5]**

I marked it as 4 for feasibility because I see this as a bigger barrier to authors at submission (rather than reporting the RRID alone), although I agree that is is necessary for reproducibility. This is based on my knowledge of authors compliance with depositing their raw data. I would also expect peer reviewers to query validation of all newly used antibodies as part of the peer review process.

**[Comment 6]**

Validation data would be extremely effective. I think there could be resistance given the amount of work it requires (similar to my concerns for Q3.) In addition to providing the validation data, the validation would need to be reviewed to ensure that the experiment was carried out properly with proper controls. Would this be part of the peer review process?

**[Comment 7]**

Unless the validation protocols are to be standardized, the power of depositing the data on validation will be uncontrolled, on the GIGO principle. If there are at some time defined standard validation methods, then it would be a good approach. The lack of a centralized place where the validation data might reside (analogous to PubMed) might be an issue. Unless someone gathers the validations...

**[Comment 8]**

Agree with this standard, although more guidance should be provided - eg does the validation data need to be from the current lot of antibody used for the study?

**[Comment 9]**

How do you determine that validation for the antibody was required to use? Or part of the experiment related to the publication? There is also a concern this eventually will not be an open and free access.

**[Comment 10]**

Issues can come with softwares required to open specialized figure formats deposited on repositories (mostly true for microscopy images, but might be true for immunoblot images)

**[Comment 11]**

Journal do sometimes ask for some validation data. I do believe in having access to the validation data and I believe that having the data in manufacturer portals (product websites) is the easiest and most direct way to access the data.

**[Comment 12]**

Validation data should be included in manuscripts, even if as supplemental data.

**[Comment 13]**

This is a very feasible option, but requires a behavior change from researchers driven by funders, journals and institutions to make this an expectation of their grantees, those submitting manuscripts and their researchers

**[Comment 14]**

Depositing data is highly feasible if the data is available, however I would not anticipate this being available for the majority of antibodies currently.

**[Comment 15]**

Generalist repositories often do not require or provide fields for specificity of metadata required and so this data ends up available and open but only if you know where to look, which reduces it's over all impact

**[Comment 16]**

As for any data sharing requirement, the problem with feasibility is for large multidisplinary journals to enforce the requirement at scale or publishers to enforce across a portfolio. Even where there are mandatory data access statements, enforcement currently entails that journal staff manually check that the data is appropriately deposited, which is not scaleable (e.g. for large multidisciplinary journals). This might become more automated in the future.

**[Comment 17]**

Would be an ideal practice, however, impractical to expect across all journals. in addition, the primary validation that might have been deposited for one journal does not guarantee that the data shown in a subsequent manuscript is based on a lot with comparable performance.

**Item R5**

*Transparency Badges or Tiered Scoring. Journals should introduce a tiered scoring system or transparency badge to recognise papers that meet higher standards of antibody reporting and validation.*

**Participant Comments**

**[Comment 1]**

If I understand this proposal correctly, it is about rewarding a paper in a journal with a high score for transparency. I believe that instead that journal should have a clearly stated standard for antibody validation and reporting and hold to that standard so that all papers published will be understood to meet that standard. Further rewards are not needed.

**[Comment 2]**

how would badging systems be sold to journals (badges on top of the journal were implemented by TOP guidelines and I have not seen many of those)? How will they be implemented? - every paper would need to have a badge? every antibody paper would need to have a badge? If so, the editors would need to give badges which they may not have the expertise to do. Managing Editor often = English major; Academic Editor is usually too busy for daily tasks

**[Comment 3]**

I think this approach is too light and may not be influential enough - particularly if broad uptake is low across the community such that if nobody bothers to go the extra mile, then it will be self fulfilling

**[Comment 4]**

Several papers have use only one or two antibodies for as part of established assays, so I don't see the point of putting a badge on those. Potentially I could see this working as an opt-in system for authors of antibody heavy papers, where an author who chooses to include validation for their antibodies can opts in to having the quality of the validation scored. If the badge can be incentivized eg papers that are recognized as having well validated antibodies are heavily cited, this could drive uptake. However, the scoring would have to be an external system used across the journals.

**[Comment 5]**

I marked these rather low because I feel that badging per se (for instance for open science indicators) has not worked as well as intended, because the incentives are not there. Once research assessment changes, away form the IF toward reproducibility, dissemination of reliable science, badging may become more effective in promoting better reporting. Another barrier to making badging effective, is the lack of standardisation by different journals. Without a common grading system, comparison across different journals would be difficult.

**[Comment 6]**

I have several concerns about a system like this mainly 1. Is this part of the peer review process? Peer reviewers would need to assess antibody validation claims but may lack the expertise for a proper review. 2. How is the standardized criteria determined and is it identical across journals? Unless there is a benchmark “scoring” may be uneven or meaningless. 3. What counts as “validated”? Western blot against knockout tissue? CRISPR knockout cell lines? Peptide blocking? Different fields value different validation metrics, and forcing a one-size-fits-all rule could alienate authors. 4. If a badge becomes a mark of prestige, authors may game the system, cherry-picking validation data, over-claiming reproducibility, or leaning on vendor-provided data without independent verification.

**[Comment 7]**

Quis custodiet ipsos custodes? ( I had to look it up!) Who will judge how effectively the journals can judge the data presented to them?

**[Comment 8]**

would be difficult to define the Tiers

**[Comment 9]**

While this might be useful, if we start getting into transparency badges/tiered scoring for each aspect of validation that a study should require, then meaning is lost. For instance, this should also probably be used for statistical methods, animal use, figure quality checks (for fabrication/duplication) etc. Also, what is the scoring system - is it subjective or objective? Is it done by a neutral body across journals or does each journal have to establish their own system?

**[Comment 10]**

It will be difficult to enforce this among all publications consistently - especially ones who don't enforce standards already. Star methods already exists. It could also be seen as rating the quality of the publication or the science behind the publication.

**[Comment 11]**

Standards needs to be figured out to evaluate "transparency". And same standards per journals might be difficult. I think reviewers providing their thoughts of reproducibility of the experiments would be necessary.

**[Comment 12]**

A scoring system may be a distraction and take away from the the Journal's focus on the message/content of a given paper.

**[Comment 13]**

This is a feasible change, but beyond it being a visual indication that higher standards have been met, if this is not linked to a stronger incentive, i don't see it being particularly effective on its own

**[Comment 14]**

Potential to be effective but only if badges are of interest to the research community.

**[Comment 15]**

Unless all journals do this in a consistent and comparable manner it loses its effectiveness. Journals struggle to agree on consensus and variability makes it hard to really assess articles cross journals for their openness. Also, journal QC requirements will increase and the level of QC will be variable - journals may require certain things, or even mandate, but if they don't QC they might not be confirming to checking the standards they set themselves

**[Comment 16]**

I think this could only work if automated although badges themselves do not increase reproducibility and could also potentially be gamed

**[Comment 17]**

I think it will be challenging to define standards for all applications. Some people may not have funds to validate reagents and so may be penalized - particularly if their paper contains many antibodies.

**[Comment 18]**

This would require developing criteria that would have to be agreed upon amongst journals which could be challenging. Unless there is a clear incentive or benefit to gaining 'transparency badges' (e.g. featuring/highlighting higher standard papers) that approach may not be enough to boost rigour of antibody validation and reporting.

**[Comment 19]**

Not sure that a "tiered" scoring process would result in improvements in antibody validation/rigour, rather, it would direct authors/manuscripts toward journals that are willing to publish under less restrictive criteria. Not really different than the current climate.

**Item R6**

*Reviewer and Editor Training in Antibody Validation. Publishers should provide training resources for reviewers and editors to help them evaluate antibody validation information in submitted manuscripts.*

**Participant Comments**

**[Comment 1]**

In my experience as a reviewer of grants and papers, there is so much work involved in evaluating/reviewing the science that reviewers make short work of answering questions about such things as antibody validation. I believe the journal should take control of this issue to result in a far more consistent and impactful outcome.

**[Comment 2]**

This would be nice but editors may not have the slightest idea what an antibody is and why it is important. They would need to read the paper and they usually don't. Reviewer training is more feasible especially if it comes in the format of a checklist that they will need to fill out.

**[Comment 3]**

Editors and reviewers might not have the necessary expertise to evaluate reports.

**[Comment 4]**

Potential barriers - this will require a standardised approach to make this consistent across editors and between different journals. They would all need to meet this standard to avoid individuals "shopping around" for those with the easiest criteria to fulfil

**[Comment 5]**

While thorough assessment by reviewers or checking by editors could be a very effective way to evaluate the submission of validation data, I think currently it is not feasible. Reviewers will already be spending time reviewing the scientific content of the paper, so I think it is unrealistic to ask them to further review the validity of the data. This may work if technical reviewers are invited specifically for this purpose, but then there are questions as to which papers are put forward for a technical review of antibody data. It is currently also unrealistic for editors to spend time checking antibody validation, unless there is a very simple system in place.

**[Comment 6]**

I marked reviewer and editor training as a 7 for effectiveness in improving reproducibility but much lower for feasibility because I feel that journals are not best placed for training reviewers and editors adequately. I feel that this type of training should be conducted by institutions as part of the standard training for PhD students (maybe in collaboration with guidance from publishers) or organisations dedicated to improving peer review.

**[Comment 7]**

I highly doubt that any PhD- or MD-level scientist would invest time in learning more about how to tell if an antibody is "good" or not in a manuscript. I think the vast majority of researchers already consider themselves experts if they worked with antibodies during their training.

**[Comment 8]**

I think ultimately it should land on the editors. Anyone who has had papers reviewed knows that some reviewers are thorough while others glean over and provide next to no feedback. Relying on reviewers will inevitable end with some papers having a thorough validation review while others do not. I think the best path would be to have a dedicated role in the journal review/editing process for reagent validation.

**[Comment 9]**

It would depend on perhaps unenthusiastic and time-pressed reviewers and editors using the resources effectively.

**[Comment 10]**

would be difficult to provide education for reviewers and discourage people from participating as reviewers

**[Comment 11]**

This is a subjective assessment and most reviewers are likely to ignore training resources to spare time when they're already reviewing for journals in addition to their day job.

**[Comment 12]**

Assume is they are reviewing they are qualified to scientifically review the publication.

**[Comment 13]**

Exactly, I like this a lot

**[Comment 14]**

This is important, but key to effectiveness is consistent application across reviewers and editors. I think a more effective approach would be for journals to appoint a specialist editor that focuses on issues of reprehensibility, in the same way that some focus on animal welfare considerations

**[Comment 15]**

Providing training resources does not guarantee that these will be used. Could be effective if journals go beyond training and insist that (a) an evaluation of antibody validation is provided with every review, and (b) validation information must be included for publication.

**[Comment 16]**

Again, reviewer tasks are already onerous and unless there is some assistance in checking this automatically it is unlikely that reviewers will be able to do this within their reviewing capactiy

**[Comment 17]**

training resources themselves are unlikely to help at this stage. They would need to be targeted to make them available to a reviewer of a relevant paper, which would require that the paper is appropriately flagged in the system. Generic resources available on e.g. a journal website are unlikely to be read. Enforcing training across large reviewer pools is probably not feasible. Editors of journals that receive a large number of relevant submissions might be incentivized but this is unlikely to be the case for large multidisciplinary journals. Guest Editors of e.g. special issues might also lack awareness or incentives. In principle resources could e.g. perhaps be part of the submission guidelines for relevant special issues but that would also require bespoke changes to journal workflows. Training really needs to happen before the paper gets published and ideally before the researchers do the study. Publishers and journals can help with providing resources but this will only be effective I think if institutions/ scholarly societies/ National Academies also co-ordinate/collaborate so that the training and messaging at all stages of the research cycle are aligned.

**[Comment 18]**

Providing guidelines would be useful but I think it would be difficult to implement a training program given the number of reviewers. Prefer an automated AI tool.

**[Comment 19]**

Training and education at all levels is key. It should start at one's earliest introduction to bench research. It is an unfortunate reality that researchers can advance their careers, notoriety, and influence and still have only a rudimentary understanding of antibody validation, QC, and protocols.

**[Comment 20]**

very little way to enforce this

**Item R7**

*Automated Tools to Flag Validation Concerns. Publishers and other stakeholders should support the development and use of automated tools to flag potential antibody validation issues in submitted manuscripts.*

**Participant Comments**

**[Comment 1]**

I have had the pleasure of testing out SciCrunch (I think that is the name of the program) and found it remarkably effective and a version of such software should absolutely be in place at all reputable journals.

**[Comment 2]**

see PMC12338114

**[Comment 3]**

Whilst automated tools would be good and reduce the burden on editorial office, editors and reviewers, not all publishers/journals have the financial means to pay for such services or tools.

**[Comment 4]**

I don't know if this is technically possible, it may depend on the prior implementation of RRIDs, and also the RRIDs being applied at the supplier level in order to mine the correct information?

**[Comment 5]**

If a tool can be developed which can flag up problematic manuscript, then this could be effective, as editors will know which manuscripts there are concerns about, and in these specific cases, they could be flagged up to reviewers.

**[Comment 6]**

Journals would benefit from the development and implementation of these tools, this is why I marked it as an 8 on the effectiveness of improving the rigour of antibody-based method reporting. I am not aware of any tools that could reliable flag these (hence the 6), but it seems that if/when available journals could use them to facilitate the review process and alert editors/reviewers of technical concerns with the reagents that could potentially invalidate the conclusions reported.

**[Comment 7]**

Someone has to develop and refine the software.

**[Comment 8]**

Automated tools can flag missing metadata or inconsistent reporting of antibodies, but they struggle with vague terminology, incomplete vendor data, and field-specific validation standards. At best they act like a spell-checker for reporting, not a true judge of scientific validity and risk being gamed or ignored if journals and funders don’t back them with real incentives.

**[Comment 9]**

It would be a good idea, but feasibility depends on an RRID or other hard identifier of the antibody in question. And I wonder how "performance issues" would be universally judged.

**[Comment 10]**

I like the specific idea of flagging antibodies that have been withdrawn from the market due to poor performance. The effectiveness of the tool would really rely on the data the tool is drawing from. Especially since differences in methods across groups could result in some groups seeing specificity of signal while others do not.

**[Comment 11]**

It feels like this tool (flagging withdrawn abs) should be available for authors prior to doing all of the work. If it is public the journal reviewers would also likely use it. It is also likely a fraction of the issues.

**[Comment 12]**

I would need to see the performance of such tools before giving any rating

**[Comment 13]**

Most manufacturers do not state why an antibody was withdrawn from the market; the automatic assumption should not be "performance issues." Antibodies can be pulled from the market for any number of reasons including supply, low revenue, poly to mono projects, and specificity among others. An automated flag would be unfair to the publication and antibody without determining the history of the product.

**[Comment 14]**

Antibodies can be removed from the market for reasons other than specificity concerns. There can be manufacturing issues, for example.

**[Comment 15]**

A great option, but who will fund the development of such a tool and maintain it so it is always up to date

**[Comment 16]**

This is the way forward, and it is possible. But requires funding and uptake by journals to implement, which could be a blocker as their internal systems are probably inflexible.

**[Comment 17]**

Feasibility will vary for different publishers and/or journals and depend on the cost of developing the automation or buying third party tools. SciScore is one such tool (for reproducibility more generally) but it's hard to get this prioritized at e.g. a large publisher when there are many other competing priorities and is only available for certain submission systems

**[Comment 18]**

Rejected images would need to be checked by a human. Encourage researchers to check their images as soon as generated and not to wait until submission.

**[Comment 19]**

This approach could end up penalising work that was performed before said antibodies were withdrawn from the market. The users at the time may have not been aware of these performance issues. Automation is always a very attractive solution on paper but it requires embedding some nuances from conception.

**[Comment 20]**

These tools exist and can be quite powerful, but I don't know how accessible they are

**[Comment 21]**

While this could be an effective way to minimize the impact of bad antibodies being cited, the feasibility of publishers having access to this information seems low. It also brings up questions such as "do all papers that have used these antibodies need to be retracted?"

**Item R8**

*Validation Strategy Reporting Using IWGAV. Authors should describe how antibody specificity was assessed, making reference to the International Working Group on Antibody Validation (IWGAV) framework.*

**Participant Comments**

**[Comment 1]**

In my experience a fundamental limitation on improving antibody validation and reporting it is the fact that many authors simply don't understand the issues or importance of doing so. Making such researchers fill out such forms I fear will only lead to incorrect information being generated, and possibly adding to the confusion.

**[Comment 2]**

checklist is pretty easy to put into the journal instructions to authors, but someone has to follow up with journals and author compliance is the main issue

**[Comment 3]**

This would help confidence in antibody validation.

**[Comment 4]**

Barriers may including understanding what the minimum standard is, although this is a good idea to help with awareness of the five pillars

**[Comment 5]**

I have concerns about how feasible this is as the validation data would need to be peer reviewed, and as noted in Q8, I have concerns that this is one too many demands on peer reviewers. However, if authors can opt in to submitting a validation check list, which can go to a technical reviewer eg early career researcher in the lab of one of the main reviewers, and this can result in a badge in the paper recognising that it meets high standards, then it might be more feasible (see also comments on Q5).

**[Comment 6]**

I marked this as a 6 for feasibility because adequately implementing this policy on reporting Ab validating strategy, seems quite difficult. Journals can have guidelines on this that refer to the IWGAV pillars, but adequate implementation of this by an automated tool is more challenging and for now should lie with the reviewers. Reviewer training per se is difficult hence the 6.

**[Comment 7]**

Researchers should be doing this for every experiment. I think the hurdles will be getting them to share the data and getting the journals to throughly review the information.

**[Comment 8]**

A checklist is easy to implement and a good reminder. However, doing this at the time of publication submission is probably too late in the game. Research groups are not likely to go back and try to validate the antibody after the experiment is completed.

**[Comment 9]**

Is there a large enough of catalog of abs available that are validated using IWGAV? Is all of the data available showing IWGAV developed using appropriate KO cell lines that are correct?

**[Comment 10]**

Again, this data will be most effective available in manufacturer portals (product websites); authors could refer to those portals in the supplemental information of a journal if needed.

**[Comment 11]**

Antibodies must be validated in the context of a given application. Broad, sweeping assumptions of specificity based on the use of a given pillar in a given application are potentially misleading.

**[Comment 12]**

Is everyone in agreement on the utility and validity of the International Working Group on Antibody Validation (IWGAV) framework?

**[Comment 13]**

It is unlikely that authors will be aware of the framework or have performed these experiments. Further education and training would be required across the research community before this would be effective.

**[Comment 14]**

This is an ideal situation, but without community agreement up front, leaving it to publication is unlikely to result in uptake. More effort needs to be placed on showing authors what the benefits of doing this are for advancing science so it does not "feel" like another pointless task associated with the hurdle of getting published. It also needs to be consistent across publishers and if it is not included in the text of the article will be difficult to be truly beneficial in the long run - and then you run into the paper length restrictions of some publishers.

**[Comment 15]**

I've opted out as I don't have specific expertise on, or experience of, the framework as a whole although I strongly support authors reporting validation along recognized standards within a discipline. However, adding another checklist to any manuscript submission process becomes problematic as evidence suggest that authors often don't know how to fill these out and will also fill them out incorrectly if they think their chances of being peer reviewed are higher if they do. Checklists during the submission process are in general increasing across all disciplines (e.g. around open science - some have likened this to 'death by a thousand 10 minute tasks'). For some multidisciplinary journals checklists can also slow down the submission process and act as a barrier to submission. This may reduce the incentive for publishers to encourage them or to enforce them (e.g. the lesson from ARRIVE). Checklists, once filled in, have to be assessed either manually or through the development of automated screening. I'm not saying that reporting such standards should be discouraged but without enforcement and monitoring evidence suggests checklists do not help.

**[Comment 16]**

Most independent research labs are not equipped to provide this level of robust validation. It is an important aspirational goal but perhaps needs a consortium approach.

**[Comment 17]**

many papers use dozens of antibodies making this less feasible. Also many authors would likely do this right before submission and maybe some mAbs wouldn't have been assessed this way?

**[Comment 18]**

Requiring the inclusion of validation methods with all papers is a good first start for improving reproducibility. It won't completely solve the issue but it will at least ensure that researchers are validating the antibodies they are using before they begin their experiments.

**Item R9**

*Genetic Validation Evidence for Critical Applications. Where antibodies are critical to key experimental conclusions, authors should either present validation data based on a genetic strategy or provide a scientific justification for why a genetic strategy is not feasible or necessary.*

**Participant Comments**

**[Comment 1]**

The use of KO/KD genetic strategy should be required but allowance for circumstances where it may not be feasible should be in place also.

**[Comment 2]**

would be great to do but not sure how this can be enforced

**[Comment 3]**

One issue could be the availability of methods for validating data for essential genes - it may be in this case that you need to accept an alternative validation method.

**[Comment 4]**

I have concerns that mandating that a genetic strategy is required may not be feasible in certain model systems or for certain epitopes. However, when antibodies are critical to key experimental conclusions, authors should be providing validation, and it is reasonable to expect referees to check this.

**[Comment 5]**

Same comments as Q8, but marked higher for effectiveness, as here in Q9, for studies that rely heavily on the use of Ab, using genetic approaches seems critical for demonstrating Ab performance and thus validity of the conclusions

**[Comment 6]**

This is fine if you have KO data for a specific application (i.e., WB, ICC/IF, IHC, etc.), but there are plenty of high-performing antibodies that either cannot be validated easily through genetic strategies (e.g., PTM antibodies, pathogen antibodies, essential protein antibodies) or no KO/KD data exists.

**[Comment 7]**

It is very difficult and expensive to get knockout tissues/samples and for some genes (lethal KD/KO) impossible.

**[Comment 8]**

Often doable in these days of CRISPR-Cas. But maybe a step too far for many labs. Certainly an excellent idea.

But I don't think that previously published data helps in these contexts, because we are always considering the antibody in the here and now. In the users hands as they have it. Not some other (RRID?) antibody in someone else's hands.

**[Comment 9]**

Genetic strategies (especially knockout) should be the gold standard. Presenting data would be the best outcome, but citing previous validation data could be sufficient although would not account for method/lot differences that might affect specifcity.

**[Comment 10]**

If the genetic controls are not available, this sort of validation is extremely difficult

**[Comment 11]**

See question 8

**[Comment 12]**

Knockout or Knockdown data are important but no one piece of data (genetic, orthogonal, independent antibody, tagged protein, IP-MS) trumps the others; the more validation data you have, the more confident you can be in the specificity of the target antibody. Some protein targets are lethal if knocked out.

**[Comment 13]**

KO data are, of course, compelling, but are only one data point in what should be a more comprehensive validation strategy at the level of the application.

**[Comment 14]**

I imagine these studies will cost money to do and need to be built into grant applications - will funders see this as good value for money?

**[Comment 15]**

As above, authors may not have performed these experiments (although offering the option to reference pre-existing validation data derisks this). Journals will need to insist on this information being provided for authors to undertake the experiments.

**[Comment 16]**

this should be straightforward for authors to report where there are the data available and could be included in e.g. author guidelines but it again comes down to who is accountable for the checking and enforcement.

**[Comment 17]**

Funds need to be made available to researchers to support this work

**[Comment 18]**

Importantly, the value of gene-editing based antibody validation strategies is dependent also on the orthogonal validation of the gene-edited system!!! It is not trivial.

**[Comment 19]**

unclear who determines whether the antibody is critical -- a reviewer could argue they are all critical and ask for the validation to be done.

**[Comment 20]**

While this is undoubtedly the best way to demonstrate antibody specificity it is also time consuming and expensive

**Item R10**

*Antibody Validation Plans in Grant Applications. Funders should include a section in the application form where applicants must detail the steps they will take to validate the antibodies used.*

**Participant Comments**

**[Comment 1]**

After having invested my time in talking with NIH Institute leadership to fund more antibody validation research, I now understand that is not going to happen. But every funding agency should be encouraged to require more than simply a section on Reproducibility but specifically require validation data for any antibodies that will be used and critical to the proposed studies.

**[Comment 2]**

NIH has this requirement: key biological resources supplemental file https://grants.nih.gov/grants/guide/notice-files/NOT-OD-17-068.html

example compliance document https://library.ucsd.edu/research-and-collections/research-data/_files/ExampleAuthenticationKeyBiologicalChemicalResources201609b.pdf found at https://library.ucsd.edu/research-and-collections/research-data/plan-and-manage/nih-policy-on-rigor-and-reproducibility.html

**[Comment 3]**

I feel as though this could become another box ticking exercise for the applicant, which is filled with little care and using generic text each time

**[Comment 4]**

I marked 6 for feasibility as I am not sure how easy it would be for funders and grant committees to evaluate this extra information.

**[Comment 5]**

The main issue here is follow through. In many cases, once the grant is funded there is no over-site to ensure follow through. If more grants required meeting milestones for the subsequent years it would be more effective.

**[Comment 6]**

Detailing the steps is different from taking them.

**[Comment 7]**

This could be implemented in application templates across funding organizations. But it will likely become a copy-paste scenario where applicants pay lip service to the principles of antibody validation but in practice skip this step to save on time/budget. The benefit of this would be solely in serving as a reminder and soft encouragement.

**[Comment 8]**

Would you know this information prior to starting the study? Will researchers always validate their own abs or utilize other sources (suppliers websites, previous publications)?

**[Comment 9]**

It's always up to the author to validate the antibody in the context of their experiments; make sure the protein target is present in the cell or tissue model they are studying. Their job is not to show that the antibody works in general (they can refer to the manufacturer's product pages for that information).

**[Comment 10]**

Easy for funders to implement. Effectiveness depends on how essential the information is (i.e. will the project be scored based on this information, or is it an optional extra?).

**[Comment 11]**

I feel this is feasible for funders to add, but again it could get lost in the overall list of requirements and if not policed will be hard to measure

**[Comment 12]**

Put simply, you get what you pay for

**Item R11**

*Dedicated Budget for Antibody Validation. Applicants to biomedical funding schemes should include a specific line item within their budget for resources to conduct robust antibody validation where relevant to their proposal.*

**Participant Comments**

**[Comment 1]**

Unclear whether these funds would actually be aligned to this

**[Comment 2]**

I am not certain whether this extra budget is available to most funders, hence marking 6 for feasibility.

**[Comment 3]**

This is a great idea. Antibody validation is expensive but setting aside a budget ahead of time would help researchers to plan ahead. That said, research is expensive and the funding situation (at least in the US) is precarious right now.

**[Comment 4]**

That is a good idea. And the importance and utility of having it rises very sharply depending on how critical the antibody is to the research project in question. One recalls sadly ERbeta. The "but" is that it may be an expensive add on.

**[Comment 5]**

May be a challenge to include costs where there is a budget limit, necessitating a trade off between validation and key experiments. Some funders may not appreciate the strategic importance of validation and therefore be less willing to shoulder the cost. It may also slow the pace of research which could be detrimental to some organisation that rely on public donations to support their funding activities.

**[Comment 6]**

While good in practice, research proposals typically employ multiple antibodies in a study, sometimes upwards of 5-10. Would the applicant be required to validate all antibodies prior to use? Would they need to validate them in their specific experimental cell/tissue system? This would result in a pretty significant budget increase which would come at the cost of reducing the scope of the hypothesis-driven research study or cutting down the number of applications a funder can fund for a program.

**[Comment 7]**

This assumes researchers will have to validate every antibody without using other sources

**[Comment 8]**

Again, It is always up to the author to validate the antibody in the context of their experiments.

**[Comment 9]**

If specific funding is provided authors will complete the requires experiments. Easy for funders to implement (in terms of requesting the information) but in the current funding climate grants are already tight. Would this take away from the budget for answering the research question?

**[Comment 10]**

Funding the extra QC and effort will help effectiveness

**[Comment 11]**

re the feasibility question: I don't know how much this costs but my assumption is that this should be feasible and is worth the cost to funders given that it will likely increase their ROI

**[Comment 12]**

While this is a good idea I fear that funding levels are tight and getting money assigned to validate reagents will be difficult.

**Item R12**

*Funding Development of Critical Antibody Tools. Funders should create or expand targeted schemes to support the development and validation of critical antibody-based research tools.*

**Participant Comments**

**[Comment 1]**

There is no reason why these steps should not be implemented asap. In a related issue, there is currently no repository for cell lines KO'd for specific targets and such a resource would be a valuable resource for several uses.

**[Comment 2]**

Difficult to know if this is feasible - which depends on the funders have sufficient money to implement these schemes. It's possible that the money might be better spent on education?

**[Comment 3]**

Similar rationale to scoring as for Q12

**[Comment 4]**

There needs to be a larger initiative by funders to not only create validated tools but also to promote them. Currently researchers want to use cited antibodies even if they are poor. They are resistant to change even if a new well validated tool comes out. In addition to providing well validated antibodies there needs to be a strong initiative to educate researchers about the new reagents and push them to use them.

**[Comment 5]**

It would depend on a cost-benefit analysis. Otherwise funders might be very reluctant " what for? There are loads of antibodies (and they work)" type of resistance. So, a picture of the actual current costs of bad antibody validation in terms of wasted funds (and research years, peoples careers' etc. etc. as documented in the various validation commentaries in the literature) would be helpful to stimulate interest. I think we are still pretty dependent on Len Freedman's ball park estimate from a decade or more ago.

**[Comment 6]**

This may be a challenge in the orphan disease arena, where funding is limited. However, it can offer an opportunity for funders to collaborate (for example, as had occurred with the amyotrophic lateral sclerosis community in the past) in supporting antibody production and validation,

**[Comment 7]**

people may not want to spend time and resources on antibody validation if it's not required.

**[Comment 8]**

i am biased towards this suggestion so have given it a high score. but in practice, this would again reduce the budget of a smaller funding organization and could be hard to justify/implement as it is not something that is easy to explain or attractive to patients/donors for fundraising efforts.

**[Comment 9]**

Validated recombinant antibodies where sequences are publicly available would be the best solutions

**[Comment 10]**

This would potentially help some of the targets, but there are 22000 proteins.

**[Comment 11]**

Much of this could depend on the nature of the target. If there are no consistently reliable cell models that would make studying the protein of interest difficult. KO models could be lethal depending on the target and KD may not show a big difference if endogenous levels of a given protein are very high. Technology is moving towards animal-free antibodies but it can be difficult depending on the nature of the target.

**[Comment 12]**

Requires buy-in from funders to use budget specifically for this purpose - may be mixed.

**[Comment 13]**

I would include funding the development of research critical antibodies.

**[Comment 14]**

If by "animal derived" antibodies, you mean moving from polyclonals and hybridoma-derived toward recombinant, I would generally agree. Although I think that there will always be a role for polyclonals in early research and for certain applications. It is hard to make the case that research cannot be conducted until you are assured of a rigorously validated, recombinant monoclonal.

**[Comment 15]**

The vast number of antibodies available would make it challenging to create the resources described above.

**Item R13**

*Signalling Value Without Mandating. Funders should share information within the applicant guidance that antibody performance is an important limitation in many experimental methods. Applicants should be encouraged to address this in their methodological sections.*

**Participant Comments**

**[Comment 1]**

If I understand this correctly, it would involve simply stating in the instructions for applications that antibody validation strategies and data should be included to support data interpretation and reproducibility. Yes, this is an easy and useful strategy.

**[Comment 2]**

Doubtful that this would be strong enough to drive best practice, and will depend on the implementation by the reviewers, who themselves probably require educating.

**[Comment 3]**

Similar rationale as Q10, unclear how much resources funders have to enable this extra check/evaluation, maybe some of this type of assessment could be done in conjunction with AI tools

**[Comment 4]**

I like the idea of getting researchers to think about this in the application process at early stage of the work.

**[Comment 5]**

Yes. A nudge, and thinly veiled threat. But certainly a wake-up for people who STILL don't understand the issues.

**[Comment 6]**

For reasons stated previously, feasibility may vary from funder to funder.

**[Comment 7]**

Similar to Q10, I think this will just end up in a copy/paste situation where applicants are paying lip service to the principle without affecting behavior.

**[Comment 8]**

Disagree fundamentally with this

**[Comment 9]**

If it's not a mandate then incentive for applicants to do this is not there and therefore will not be effective

**[Comment 10]**

Effectiveness depends on how strongly funders insist on the inclusion of this information and Panel training in the assessment of the information.

**[Comment 11]**

This helps push the needle and raise awareness.

**[Comment 12]**

I'm not sure that sharing information alone is that effective without some consequences/reward for those authors that do not or do take heed of this

**[Comment 13]**

"Signalling Value Without Mandating" seems not much different than the current situation.

**Item R14**

*Encouraging Open Sharing of Validation Data. Funders should encourage grantees to deposit antibody validation data in open-access repositories, ideally linked to RRIDs or registry entries.*

**Participant Comments**

**[Comment 1]**

While such a system as that proposed here might help a bit, a far better strategy in my opinion is to require that all journals require such data before publishing an article that uses antibodies.

**[Comment 2]**

We will need a place to share this data that is relatively easy to use and well known

**[Comment 3]**

The issue here is the quality of the "antibody validation data".

**[Comment 4]**

This should be feasible but I have a similar concern about who will be checking the validation data to ensure robustness. I worry it will be a massive, meaningless data drop just to fulfill the requirement. It should be a reviewed/curated process but who would do this work?

**[Comment 5]**

I think you can leave the "ideally" out of the question. Unequivocal identification using for example RRID is easy and a simple step to getting antibodies validated.

**[Comment 6]**

It can be a challenge to robustly enforce end of grant reporting. Follow up requires staff resource which may not be available. Charitable funders in particular tend to focus resource on new studies (and funding commitments) out of necessity, in order to generate funds form the public.

**[Comment 7]**

The most you could do would be to "encourage". I don't think it's feasible to mandate or track compliance.

**[Comment 8]**

see comment above about RRIDs

**[Comment 9]**

Experimental parameters are important. And it's important to note if the authors' validation data is in line with the manufacturer's recommendations (dilutions, approved applications, etc.).

**[Comment 10]**

Highly feasible, but only if authors already have the validation data.

**[Comment 11]**

Feasibiilty comes down to whether there is also education/training for authors about where and how to deposit the data and also the willingness of funders to monitor and enforce deposit. Note it would be easier for publishers to enforce at publication if there is an expectation that funders will also be monitoring and enforcing this at an earlier stage in the research lifecycle

**Item R15**

*Endorsing Journal or Community Guidelines. Funders should formally endorse community-developed reporting standards that promote antibody transparency and validation (e.g., IWGAV, MDAR).*

**Participant Comments**

**[Comment 1]**

It is difficult to get funders to do things like this. We have tried, but they are frequently locked into various difficult to change rules.

**[Comment 2]**

This seems possible, but perhaps not strong enough to drive real change in researchers who have a range of other priorities. It might get overlooked

**[Comment 3]**

I am marking this as a 9 on effectiveness as I feel that this would provide the type of incentive needed to drive a change in research practice/reporting. I expect 'endorsement' to be relatively feasible by funders. Endorsement could be in the form of a request to applicants to demonstrate past adherence to community-developed reporting standards as a potential pre-requisite for consideration.

**[Comment 4]**

Feasible and a low lift. But will most likely be ignored.

**[Comment 5]**

Needs to be consistent across funders wrt which reporting standards are endorsed, unless all are held in equally in high regard

**[Comment 6]**

If funders and publishers collborate on the standards and funders then mandate those standards, publishers will generally be much more willing to ensure there is compliance. Endorsement alone will help but funders also need to require that standards are met to be really effective

**[Comment 7]**

Again, you get what you pay for. Funders should value high quality, well-validated, tools/reagents to promote robust research. "Give me six hours to chop down a tree and I will spend the first four sharpening the axe"

**Item R16**

*Encouraging Manufacturer Transparency. Funders should directly engage with manufacturers to encourage them to publish antibody validation datasets and adopt consistent metadata standards (e.g., RRID, clone ID, lot number).*

**Participant Comments**

**[Comment 1]**

If all commercial sources made available all validation data, with clear descriptions of how the data were generated, it would certainly enhance the research and help researchers choose optimal reagents to purchase. However, it is not commercially feasible for companies to be made to perform such validation, given the return they get on most antibodies. Perhaps some system for publicly awarding good actors notice of their efforts would help steer potential buyers to more reliable sources.

**[Comment 2]**

Good producers and vendors already do a lot of this. Can encourage them to do more

**[Comment 3]**

Difficultly to get manufacturers onboard unless there is a financial incentive for them to do this.

**[Comment 4]**

Suppliers should be publishing more detailed data, including negative data and RRID should be essential - although this is hard to implement. But this could tie in with the journals because if the journals require it then it will in a sense become mandatory

**[Comment 5]**

Most top manufacturers already do this.

**[Comment 6]**

I think the funding situation (at least in the US) makes this difficult. I just don't see federal funding agencies pursuingf this.

**[Comment 7]**

Of course there needs to be a carrot here. So, maybe funders would maintain and provide researchers with a list of manufacturers who do this? Why buy a reagent from a supplier who doesn't meet such standards? This might encourage a rush to get on board from reluctant suppliers. And of course, for the funders, they would know that less money was being wasted on useless (or unvalidated) reagents.

**[Comment 8]**

This is not in the scope of most funders work. Plus, finding appropriate points of contact for the manufacturers would be very time consuming and difficult. And unless the funder is funding that manufacturer and has built a relationship with them, why would the manufacturer modify their behavior?

**[Comment 9]**

In an ideal world, manufacturers would only sell recombinant antibodies. While publication of sequences would be ideal, this is unlikely to be feasible due to commercial constraints. However, an escrow service of some kind (which could be played by RRID) which curated and stored antibody sequences, and provided accession numbers to manufacturers so scientists would know which antibodies are identical

**[Comment 10]**

Smaller companies may not have the funds to register with RRID, etc

**[Comment 11]**

Many manufacturers do this already and more validation data are being added as technology improves.

**[Comment 12]**

All of these things are indeed of value, but I an concerned about the long list of requests put to funders and where this would fall on their prioritisation list

**[Comment 13]**

Antibody manufacturers can (and should) share their validation data this still doesn't account for variability within experimental samples.

**Item R17**

*Promoting Participation in Independent Benchmarking. Funders should co-fund independent benchmarking initiatives such as YCharOS or equivalent open-science consortia.*

**Participant Comments**

**[Comment 1]**

I am not aware that the NIH has any initiatives like this, and I think the present funding climate is not conducive to the initiation, or expansion, of any specific efforts.

**[Comment 2]**

I think they also need to promote these initiatives more. Their work is fantastic but I don't know how many users are aware of these initiatives and using them to direct their antibody decisions.

**[Comment 3]**

Very good idea. Money limited, but YCharOS is going in the right direction

**[Comment 4]**

It will be effective in some fields, but unlikely to occur in others.

**[Comment 5]**

The YCharOS initiative is great, but the resources required for producing knockouts for every target are high and the rate of validating antibodies is slow.

**[Comment 6]**

This would be effective and is the approach we take, but as with Q12 this would not be feasible for smaller funding organizations and is very hard to fundraise for.

**[Comment 7]**

Even if antibodies are not performing adequately as shown by an independent benchmarking initiative, manufacturers do not have to remove these products from the market.

**[Comment 8]**

YCharOS does good work, but vendors must pay to participate. This is perhaps not a sustainable model.

**[Comment 9]**

Where will the funding come from for this? Although it is important, it's not "sexy"

**[Comment 10]**

I have questioned the feasibility of this because it is incredibly difficult to persuade funders to fund any infrastructure let alone to sustain that funding.

**[Comment 11]**

It would be important to ensure that this doesn't only occur for key targets.

**[Comment 12]**

This type of consortium approach seems like the only way to make a real change.

**[Comment 13]**

While these initiatives are effective it will take some time before enough antibodies are characterized to make an impact.

**Item R18**

*Promoting Participation in Independent Benchmarking. Funders should promote industry participation in independent benchmarking initiatives by asking applicants to consider working with initiatives such as YCharOS.*

**Participant Comments**

**[Comment 1]**

There should be a sales person that advises on this. Strict rules generally govern what funders are able to say and do, but even a few words from a funding associated person can be a good marketing strategy towards industry. Private funders can say much more, but command less respect.

**[Comment 2]**

It would be great to have industry involved in the initiatives, but industry scientists are often constrained by the organisation in terms of resources to support these initiatives. The funding bodies have little influence over industry participation

**[Comment 3]**

I don't know how actively industry relies on external funders. If they are not relying on the external funding then it would be hard to push them to become involved.

**[Comment 4]**

It would seem to depend on the importance of the antibody in question to the project in question. Only a big research group might have the necessary facilities to do this. And we want all labs to validate antibodies. But if a working structure for the collaboration could be found (i.e. who prioritizes which project YCharOS does first if it has a choice?) then good thing.

**[Comment 5]**

Not convinced of feasibility, as industry tends to plough their own furrow.…

**[Comment 6]**

This would only be feasible if the applicant could have YCharOS participate on the grant as a co-investigator to receive funds for the work. I don't think YCharOS has a large pot of money to validate antibodies pro bono. Plus, the timeline associated with YCharos adding a target to their pipeline and performing the validation might dramatically affect the project timelines (or worse yet, find no suitable antibodies and thereby result in the need to terminate the grant).

**[Comment 7]**

Funders have limited influence on industry partners.

**[Comment 8]**

I'm just not sure about the effectiveness of promotion alone - it could help if there were many funders involved but calls from e.g. one or two is less likely to be effective

**Item R19**

*Tailored Education and Training on Antibody Validation. Training in antibody validation should be offered in relevant bioscience courses and programmes.*

**Participant Comments**

**[Comment 1]**

It would be difficult to get all institutions on board. Different institutions, countries, will have different priorities and policies.

**[Comment 2]**

Difficult to know how to influence this across a broad range of institutions?

**[Comment 3]**

I set something like this up at University of Turku. It is so important that the young bioscientists see what a chaos bad antibodies cause, and much easier to teach them than the oldies. Start at the roots.

**[Comment 4]**

Need to train the senior staff as well!

**[Comment 5]**

I believe that in many cases the emphasis on validation and proper controls comes down to lab culture. If a student is trained to validate antibodies but the PI/senior lab members do not agree this is a priority, it will not be done. Also, implementing a board training across institutions would be difficult.

**[Comment 6]**

Antibody validation should be recognized as a research discipline and incorporated into cell biology curricula.

**[Comment 7]**

How will training be funded? Who will deliver?

**[Comment 8]**

I think education and training will be key to ultimately improving antibody validation - so that it just becomes the norm. Requirements and enforcement by funders or publishers will only be effective if researchers understand why it is so important and are trained appropriately (it's just too late by the time a study gets to publication). I have said that this should be very feasible in principle to implement but bear in mind that I'm not really sure how easily such training can be introduced systematically into institutions that do not have this and/or are unaware of the importance.

**[Comment 9]**

Not necessary for undergraduates but should be part of PhD programs. Practical training would be challenging and may have cost barriers but a lecture on best practices for using antibodies and other reagents should be feasible.

**[Comment 10]**

Better training is required, the questions that need to be addressed are: who funds this, who is responsible for the training, who sets the standards, etc

**Item R20**

*Research Integrity and Ethics Frameworks. Universities and research institutions should incorporate antibody validation expectations into research integrity and ethics review frameworks.*

**Participant Comments**

**[Comment 1]**

While such strategies should absolutely be in place today, the main barrier to doing so is ignorance of the issues by the leadership at all levels of academia.

**[Comment 2]**

This could work as long as the guidance is clear enough, and ideally uniform across institutions

**[Comment 3]**

A lot of research does not go through ethics committees.

**[Comment 4]**

I don't really see this as an ethical issue. Lumping it into ethics with big issues like image manipulation, data fabrication, animal use, etc doesn't seem appropriate to me, personally. Also, if a lab publishes without antibody validation data, could ethics complaints be lodged against them?

**[Comment 5]**

Again, there is probably a long list of things that should be done, and getting something to the top of the list to implement is challenging. Demonstration of what is lost to future scientists by omitting this data would provide value

**[Comment 6]**

Incorporating training into existing ethics and integrity frameworks is a hugely important signal to researchers that this is not just good research practice but there is a moral imperative to ensure it is done correctly because of the immense damage done without appropriate validation. Such frameworks could include the consequences (economic, health, equity etc). Such training will hopefully help make researchers think twice before skipping a step or help incentivise researchers to do the extra steps required for validation.

**Item R21**

*Support for Local Champions of Best Practice. Universities, research institutions and learned societies should recognise and support local champions or experts in antibody validation.*

**Participant Comments**

**[Comment 1]**

While I believe this is a much needed item, I don't believe this issue has much likelihood of rising to a sufficient level of awareness that is will be supported by an institute at this time.

**[Comment 2]**

We tried this at our university, maybe it was not the right time, but we could not get a reproducibiliTea group to launch properly

**[Comment 3]**

Being rolled into outreach training would make good sense

**[Comment 4]**

Of course, it won't work if the users don't use the facility...

**[Comment 5]**

I like this idea. It has the flexibility/opportunity to educate multiple levels of the system (students, postdocs, PIs) and provide a direct connection to someone for further discussion, insight, and support.

**[Comment 6]**

High turnover of research staff - how to maintain long term champions?

**[Comment 7]**

all of the questions in this section are not going to make antibody research more reproducible per se but will lay the foundation for increased awareness and adoption of best practice for researchers and disciplinary communities. Without laying such a groundwork of training and advocacy (as this specific question promotes), antibody validation will never become the norm.

**[Comment 8]**

Would these roles be a full-time job or would this be someone doing it in addition to their research? It would require a lot of time but if it is a full time job who would fund it?

**Item R22**

*Establishing a Coordinated Roadmap. Stakeholders should work together to develop a shared roadmap for improving antibody validation practices by 2030.*

**Participant Comments**

**[Comment 1]**

Organizing meetings of stakeholders at funding agencies (e.g., NIH), national/society meetings, and others should be encouraged and enacted asap

**[Comment 2]**

It may be difficult to reach consensus, as shown by prior work within the IGWAV group, which may result from different motivations and drivers for different individuals

**[Comment 3]**

I could see these types of cross-stakeholder efforts getting bogged down. Everybody thinks they know best.

**[Comment 4]**

Scored low on feasibility as there will be challenges achieving cross sector collaboration and leadership.

**[Comment 5]**

Engages various groups for informed recommendations across sectors.

**[Comment 6]**

This is a conversation that's been going on for about 30 years. I published a widely cited (my most cited) commentary in 2015 making the case for recombinant antibodies with published sequences. Since then there has been some improvement in antibody providers striving to sell recombinant antibodies, rather than polyclonals, but progress has been so so so slow. Everyone points fingers at everyone else to indicate where the responsibility should lie

**[Comment 7]**

How is this different from the IWGAV?

**[Comment 8]**

Time and funding are always major factors.

**[Comment 9]**

This effort, broadly, is too fragmented. Alignment feels difficult to achieve across the various initiatives.

**[Comment 10]**

Funding is required to get the right people together to create a roadmap and continued funding is required to get it implemented. These things traditionally take years or decades to achieve across all the various stakeholders but you have to start somewhere and gather momentum

**[Comment 11]**

Rating 7 only because 2030 is not so far away and roadmap development involving multiple stakeholders can be a lenghty process.

**[Comment 12]**

It is a great aspiration, just unsure that it could work

**[Comment 13]**

There have been at least 2 initiatives to implement a framework for antibody validation across stakeholder groups that have started but were not able to develop a final product. The reasons for failing were different (one was funding related and the other was inability to reach a consensus) but they are illustrative of how difficult a problem this is to tackle.

**Item R23**

*Shared Infrastructure for Validation Data and Standards. A coordinated, shared infrastructure should be developed (or expanded) to aggregate antibody validation data, provide standardised formats and tools for sharing, and enable journals, funders, and institutions to access validation status or red flags.*

**Participant Comments**

**[Comment 1]**

A system such as linking NCBI protein data to antibody validation data for antibodies targeting each protein would make it much easier for researchers to find and obtain optimal antibodies for their specific uses/needs

**[Comment 2]**

Good idea but unclear if the commercial enterprises would want to be involved. Could work for the not-for-profit organisations however

**[Comment 3]**

Again, I see this bogging down. This would be a massive undertaking.

**[Comment 4]**

Funding might be an issue. And how would commercial for profit entities view the idea?

**[Comment 5]**

As mentioned, there are many websites out there that don't speak to eachother (eg RRID, Benchsci, YCharOS, CiteAb, etc). Building bridges across these sites would not only improve information offered, but also hopefully increases awareness of these resources by amplifying messages and consolidating communications.

**[Comment 6]**

to make this valuable the issues raised in the RRID question need to be resolved. Again, best approach would be to only use recombinant antibodies, and have sequences held somewhere in escrow.

**[Comment 7]**

Who would be qualified enough to decide which is good or bad, as well as not have a stake in the decision?

**[Comment 8]**

Suggest utilising existing infrastructure as far as possible

**[Comment 9]**

This would be a fantastic resource but only if it had sustained funding and extremely good, and community-based ,governance. There are three resources I would like to point to that might be useful in thinking about any development of a coordinated and community based infrastructure or organisation:

1) The FOREST framework: a values based framework developed for organisations involved in scholarly communication- https://www.nextgenlibpub.org/forest-framework

2) The POSI principles, a set of guidelines by which open scholarly infrastructure organisations and initiatives that support the research community can be run and sustained - https://openscholarlyinfrastructure.org/

3) Invest in Open Infrastructure , an organisation that specifically addresses the funding requirements of, and investments for, open scholarly infrastructure

I know some of the organisers at all three initiatives if you'd like me to put you in contact with them

**[Comment 10]**

Resource needs to be available to researchers.

**[Comment 11]**

Again, definitely worthy goals - the issues outlined above apply here too.

**[Comment 12]**

I think sharing high quality data that can be used by all platforms, in the way it suits their platform and users, will have more impact than trying to develop a single platform.
