## Supplementary material for "A multi-stakeholder Delphi consensus study proposes actionable solutions to address antibody validation failures": S8 Text

**S8 Text. Round 2 qualitative feedback (participant comments).**

*Free-text comments provided by Delphi panellists during Round 2 (n = 30 participants, 22 items: 12 re-rated from Round 1 and 10 new items). Comments are presented verbatim and organised by item. Comments are numbered sequentially within each item; numbering restarts for each item and does not track individual participants across items.*

**Item R3**

*Authors should report the dilution ratio at which each antibody is used. In addition, authors should state the antibody protein concentration where possible. Examples include dilution ratios such as 1:1000 or 1:500, or concentrations expressed in µg/mL.*

**Participant Comments**

**[Comment 1]**

I would propose a re-wording to prioritize protein concentration (when available) over dilution and to include a statement to the effect of total dilution from commercially obtained source. In my lab we routinely dilute 1:1 with glycerol for storage at -20C and this 2-fold dilution is rarely included in calculations/reporting by lab staff. Also, if no protein concentration is available and not commercially obtained then concerns rise.

**[Comment 2]**

Would require all journals to implement - unclear how challenging this would be? Nevertheless - seems very feasible to expect them to do so. But would require a coordinated effort between main / top journals. Otherwise this could be a disincentive for publishing in one journal versus another

**[Comment 3]**

Clarification: I previously marked feasibility as 6 because I was considering the feasibility of a meaningful policy at journals, where compliance with the policy is checked (the barrier to this being that resources at most journals are limited). If, however, this question refers to the feasibility of journals having guidelines on their websites such as 'Authors should report the dilution ratio at which each antibody is used. In addition.…" then the scoring would be higher. So, I have moved it to 8. My experience with journal policies is there is that there is little to no author compliance unless policies are mandated. I feel that the biggest barrier to improving reproducibility (assuming that the policy is not mandated by funders, which would be the best incentive for behavioural change) is for journals to have the resources (or reliable technology) to check for compliance on reporting, as an incentive to publish in the journal. The tools that I tested so far do not reliably identify eligible manuscripts, eg. studies where this policy would be applicable, is the item reported as expected as per the policy, is it coherent across the paper etc. I expect tools to improve and this to be automated in the future so I am hopeful that with technology reproducibility will improve.

**[Comment 4]**

I don't accept that a provider should describe an antibody concentration in the material they sell as 'not always available'. I understand that while we still have polyclonals on the market the concentration of IgX in the product may be an unknown, and due to storage and manufacturing issues the amount of "functional IgX" in the product may anyway be variable, but it really is not hard for a serious provider to tell the customer the concentration of antibody in the stuff he is selling. This at least lets everyone compare the data sets using a particular antibody. If the provider can't do that it implies that they really just put the material in a box and send it out.

**[Comment 5]**

Examples include dilution ratios such as 1:1000 or 1:500, and concentrations expressed in μg/mL (suggest AND instead of OR)

**[Comment 6]**

I think it would also be helpful to note "used manufacturer's recommended dilutuion."

**[Comment 7]**

Protein concentration should be reported as much as possible as dilution ratios alone may not be sufficient information for others to reproduce an experiment accurately.

**[Comment 8]**

For monoclonal antibodies the concentration in ug/ ml should be used —- different sources will have different concentrations and dilutions could be confusing. Dilutions for polyclonals is ok but those aren’t renewable

**Item R4**

*Deposition of Validation Data in Open Repositories. Journals should require authors to deposit antibody validation data in an open-access repository, linked to a persistent identifier (e.g., RRID, DOI).*

**Participant Comments**

**[Comment 1]**

While if implemented I think it would be great and effective, I don't see this as likely to be implemented as it requires considerable additional work to put together an additional manuscript. Also, I think the data belongs in the manuscript (likely as supplementary data) where the antibody is used. If reporting on the use of a newly made antibody and used for the first time it should be required in that manuscript. Journals should also require identification of any new antibodies and the validation data.

**[Comment 2]**

Requirements for validation need to be clear. Zenodo submitted information will not enforce any requirements. Antibody registry will only enforce validity of the information not the reagent. If there is appetite for this we need a specific repository.

**[Comment 3]**

There needs to be clearer guidance, expectations for the validation data, online repositories where data is stored.

**[Comment 4]**

Perhaps this best sits with the journal, and should be included in the peer review process. I agree with the comments about the review being necessary to ensure quality.

**[Comment 5]**

For this to be useful, the quality of the data would need to be reviewed. This should ideally be done as part of the peer review process of the associated manuscript. Also, this wound need clear guidelines on what is needed, so that journals can check that the required validations have been provided. This would also require an additional series of checks for editors, which unless it can be somehow automated, would be an additional potentially time consuming task.

**[Comment 6]**

Feasibility remains low to me because as noted in Q3 making this a requirement for journals is currently challenging given limited resources/technology.

**[Comment 7]**

I agree with the criticisms. The precise nature of the validation data and its format would need to be standardized, otherwise we are likely making a new form of chaos. And who defines the standard? OGA / YChaROS and EuroMabNet might like to chat with each other on this point.

**[Comment 8]**

I rated it high for the sake of transparency. I think that a standardized system for depositing and sharing validation data would be better, but this is a heavy lift.

**[Comment 9]**

Would definitely be best to have researchers include antibody validation data as a supplemental figure in the publication so it is in context of the experimental design and intended use and is easily findable for investigators reading or trying to replicate that work.

**[Comment 10]**

I think finding a place to store, having it all in a useful / consistent format is challenging.

**[Comment 11]**

The antibody validation data is on the manufacturer's websites. In fact customers have reached out to us for validation data for journals and we have directed them to all of the data available on our website and that has been satisfactory to top-tier journals.

**[Comment 12]**

I've pushed the feasibility of this one up because it is feasible if there is sufficient funding and willingness to support this. Many other disciplines have subject specific repositories and publishers have to cope with this dynamic and evolving landscape. Making this feasible is no different therefore, than for all data sharing policies where data deposition is asked for (although publishers have a hard time navigating the landscape and enforcing deposition because of highly variable repository standards). Reporting of RRIDs and the expectation of validation data deposition could become an integral part of data sharing policies at publishers. Initially this could just include specific wording in policies on websites without checking for compliance (but even having appropriate wording is a big step for most publishers). This doesn't guarantee enforcement but could help raise awareness among researchers and capitalise on the increasing interest of funders in the data sharing policies of publishers. Monitoring RRIDs in articles to look at the effectiveness of publisher policies in relation to antibodies could be done independently to showcase good practice. RRIDs could also potentially be included in the metadata that publishers submit to Crossref in the same way they submit ORCID data etc. This would also help track which papers use antibodies and enable "cited-by" for antibodies (addressing the incentive problem). Because DataCite also picks up data from Crossref, DataCite could link RRIDs to datasets containing validation data etc and enable bidirectional discovery. This could potentially be a lower-cost way for publishers to embed RRIDs into their workflows and to raise the expectation that validation data must be deposited.

**[Comment 13]**

I agree with other comments about the need for a clear definition of "validation data". Additionally, this would require trained data checkers (who would that be? which stakeholder would be responsible for providing the quality check? how often would that data be checked/reviewed/updated?). Although this could be useful for the community, it would be quite difficult and burdensome to implement.

**[Comment 14]**

I am confused as to how you want me to "re-rate' this question. The responses from reviewers are valid. The suggestion " in/with the publication or on manufacturers websites" is excellent, however, hard to enforce.

**[Comment 15]**

I think requring authors to deposit in the manuscript makes sense, perhaps alongside a requriment to deposit in a third party site if its not in the manuscript. I feel that depositing in manufactorers websites would be less powerful as they are not independent and may not be maintained following mergers etc.

**Item R5**

*Transparency Badges or Tiered Scoring. Journals should introduce a tiered scoring system or transparency badge to recognise papers that meet higher standards of antibody reporting and validation.*

**Participant Comments**

**[Comment 1]**

To make this more feasible, we need a system of tiered scoring that can be easily adopted by all stakeholders.

**[Comment 2]**

I still do not believe that this will be enough of a motivation for people unfortunately. The stakes need to be higher - e.g. article cannot be published without minimum validation data

**[Comment 3]**

If a society driven set of guidelines are established, I can see this being something that could eventually being implemented by journals. Journals setting the standard is an interesting point, as this would need a culture shift in researchers too the approach to routinely validating antibodies. I think it would be unfeasible for journals to suddenly start requiring all antibodies to be fully validated. Some form of bade or checklist, supplied with compliant manuscripts is used in other fields.

**[Comment 4]**

The barrier to this is the lack of incentives in the research community to adopt open science more broadly, progress is slow and PLOS is doing great work to improve this, but more is needed from funders and institutions to include open science in their research assessment frameworks.

**[Comment 5]**

Yes, I agree with the comments from other 1st round users. Standardized reporting and administration would be tricky, and I think badging implies standards, which as noted have yet to be administered. The need for the impartial 3rd party watchdog/organization emerges again.

**[Comment 6]**

I agree with previous comments that this badge system would be difficult to implement, likely not standardized, and not very informative. There is also no incentive for having the badge on a publication and therefore would not change behavior of investigators using the antibodies. If a paper has the badge, someone trying to use that antibody might misinterpret the signal (eg antibody was KO validated in WB but another researchers assumes it was also KO validated in IHC).

**[Comment 7]**

We see this as an additional piece of data, but hard to manage and ensure consistent.

**[Comment 8]**

The journal should set the standard for all manuscripts accepted; there is no need for an additional badge.

**[Comment 9]**

on reflection, I don't think journal-scored badges will work in the end because journals per se are not incentivised to maintain this and there will be inconsistencies across journals and publishers. Any system of scoring or checking needs to be done independently of the journal or publisher.

**[Comment 10]**

I would still maintain that this would not be easily adoptable by different journals because developing and agreeing on standard criteria would be challenging.

**[Comment 11]**

all of the reviewers' responses are valid.

**Item R6**

*Reviewer and Editor Training in Antibody Validation. Publishers should provide training resources for reviewers and editors to help them evaluate antibody validation information in submitted manuscripts.*

**Participant Comments**

**[Comment 1]**

This plan adds to the burden of reviewers, who will have a broad and varied interpretation of what constitutes adequate antibody validation. I believe it should be the responsibility of the journal to have an editor assess the quality of antibody validation, thereby establishing a clear and consistent standard for the journal.

**[Comment 2]**

If there's a central resource provided by Only Good Antibodies that would work as a Gold standard which stakeholders can abide by.

**[Comment 3]**

It is good to provide information and resources - However, it's possible that despite providing the resources, the main reason for NOT validating antibodies may not be a lack of awareness, more an avoidance of something seen as mundane and detracting from "doing science"

**[Comment 4]**

If these are a community driven consensus, and are a freely accessible resource, as suggested, this could be effective.

**[Comment 5]**

My scoring on feasibility remains uncertain because even with available resources (thanks OGA for creating these) ensuring that editors/reviewers take the time to learn from these is challenging. Having said this, I do think it is worthwhile to have the resources available and I am sure that most journals would be open to including these on their websites and/or promoting them amongst their editorial board.

**[Comment 6]**

I think freely available resources from Only Good Antibodies would be more effective than individual publisher resources, minimising variation across publishers. Feasibility largely relies on reviewer interest and proactive engagement with reviewer resources, which can be low. Improving accessibility to resources (eg linking to these in reviewer invitation emails for antibody validation articles) would help, but would be difficult to implement on the publisher side.

**[Comment 7]**

Publishers may really not feel that they should have this role. But it would certainly be helpful, so that everybody knew how such an assessment should be made... although I do wonder if now an AI tool might be generated to help implement this... after all we generally agree on the necessary information needed.

**[Comment 8]**

It depends on what the training resource is, how influential it is, and whether it can be standardized for many validation situations

**[Comment 9]**

Having training videos would be relatively easy to implement if publishers simply provided links to an external training resource in the reviewer materials. But i do not believe reviewers would actually watch the video or participate in the training. They aren't getting paid to review for journals - what is the incentive for the extra work?

**[Comment 10]**

The journal should set the standard for all manuscripts accepted and reviewers should be aware of those standards.

**[Comment 11]**

if the resources are provided from an independent and credible resource (such as OGA) then I think such training would become both more effective and feasible although it would remain limited from the publishing side. I think that publishers will most likely link to the external resources without providing additional training (in the same way they link to resources around consort etc ). Genuine effectiveness will rely on editors and reviewers reading the material and my hunch is that editors and reviewers in larger journals with distributed editorial boards (like megajournals) are less likely to read the material.

**[Comment 12]**

Re-reading the question, this could actually add to reviewers and editors workload. Who will ensure that they take the resources' content into consideration when reviewing manuscripts?

**Item R7**

*Automated Tools to Flag Validation Concerns. Publishers and other stakeholders should support the development and use of automated tools to flag potential antibody validation issues in submitted manuscripts.*

**Participant Comments**

**[Comment 1]**

I believe that a package like SciScore would be great for this and could be updated often enough to maintain relevance and accuracy. The additional costs could be a factor and ideally would be absorbed by the journal and not passed on to authors. A group of invested people could potentially put together a program comparable to SciScore, perhaps using funding from various sources, to enable this approach.

**[Comment 2]**

RRIDs already exist and are kept up to date by vendors submitting their data regularly. RRIDs link to papers and can flag bad reagents pre publication or post. Sciscore also exists and works quite well. The cost to publishers is on the order of a couple of dollars per manuscript. The will to improve manuscripts is the thing that is missing for many publishers.

**[Comment 3]**

The main barrier to the feasibility, is there's a learning curve in use of a tool and likely cost attached.

**[Comment 4]**

Would need peer reviewed validation data, with a community agreed standard to define what a poorly performing antibody is.

**[Comment 5]**

SciScore is currently a limited tool and can often rate less standard article types poorly as it applies the same checks indiscriminately. I agree that internal human checks would still be necessary and might be unfeasible for publishers.

**[Comment 6]**

I think the point of linking the "bad antibody" information back into the existing literature is very important. I also think that RRID has made one of the very few solid efforts that help. It would be great if the providers, especially the majors, decided to implement this systematically. The fog would clear a bit. The key word is "support" which means money and investment.

Maybe an AI could give an "antibody validation rating" for manuscripts? It wouldn't be too difficult. That would be an unpopular task for OGA to set up!... So 10 for "all antibodies used in this study credibly validated by supplier and user as F2P" to zero for " no evidence that either supplier or user has succeeded (sic) in showing that any of the antibodies used bind their postulated targets in the experimental contexts described".

**[Comment 7]**

I think antibody validation can be nuanced so the system would need to be very sophisticated to prevent the honest mistakes (ie not data fraud or image manipulation).

**[Comment 8]**

Criteria for what would initiate the withdrawal would need to be clear.

**[Comment 9]**

When an antibody is removed from the market, it is not disclosed as to why it was removed. (Poor) performance issues is only one possible factor. Low(er) sales, polyclonal to superior monoclonal switch, creating a better clone with more applications for a particular target - these are also valid and not negative reasons to discontinue a particular antibody product.

**[Comment 10]**

I think that this development is becoming increasingly feasible for the really big publishers who are already developing automated tools. As AI develops this is likely to become much easier and more cost effective. However, it is likely to be limited to the big publishers because smaller/scholar publishers are unlikely to have the capital to develop these tools independently (unless they partner with the big publishers). Like much of the recommendations listed in this exercise, there will be a split between what the large publishers are able to do and the smaller ones, who are more resource constrained. And for the larger publishers, the more it is in their own interests to do it, the more likely it is they will implement it. For the large publishers therefore I think it comes down to making a business case for them (unfortunately). I think there is a case to be made centered on improving the rigour and quality of articles that are submitted to them (and which are therefore more likely to convert into publications...)

**[Comment 11]**

Ai is improving rapidly so it might be more feasible in the future

**Item R8**

*As part of the submission process, authors should describe how antibody specificity was assessed, making reference to the International Working Group on Antibody Validation (IWGAV) framework. A brief paragraph, checklist, or table should indicate which of the IWGAV pillars were used, if any (e.g., genetic, orthogonal, independent antibody, tagged protein, IP-MS).*

**Participant Comments**

**[Comment 1]**

It might be worth considering a proposal in which journals be encouraged/required to include this issue on their website in the list of items needed to submit a manuscript, so that authors are not learning about this late in the process. Included in this requirement would be a link to the paper describing IWGAV issues so that authors can know what they are heading into.

**[Comment 2]**

There may be a barrier for adoption for less well funded institutions or for labs in the Global South without access to suitable local testing facilities.

**[Comment 3]**

If clear, community driven guidelines can be established, then journals could request compliance, or in the first instance this could be presented as an optional checklist/statement. In terms of there being a barrier as labs are not equipped, as more publications include validation, the required information may be out there in the literature. Otherwise, this should be a concern for reviewers at peer review (as note in above query relating the providing guidance for reviewers).

**[Comment 4]**

Well it's feasible to ask, but I agree with the criticism that "at submission" is too late, and if authors aren't equipped to complete validations to show the antibodies they are using are in fact workng correctly, then maybe they should be submitting a manuscript using them? I think that is the problem OGA and the rest of us are so concerned about.

**[Comment 5]**

More education/training may be required for people to understand more about antibody validation and the working group recommendations.

**[Comment 6]**

Having the checklist as part of the submission process would be feasible (similar checklists are already required for use of animals), but at the point of publication the data is already gathered so I do not see researchers going back and validating antibodies post-hoc as it would expend more time, money, and risk uncovering an answer that you don't like.

**[Comment 7]**

this would require an additional question and field in any submission system (scholar ONE, editorial manager etc., although note that Wiley has now developed its own bespoke platform called Research Exchange). Is the expectation that journal staff/vendors will then check the answers or that the information will be checked by editors and reviewers? For specialty journals that receive a lot of papers using antibodies, it does make sense to have the extra field. For large multidisciplinary fields it becomes more problematic to add yet another field on a specific discipline although doable (e.g. drop down about wheter the article uses antibodies etc). Policy wording on journal websites would also need to be changed to include information about IWAG and an acknowledgement that not all labs have the ability to do this. Reviewers and Editors would also need training.

**Item R9**

*Where antibodies are critical to key experimental conclusions, where technically feasible, authors should either: present validation data based on a genetic strategy (e.g., knockout or knockdown), or cite previously published genetic validation data.*

**Participant Comments**

**[Comment 1]**

Journals can mandate this, but ultimately needs clear guidelines, from the community, to editors and reviewers as the what is required, and in what contexts.

**[Comment 2]**

If the approach is not technically feasible in a particular system (or financially accessible) then the authors should specifically state that in M&M.

**[Comment 3]**

Not everyone has access to knockouts or KO generation capabilities, and that doesn't necessarily mean that their antibody lacks validation. There are ways to validate without KOs.

**[Comment 4]**

Unclear if this would change behavior. And difficulties in obtaining KO systems or running these experiments for antibodies to post-translationally modified targets would be difficult.

**[Comment 5]**

There are targets where this is not an option

**[Comment 6]**

Again, it depends on the target protein. It is also important to note there are multiple hallmarks of validation; no one validation method is better than another, but the cumulative effect of several validation methods that is most powerful.

**[Comment 7]**

Clarifying in the wording that feasibility depends on whether the lab has the technical ability makes it implicit (to me) that there is an opt-out if you can't do it which makes this a more reasonable ask and therefore potentially more feasible for journals to implement. But the effectiveness then seems ambiguous/diluted to me. Would the author then provide the reason why they e.g. cannot do it (if they can't). And if they can't provide the validation would the assumption be that the paper is rejected before peer review. It's not clear what the expectation is for the journal decision given there is now a clause that allows you to essentially tick the box even if you haven't validated the essential experiments. I guess I don't see the point of including a request for genetic validation in the first place. I prefer the wording of the original - it's then up to the authors to explain to the editor why they don't have this and for the editors/reviewers to assess whether the explanation is reasonable.

**Item R16**

*Encouraging Manufacturer Transparency. Funders should directly engage with manufacturers to encourage them to publish antibody validation datasets and adopt consistent metadata standards (e.g., RRID, clone ID, lot number).*

**Participant Comments**

**[Comment 1]**

While funding agencies have power, I believe the responsibility lies overwhelmingly with the researchers and the journals who publish the work. I also have some doubts as to the ability of funding agencies in one country to make demands on a company in another country.

**[Comment 2]**

It may be difficult to get Funders to do this work, since some funders don't just have interests in science but philanthropy in general.

**[Comment 3]**

It's true that this could seem out of scope for funders. However, they do have a significant influence so it would be good to see how this can be leveraged to drive best practice

**[Comment 4]**

Having read the comments above, especially the one around why Ab manufacturers would be influenced by the funders, I realise that this approach may not be so effective after all. Maybe funders should focus on the funding transparent and reproducible science. This in turn will create the incentive for manufacturers to change their practice if demand for their products change.

**[Comment 5]**

... in answer to the query "why would manufacturer be influenced by the funder" and especially thinking about the situation in the US - it seems to me that the sooner the manufacturers get on board with this the better. As I originally wrote, if the funders (or their bosses) realize that the manufacturers are selling poorly made and documented products, then it is only a matter of time before they decamp. It is in the interests of everyone, except the cowboys, to insist that users of (especially) public funds, i.e. the researchers, and clinical laboratories suffering from poorly validated antibodies, only use antibodies that are professionally validated. Otherwise funds are wasted and should be withdrawn. It is a tricky rope to walk, because once again, the issue is who decides who is selling a " properly validated antibody". As the man said, " we know it when we see it", but a set of standard requirements, agreed by all is hard to define. At the minimum, it really can't be too hard to implement publication of consistent metadata.

**[Comment 6]**

I agree with the concerns listed above. This is outside the scope of funders, the vast majority of funders don't have a relationship with manufacturers, manufacturers don't have an incentive to follow through on funder requests, and many funders are government-affiliated which would not allow this.

**[Comment 7]**

In principle, I think it is feasible for funders to encourage manufacturers to do this e.g. in public statements, perhaps in collaboration with other funders (similar to cOAlition S statements about publishing). I don't know if manufacturers would listen however given that publishing their validation data might put them at a commercial or competitive disadvantage (in their view).

(As an aside, I wonder if there is a role for publishers to independently incentivize manufacturers - perhaps providing a dedicated data journal for antibody validation studies or giving priority in peer review for papers using antibodies with published validation data).

**[Comment 8]**

It would be challenging for funders to directly engage with manufacturers, and even if they managed to, what leverage would they have? Encouraging the adoption of consistent metadata standards could be welcome, but it would probably have to come from some sort of regulatory body instead of funders.

**Item R17**

*Funders should collaborate with independent benchmarking initiatives such as YCharOS or equivalent open-science consortia. They should directly co-fund the validation of antibodies against key targets to create suites of validated tools that can be used by researchers to improve data quality and thereby increase return on investment for the funder.*

**Participant Comments**

**[Comment 1]**

Feasible, but where is the money coming from to do the job?

**[Comment 2]**

Whist this is an extremely important issue, many funders (in particular single disease charities) may struggle to justify spend on what may be considered more of a general infrastructure issue and less specific to their charitable aims.

**[Comment 3]**

While impactful, this type of funding falls outside the scope of what general foundations/funders support. Charity foundations operate on donations from patients or loved ones, and using precious funds on antibody validation is to far removed from improved treatments and cures to be enticing for donors. Government funders also don't have a mechanism for this support. And most funders operate through open funding callouts which would require the right funding opportunity for YCharOS and for YCharOS to apply to the grant mechanism (and successfully align with the callout and pass review).

**[Comment 4]**

These associations need accountability to researchers and manufacturers

**[Comment 5]**

I still remain unsure about the feasibility of this given funding constraints but if there are the funds available and a long term commitment by funders then it seems very feasible.

**Item R18**

*Funders should promote industry participation in independent benchmarking initiatives by asking applicants to consider working with initiatives such as YCharOS or equivalent open-science consortia as part of their work. This could be achieved either by including the recommendation in the guidance notes or flagging to applicants at the panel review stage.*

**Participant Comments**

**[Comment 1]**

Just including such wording would help raise awareness of the issue and in the current wording does not add to the burden.

**[Comment 2]**

Although this is a good idea, I strongly question whether funders have significant enough leverage over industry to make this a reality.

**[Comment 3]**

Well, it all comes down to things like OGA/YCharOS, doesn't it? And there are a lot of antibodies to test. If you recall, the CD antibody testing was a community effort established because the research community became so frustrated with the "inadequate antibodies" available to define the cell populations. I don't know enough of the history to say more at the moment, but that model worked and is working well.

**[Comment 4]**

Sure, funders can suggest that applicants work with YCharos. But if no additional budget is provided for this work and the applicant must divert some of their funds to support the YCharOS work, then it is not going to happen. If the funder mandates the applicant work with YCharOS and pays for the antibody characterization, then the timelines for data return are too long and the project will be significantly delayed.

**[Comment 5]**

clarifying the wording helps me to see that this could be an effective route and is more potentially feasible

**[Comment 6]**

I think this point could be clarified, does industry refer to pharma companies or the antibody manufactrers.

**Item R21**

*Universities, research institutions and learned societies should recognise and support local champions or experts in antibody validation (e.g., through the Antibody Champions Scheme).*

**Participant Comments**

**[Comment 1]**

I would change the wording to be a bit more inclusive of other efforts and not limit it only to the Antibody Champions Scheme, perhaps "support efforts such as the ...". Universities with antibody/monoclonal core facilities and/or microscopy/imaging cores/centers should absolutely be encouraged to include such staff.

**[Comment 2]**

It is excellent that the UK is driving this. It would be a model for international uptake. Of course, "advice on antibody selection " is vulnerable to unscrupulous providers gaming the system. This is why I still greatly admire what CiteAb have done because it is impartially looking at usaqe as an indicator of quality.

**[Comment 3]**

There may be limited effectiveness with a voluntary scheme. As little additional inducement for the 'champions' (eg reimbursement of training time) combined with funders making institutional support for the scheme a requirement of their T&Cs, could greatly enhance its impact.

**[Comment 4]**

Would the manufacturers- the scientists who make and work with the antibodies on a daily basis be part of this scheme?

**[Comment 5]**

Considering the financial challenges facing universities at the moment, and the upcoming REF exercise for instance, it may not be easy for them to have the resources to support the scheme.

**Item R23**

*Shared Infrastructure for Validation Data and Standards. A coordinated, shared infrastructure should be developed (or expanded) to aggregate antibody validation data, provide standardised formats and tools for sharing, and enable journals, funders, and institutions to access validation status or red flags.*

**Participant Comments**

**[Comment 1]**

Costs and feasibility aside, this would be huge step towards addressing the key questions.

**[Comment 2]**

would require significant buy in from stakeholders in terms of time and cost. Concerns around the level of undertaking and ability to commit to this in the near term due to current funding climate

**[Comment 3]**

Standardization is the challenge. But, yes, a huge undertaking. For cell surface molecules on cells that can be adequately put through flow cytometry it is doable, but for the rest... very tricky.

**[Comment 4]**

This would be a huge undertaking to staff, QC, and keep updated.

**[Comment 5]**

I've pushed the feasibility of this up to 7 because I think in part it is a matter of enough people wanting this to get it off the ground. There are really good examples of community governance around activities (including the Barcelona Declaration or OpenCitations, which both started out as volunteer based and then attracted support from funders and institutions). I think it is really important that the creation of this be at least attempted and think its feasibility is higher because there are good examples that are working that could act as a model for the approach.

**[Comment 6]**

The panel made some pertinent comments regarding funding and governance requirements. I have to revise my rating.

**[Comment 7]**

I think at the moment this proposal is a bit too vague to score.

**Item A1**

*Journals should have a clearly stated standard for antibody validation and reporting.*

**Participant Comments**

**[Comment 1]**

Due to the many and varied uses of antibodies in research the wording by the journal would have to be inclusive. This is also why I believe the journals should have editors with extensive knowledge and experience in this arena to be the final judge of whether or not validation standards are being met.

**[Comment 2]**

Might be challenging to get a uniform standard across journals, and have a coordinated implementation. Might be difficult to decide on minimum standards?

**[Comment 3]**

There would need to be a consensus from the community. If this is in place, then it is more feasible for journals to have clear standards.

**[Comment 4]**

This approach would only be effective if there were community standards already in place or consensus across most journals, hence my 6 on both.

**[Comment 5]**

This would be more feasible and effective if there was external standards guidance that could be linked to, rather than individual journal policy.

**[Comment 6]**

If YCharOS / OGA acted as an interface to set up an agreed standard reporting template, and aligned it with the major publishers it might help.

**[Comment 7]**

Again, a checklist may not be feasible depending on the specific target(s). It is also important to note there are multiple hallmarks of validation; no one validation method is better than another, but the cumulative effect of several validation methods that is most powerful.

**[Comment 8]**

If the standard was a requirement then it would be effective, however it would have to be agreed across journals or there is a risk that authors would publish elsewhere.

It is feasible for journals to create standards, however the feasibility of these being adhered to in the short term (as work will have already been completed) is unclear. This would require a long lead time to make researchers aware of standards prior to starting work.

**[Comment 9]**

Having a clearly stated standard even if the policy is not fully aligned with the gold standard is a start and a means for journals to signal they are on the right trajectory. There could be a tiered approach for journals to follow (a bit like there has been for data sharing policies - from encourages to expects or mandates etc) that could help provide a transitional path for journals to improve reporting practices.

**[Comment 10]**

Unclear if same standards for all journals of if would differ

**[Comment 11]**

If different journals have different stated standards, then this would complexify submission processes. This could also be challenging because these standars would have to be developed and agreed across fields. **[Comment 12]**

A set of external standards would make this more feasible to implement

**Item A2**

*Detailed protocols and validation data should be included in the manuscript or supplemental data.*

**Participant Comments**

**[Comment 1]**

This should be true only if such data is not already available in publications or manufacturer's websites (with sufficient details to assess)

**[Comment 2]**

Journals would need to police this practice and most will not do so, but even some journals will help the whole

**[Comment 3]**

One would like to think that researchers have carried out the relevant validations, and if they have the data, they should be able to share it in a submission. The main issue I see is compliance from researchers.

**[Comment 4]**

Not sure that these would be effective just yet, before more work is done upfront with changing the research culture. Detailed protocols should be shared on protocols.io and a link to this added to the paper methods section. Validation data would better exist in one repository or few repositories dedicated to this.

**[Comment 5]**

Depending on length, placing this in supplemental data or repository would be more feasible than in the main manuscript.

**[Comment 6]**

Well, either the users have characterized the reagents they claim to produce the results they are showing, or they haven't so I think this approach would cut out a lot a sloppy science. The devil lies in that nice word "detailed". If a set of standard validation protocols were to be made available, i.e from OGA, and variation from the protocols were to be noted, then maybe it would work.

**[Comment 7]**

Would be amazing if authors complied. But think this coming at the submission phase would reduce compliance.

**[Comment 8]**

Providing the format and locations to store could be challenging

**[Comment 9]**

Protocols are feasible, validation data may not be available.

**[Comment 10]**

Supplemental data is not machine readable and so is less effective

**[Comment 11]**

This might be effective if there was enforcement, curation and review of the data and if the data are machine readable in the main body of the manuscript. Putting data like this in the supplementary material essentially makes it invisible and undiscoverable by machines and I would strongly urge that at least this option is removed. Overall I cannot support this recommendation as I don't think publishers should be storing data inside manuscripts. Not least, these data ought to be in a publicly available repository and articles should be linking to that data via an RRID. Reporting that the validation has been done with links to the appropriate resources should be in the manuscript. The actual data needs to be maintained, checked and curated in an appropriate repository. Protocols could be published separately by journals but there are also other platforms that could publish these.

**[Comment 12]**

It's probably feasible, but this could represent more work for authors. The level of detail required would have to be defined and agreed amongst journals. **[Comment 13]**

Adding validation data to supplememtary material would mean that it is not machine-readable, and therefore not easily discoverable

**[Comment 14]**

data should be stored in a publically available repository and articles should link to the data via RRID [Disagreement between experts on most appropriate location for data - in MS/data repostory/supplementary data?]

**Item A3**

*Journals should appoint a specialist editor that focuses on reproducibility.*

**Participant Comments**

**[Comment 1]**

Many journals might lack the resources to recruit a specialist editors for this purpose.

**[Comment 2]**

Not sure that journals, in the current climate, have the capacity to do this

**[Comment 3]**

Ideally, resources would be available to all editors on the requirements for reproducibility within their area, and more experienced editors would be able to share resources with less experienced, or provide advice. Appointing a specialist editor would not be feasible. Also, it is not clear how this would work - a large number of manuscripts involve antibody work so would the reproducibility editor screen them all? What they directly handle any?

**[Comment 4]**

Interesting idea, it would need consensus from various journals on what the role would involve so that standards are consistent to authors.

**[Comment 5]**

Feasibility depends on the scope of the role - an Editor checking every manuscript for reproducibility issues would likely be unfeasible, however checking escalated papers might work, or writing internal guidance/editorials for resources. A standalone reproducibility role might also be unfeasible depending on Editorial Board capacity/Editor remuneration policy, however the responsibility could be split across a few Editors, and it might be an additional, more specialised role, that some Editors would be interested to have.

**[Comment 6]**

Money is in the way.

**[Comment 7]**

How would you prioritize what manuscripts this editor is assigned or reviews?

**[Comment 8]**

I like this idea and think it is feasible. Journals/publishers could also appoint reproducibility editors across a suite of journals (or for a particular discipline) for efficiency. In some ways this would be akin to the dedicated research integrity specialists that publishers (and journals) are increasingly recruiting. A knock on consideration for feasibility is that if the new role is effective, it may create cases that need to be escalated to different journal staff. The role and responsibilities would need to be clearly laid out and be appropriate (they couldn't for example check every paper being submitted or accepted) and what would happen where articles failed etc. Such a specialist could also help train/raise awareness among other Editors and reviewers.

**[Comment 9]**

It would probably not the best approach to have just one person checking for validation data details. Additionally, what would be the baseline for the editor to agree that one manuscript contains reproducible experiments? **[Comment 10]**

Publishers would not have the resources for this

**[Comment 11]**

What standards would they be judging the MS by

**Item A4**

*Funders should maintain and provide researchers with a list of manufacturers who validate their antibodies.*

**Participant Comments**

**[Comment 1]**

I wish such a list could be generated but I don't think it can and I don't think the funders have the bandwidth to enforce its use.

**[Comment 2]**

Its true that some manufacturers make more effort than others, it does not mean that an individual antibody is more reliable. Therefore, it might give a sense of false confidence in a particular product, and may negatively impact the investment in direct validation carried out.

**[Comment 3]**

It would be useful to know if an antibody has been validated. If there are a set of standard validation assays agreed on by the community then it would be useful if manufacturers performed them (although I don't know if this is feasible). It is not clear to me if the funder is the person to maintain this list.

**[Comment 4]**

Not sure funders would have access to this.

**[Comment 5]**

1) "...who appropriately / correctly/ adequately / acceptably / rigorously validate their antibodies " perhaps? Select the least toxic term for emphasis.

2) the funders would need to feed back to OGA/YCharOS or other accepted impartial sources for support, because also the funders are in new territory here.

**[Comment 6]**

Even manufacturers that validate antibodies might not have KO validation on all of their products. Researchers might misinterpret this list and have too great of a sense of security in the products and use them in applications that haven't been validated. And this is well outside the scope of a funder's role.

**[Comment 7]**

Hard to define what is included

**[Comment 8]**

Who will create the list?

Validation is not the primary concern for researchers when selecting an antibody.

**[Comment 9]**

I don't think this is a role for funders. Having such a 'white list' might be effective if it came under the responsibility of the community/discipline e.g. under the remit of the community infrastructure (as part of OGA or YCharOS). Funders are too variable (esp at a global level) and I don't think they would keep it maintained and updated. It needs to be done by an independent body that is not controlled by govt or the private sector)

**[Comment 10]**

Need to define who sets the standards

**[Comment 11]**

This could be effective in theory, but considering all the other elements funders have to worry about, I'm not sure it would be realistic to ask them to do this. This also poses the challenge of keeping the list updated.

**[Comment 12]**

NIH will never do this **[Comment 13]**

Not a role for funders

**[Comment 14]**

Who would maintain it

**[Comment 15]**

Who would set the standards

**[Comment 16]**

Could give a false sense of security - what is an appropriate antibody/protocol for one application may not be appropriate for another

**Item A5**

*Funders should mandate the use of recombinant antibodies (or at least monoclonals) when available. Justification should be required if a polyclonal is used.*

**Participant Comments**

**[Comment 1]**

Too many good polyclonals in use today and if a lab has been using one or a mAb for years they are not going to switch.

**[Comment 2]**

This might be target dependent.

**[Comment 3]**

It's not clear if Funders will mandate their researchers to do this.

**[Comment 4]**

Some academic labs have bespoke antibodies that they have made that are polyclonal - they may not have sufficient funds of knowhow to make monoclonals. As long as these antibodies are well validated and that data is shown, it seems reasonable that they can be used with caution. What about the cases of low abundance targets where signal:noise is an issue and monoclonals are not effective? Not many recombinant mixed monoclonals are available. The best approach is to ensure the author has tested an alternative, and flag any residual reproducibility risk.

**[Comment 5]**

I suspect that labs who have long used particular polyclonals will be reluctant to change, and will are that they work in their hands, and reliably give the expected signal.

**[Comment 6]**

This would only be impactful if funders coordinated this to some degree.

**[Comment 7]**

Not just recombinant. RRID-defined and better published-sequence recombinant.

**[Comment 8]**

This is too granular of information for a funder to review in applications or monitor in reports. Studies might be using a dozen antibodies. This could be encouraged but there is no way to mandate or enforce this.

**[Comment 9]**

There are applications which benefit from the use of polyclonals

**[Comment 10]**

Monoclonals is significantly more feasible than mandating recombinants.

**[Comment 11]**

It is feasible for funders to mandate this but mandates themselves may not shift actual practice if there are not the resources to support researchers and others to do this. The experience of mandates in the past show they are are far more effective when there is at least some infrastructure and education in place and they have more of the community behind them. At this stage, I think funders need to be enablers and it is too early to actually mandate. Once there are resources, infrastructure, training, etc in place then mandates can work. Putting a mandate in place when it is not yet possible for the mandate to be implemented is ineffective. The timing needs to be a careful balance of what's possible now as well as what's desirable.

**[Comment 12]**

Validation data for each Poly batch used should be generated and shown.

**[Comment 13]**

I think as long as there is the option to choose to use a polyclonal, but to have to justify it, then this could be a good approach that educates. A bit like requring documentation of the validation that has been carried out. **[Comment 14]**

Labs using existing polyclonals will be reluctant to change

**[Comment 15]**

this would only be impactful if funders cordinated

**[Comment 16]**

Funders haven't got the capacity to enforce this - each projects may be using many antibodies

**[Comment 17]**

some applicatinos benefit from using polyclonals

**[Comment 18]**

mandates themselves do not shift actual practice if there are not the resources and knowhow to support researchers to carry it out

**[Comment 19]**

As long as there is still the opportunity to use a polyclonal if necessary, it would be good

**Item A6**

*Funders should mandate that if funds are used to develop an antibody as part of a research project, the reagent must be a recombinant antibody and must be made available to the wider community.*

**Participant Comments**

**[Comment 1]**

I think this "rule" needs a time element. A lab that invests the time and money and effort in generating a new, rAb for use in their research should be given some length of time to use it and share it with select others, before requiring submission to a place for distribution widely. Something like one year after the publication of its first use?

**[Comment 2]**

As before, I like this idea, but I am not sure how funders have the resources to check compliance with this.

**[Comment 3]**

In theory, many journals already demand that reagents used in manuscripts published must be made available to the community. But, depending on the reagent, this may be a really big job for the researchers concerned. So one question is: would such reagent be available in practice? I favour this approach but some may not. Plus, making a recombinant antibody is still an undertaking. But, if possible, should be encouraged.

**[Comment 4]**

This would require follow up to enforce the mandate , post completion of the award, requiring resource that some charitable funders may not be able to commit to

**[Comment 5]**

Development of recombinant reagents take 12+ months, which is why academic investigators go with quicker processes (polyclonals or hybridoma). Grant timelines are not realistic to mandate recombinant antibody development. Also, to enforce the requirement to make antibodies available to the wider community the technology transfer office needs to be involved and strict contract language needs to be included. Even as a funder with 15 years of experience in this type of sharing policy, we still cannot reliably have institutions deposit their antibodies at repositories or license them to companies.

**[Comment 6]**

This seems feasible as it sets out the requirement before the work has started. Funders should also required reporting about how the recombinant will be made available.

**[Comment 7]**

May incur extra costs **[Comment 8]**

Would Funders have the resources to enfoce it

**[Comment 9]**

Grant timelines make this unrealisitc - recombinanat reagents take over a year to generate

**Item A7**

*Antibody manufacturers should be encouraged to assign RRIDs to their products at source.*

**Participant Comments**

**[Comment 1]**

There are some suppliers that will ignore this, in the same way they have ignored the calls to improve validation

**[Comment 2]**

Really, an easy one.

**[Comment 3]**

this would be fantastic.

**[Comment 4]**

Listing the antibody's full name, including clone and possibly Lot# along with the manufacturer's name is more than sufficient.

**[Comment 5]**

This should be feasible - but it requires manufacturers to be willing and I don't know whether manufacturers take heed of funders (in the same way that some funders have mandates around open access and metadata, which are then ignored by some publishers)

**Item A8**

*Manufacturers should perform a standard set of validation experiments on their reagent antibodies and make the data available.*

**Participant Comments**

**[Comment 1]**

Not going to happen. Too expensive.

**[Comment 2]**

If the researchers require this, then manufacturers will more likely make this available.

**[Comment 3]**

"Standard set" may be hard to implement. Once again, it demands an agreed set of protocols. Agreed : between manufacturers; between researchers; within the community? How to get to an acceptable consensus. And no the 5 pillars won't work, because protocols vary between labs and between researchers. But, if the protocols could be agreed upon, yes, great idea. And the manufacturers are supposed to do this anyway, aren't they?

But are you distinguishing between suppliers (OGS agreements.…) and the people who make the reagents?

**[Comment 4]**

This is what our company does. And Validation is a continual/on-going process for every antibody and every product. We have always believed in transparency. We encourage customers to ask for validation data since all of the validation data cannot be displayed on the website.

**[Comment 5]**

I don't have any sense about the feasibility of this. I would think that they have the capacity and capability to do this but for some it might incur a significant loss of revenue so there would be no incentive to make it feasible?

**[Comment 6]**

Will affect price of reagents and increase development costs.

**[Comment 7]**

Some pane; members felt this isn't feasible owing to the cost

**[Comment 8]**

Others suggested reputable manufacturers are already doing this

**Item A9**

*Manufacturers should shift production to all recombinant antibodies.*

**Participant Comments**

**[Comment 1]**

By manufacturers I assume you mean commercial distributors who don't actually manufacture most of the antibodies they sell, as I understand the market today.

**[Comment 2]**

Who would mandate this?

**[Comment 3]**

I scored this as 3 as I do not think that manufacturers will shift their production practices without changes in customer demand.

**[Comment 4]**

Do recombinants work well for small-molecule targets?

**[Comment 5]**

The resources and timing to do this would be long

**[Comment 6]**

Poorly characterized recombinant monoclonal antibodies are still available commercially. Recombinant manufacturing helps with consistency, but proper characterization by the vendors remains critical; further, laboratories using antibodies are responsible for ensuring performance in the specific assay/protocol.

**[Comment 7]**

I don't have a sense of how realistic this is? **[Comment 8]**

recombinant antibodies still require validating to ensure they bind to the target protein

**[Comment 9]**

How would this be enforced?

**Item A10**

*A learned society for antibody validation should be established.*

**Participant Comments**

**[Comment 1]**

The antibody society and journal antibodies seem like they exist. Perhaps taking over leadership?

**[Comment 2]**

Depends on the time constraints of those involved, but overall a good idea. Difficult to judge the level of impact this oculd have on wider scientific community

**[Comment 3]**

Not sure how this learned society would overlap with existing organisations trying to achieve the same thing.

**[Comment 4]**

Funding?

**[Comment 5]**

New learned societies can take a long time to be set up, get community recognition, buy-in and funding, build reputation and relationships with commercial companies, funders, publishers etc, as well as having a lot of administrative burden. A large part of their function is also in organising conferences and meetings, and sometimes having their own journal. With the functions outlined above, it may be more feasible and effective for existing initiatives like OGA and YCharOS to carry these out.

**[Comment 6]**

I do hate the badge system. But otherwise: yes good idea. Independent assessment of validation data is a big one, and may be very hard.

**[Comment 7]**

who is paying for this?

**[Comment 8]**

The more scientists that are educated on antibody validation, the better.

**[Comment 9]**

Would require sustained funding.

**[Comment 10]**

I think creating a separate body for this would be ineffective when there are organisations already that could take this on -- or collaborations of several different orgs coming together under a coordinated, shared infrastructure (Q23). I think more effort should be put into the shared (and appropriately governed) infrastructure.

**[Comment 11]**

Before establishing a new learned society for antibody validation, it would be wise to:

- use existing channels for dessimination of good practices

- bring together existing societies (with members in the life sciences) that could perform some of the functions suggested

- establish stronger links across the ecosystem to encourage dialogue - and eventually achieve standards harmonisation - among the different stakeholders

**[Comment 12]**

Existing organisations should work together instead
